# Bioorthogonal in-cell Labeling and Profiling of *N*^6^-isopentenyladenosine (i^6^A) Modified RNA

**DOI:** 10.1101/2022.07.07.496599

**Authors:** Sheng Wang, Yuanyuan Li, Hongling Zhou, Li Wang, Youfang Gan, Shusheng Zhang, Ya Ying Zheng, Jia Sheng, Rui Wang

**Affiliations:** Hubei Key Laboratory of Natural Medicinal Chemistry and Resource Evaluation, School of Pharmacy, Tongji Medical College, Huazhong University of Science and Technology, Wuhan, Hubei 430030, China; Innovation Research Institute of Traditional Chinese Medicine, Shanghai University of Traditional Chinese Medicine, Shanghai 201203, China; Department of Chemistry and The RNA Institute, University at Albany, State University of New York, Albany, NY 12222, USA; Shenzhen Huazhong University of Science and Technology Research Institute, Shenzhen, Guangdong 518057, China

## Abstract

Chemical modifications in RNAs play critical roles in structural diversification and functional regulation of many vital biochemical processes. Several hydrophobic prenyl-modifications have been discovered in a variety of RNA species since the 1990s. Prenyl groups are the feedstocks of terpene and many other biological molecules and the processes of prenylation in different macromolecules have been widely studied. We present here a new chemical biology technique to identify and label i^6^A, a prenyl-modified RNA residue, based on the unique reactivity of the prenyl group. We also found that iodine-mediated cycloaddition reactions of the prenyl group occurs in a superfast manner, and converts i^6^A from a hydrogen-bond acceptor into a donor. Based on this reactivity, we developed an iodine-mediated oxidation and reverse transcription (IMORT) method to profile cellular i^6^A residues with a single-base resolution, allowing for the transcriptome-wide detection and analysis of various i^6^A-containing RNA species.

## Introduction

RNAs are highly versatile biological macromolecules with a diverse range of cellular functions associated with gene expression, catalysis, environmental stress responses and many human diseases.^1^ Endeavors in the field of RNA research have led to the discovery of many new types of RNAs and revealed their additional functions and potential therapeutic applications,^2^ including modulating signaling pathways by circular RNAs,^3^ RNA epitranscriptomics,^4^ and RNA nanobiology,^5^ etc. Advances in technology and the desire to elucidate the high complexity of RNA biology have accelerated the demands for constructing new molecular tools to further understand RNA structures and functions in cellular context.^6–7^ In RNA epitranscriptome, more than 170 naturally occurring chemical modifications have been identified to decorate and diversify various RNAs.^8^ Among those chemical decorations, the hydrophobic prenyl groups (mainly in i^6^A analogs introduced by the Mia family enzyme, **Fig. 1A** and **1B**) have been discovered in the early 1990s. ^9–11^ The prenyl groups including terpenes compounds have been found as fundamental feedstock in a wide range of biological processes that play key roles in metabolism and human diseases. However, elucidating the biological significance of these prenyl-modified RNAs remains very challenging due to the lack of genome wide detection tools. ^11^ It was recently demonstrated that the prenyl modification of tRNA affects plant root growth.^11^ Liu and coworkers reported the identity of geranyl modification in bacterial tRNA, as a polyprenyl group, specifically on 2-thiouridine. ^12–15^ This geranyl group has been shown to play crucial roles in base pairing discrimination, preventing frameshifting error, and maintaining translational efficiency and fidelity.

**Figure 1.**
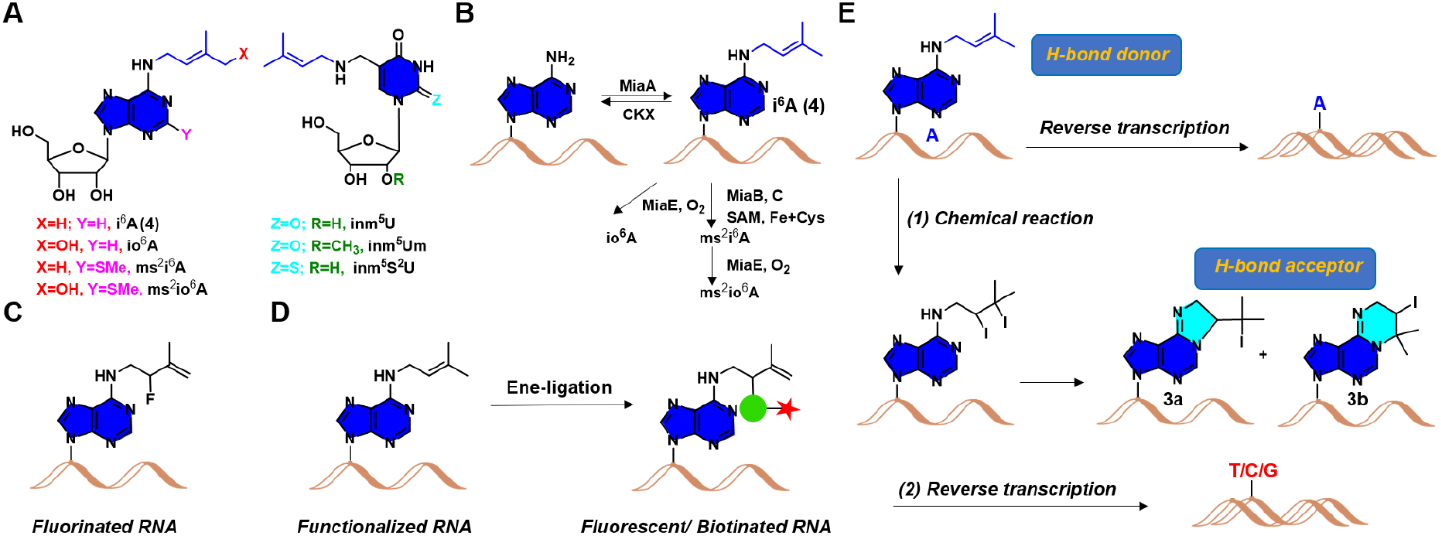
Overview of prenylated RNA residues as well as the labeling and profiling strategy described in this work. (A) Structures of prenylated nucleosides. (B) Plausible biogenetic pathways for prenylated RNAs. (C) Fluorination of i^6^A-incorporated RNA. (D) Labeling and enrichment methods proposed in this work. (E) iodine-mediated oxidation and reverse transcription (IMORT) method to profile i^6^A residues in cell.

Antigen-antibody specificity is one of the main methodologies to identify and profile RNA modifications.^16^ In addition, chemical pulldown methods have been extensively utilized in recent years to study modified residues in RNAs such as 5hmC ^17–18^, 5fc^19^, and pseudouridine^20^, etc. In contrast to using antibodies to recognize specific epitope of antigen, chemical pulldown strategies target defined functionalities on RNAs and employ highly selective chemical probes to tag the RNAs containing these modifications within the complex cell environment. We have previously developed an Ene-ligation strategy to investigate the chemical properties of the prenyl group under biologically mild conditions.^21–22^ This chemical biology method has great potential in decoding the importance of prenylated RNA residues with the emphasis on the distribution of these modifications in both healthy and diseased cells. This method can also be expanded in investigating the transcriptomic-wide profiling of these lipid-like modifications. Herein, we report a mild 2,3-rearrangement Ene-ligation based strategy (**Fig. 1C** and **1D**) for the fluorescence labeling and detection of prenyl modified RNAs via a σ-rearrangement mechanism.^23–26^ We also investigated the chemical transformation of the prenyl group in reacting with iodine and utilized this iodine-mediated oxidation and reverse transcription (IMORT) approach to profile the i^6^A residues in cellular RNA with a single-base resolution (**Fig. 1E**).

## Material and Methods

### General information

All chemicals were purchased from Sigma-Aldrich, Acros, Inno-chem, Macklin Inc, Energy Chemical and were used without further purification. Extra dry solvents, such as 1,4-dioxane, DMF, and THF, were obtained from Inno-chem in sealed bottles over 3 or 4 Å molecular sieves and stored under dried nitrogen. Organic solvents were obtained from Sinopharm Chemical Reagent Co., Ltd (Shanghai, China) and used in reactions, column chromatography and recrystallizations. Milli-Q ultrapure water (resistivity, 18 mΩ) purified through Millipore Milli-Q Advantage A1 purification system was used for all bioconjugation reactions. The reactions were monitored by thin-layer chromatography (TLC) analysis using silica gel (60-Å pore size, F254, Yantai Chemical Industry Research Institute) plates. Compounds were visualized by UV irradiation (λ = 254 nm) and/or spraying TLC stain such as a KMnO4 solution followed by electronic heating. Flash chromatography columns were performed on silica gel (60-Å pore size, 230-400 mesh). HPLC purification were performed using EasyChrom-1000 system with NU3000 serials UV/Vis. detector (Hanbon Sci. & Tech., Jiangsu, China) using an Ultimate^®^ XB-C_18_ column, 21.2 × 250 mm 5 micron (Welch Materials Inc., Shanghai, China). Separation was achieved by gradient elution from 5% to 70% acetonitrile in water (constant 0.1% formic acid) over 20 min, isocratic elution with 70% acetonitrile from 20 to 25 min, and returned to initial conditions and equilibrated for 5 min. The LC chromatograms were recorded by monitoring absorption at 254 nm and 220 nm. ^1^H and ^13^C NMR spectra were recorded at room temperature on a Bruker spectrometer (AM-600 or AM-400) operating at 600/400 and 150/100 MHz, respectively. Chemical shifts are given in parts per million, and ^1^H and ^13^C {^1^H} NMR spectra were referenced using the solvent signal as an internal standard. The following abbreviations are used for the proton spectra multiplicities: s: singlet, d: doublet, t: triplet, m: multiplet, br: broad. Coupling constants (*J*) are reported in Hertz (Hz). HRMS (TOF) were obtained from the Bruker Micro TOF ∥ Spectrometer using Electro Spray Ionization (ESI). Advion MS (ESI) were obtained from the Expression L (Beijing Bohui Innovation Biotechnology Co., *Ltd*). LC-MS analysis was obtained from the Bruker Orbitrap LC/MS (Q Exactive™) at Huazhong University of Science and Technology Analytical and Testing Center. UV–visible absorbance measurements were performed UV–visible spectrometer (Lambda365, PerkinElmer, German). Confocal Imaging was performed using a LSM 780 confocal microscope (Zeiss) with a 20× objective at 16-bit depth under non-saturating conditions. EGFP were imaged with a 480 nm (excitation) and a 510 nm (emission) and false-colored green.

### Materials

Streptococcus pyogenes (product # M0646), T7 Endonuclease I (product # M0302) Ribonucleotide solution mix (NTPs) and deoxy-ribonucleoside triphosphates (dNTPs) were purchased from New England Biolabs (USA). Transcript Aid T7 High Yield Transcription kit (product # K0441) and Glycogen (product # R0561) were purchased from Thermo Fisher Scientific. Pyrobest™ DNA Polymerase and PrimeSTAR HS DNA Polymerase were purchased from TaKaRa Shuzo Co. Ltd. (Tokyo, Japan). DNA Clean & Concentrator™-5 kit (product # D4014) was purchased from Zymo Research Corp. The DNeasy Blood & Tissue Kit was purchased from QIAGEN. The oligonucleotides at HPLC purity were obtained from TaKaRa company (Dalian, China). The nucleic acid stains Super GelRed (NO.: S-2001) was bought from US Everbright Inc. (Suzhou, China). Thiazolyl Blue Tetrazolium Bromide (MTT, CAS # 298-93-1) were purchased from Sigma-Aldrich Inc. (Shanghai, China). DPBS (CAS # 63995-75-5) was purchased from TCI (Shanghai) Development Co., *Ltd*.

### Polymerase chain reaction protocol

The concentration of DNA or RNA was quantified by NanoDrop 2000c (Thermo Scientific, USA). Polymerase Chain Reaction was conducted using a A300 Fast Gradient Thermal Cycler (Long Gene Scientific Instruments, Hangzhou, China). Briefly, each reaction was performed under conditions of initial denaturation at 94 °C for 3 min, followed by 30 cycles of denaturation (94 °C for 30 seconds), annealing (63 °C for 30 seconds) and extension (72 °C for 60 seconds) and a final extension at 72 °C for 10 minutes.

### Electrophoresis protocol

Agarose were purchased from Biofroxx (German). 50X Tris-Acetate EDTA (TAE) buffer (40 mM Tris-acetate and 1mM EDTA, pH 8.0) were purchased from Biosharp (China). 8 μL of each amplification product was separated by gel electrophoresis on 1 % agarose. The electrophoresis conditions were as follows: power supply (DYY-6C power supply, Liuyi Biotechnology) was set to 130 V. 1% agarose gels were run at room temperature (25 °C) for 20 minutes and stained with or without GoodView™ (SBS Genentech, Beijing, China). A DNA size marker (1 kb DNA Ladder, TsingKe Biotech, TSJ102, Beijing, China) was used. Gels were analyzed by in-gel fluorescence measurements on a FluorChem^®^ FC3 imager (Alpha Innotech). Fluorescence was measured with a blue light excitation wavelength (475 nm) and a green filter emission (537 nm) and/or a red-light excitation wavelength (632 nm) and a far-red filter emission (710 nm).

### Functional probes synthesis

The green fluorescent probe PTAD-DBCO-FITC (**8**).

A solution of compound **PTAD-N_3_** (10.1 mg, 0.04 mmol) and DBCO-FITC (5.0 mg, 0.01 mmol) in dry DMF (1.0 mL) under nitrogen was stirred at room temperature overnight. The reaction mixture was purified by RP-HPLC (Separation was achieved using an Ultimate XB-C_18_ column, 21.2 × 250 mm 5 micron (Welch Materials Inc., Shanghai, China) by gradient elution from 5% to 70% MeCN in water (constant 0.1% formic acid) over 20 minutes, isocratic elution with 70% MeCN from 20 to 30 min, and returned to initial conditions and equilibrated for 5 min) to give compound PTAD-DBCO-FITC precursor (6.0 mg, 0.006 mmol, 66%, retention time 22 min) as yellow solid. **^1^**H NMR (400 MHz, methanol-*d*^4^) δ 8.23 (s, 1H), 7.99 (d, *J* = 8.0 Hz, 1H), 7.54 (d, *J* = 8.0 Hz, 2H), 7.45 (s, 2H), 7.29 (d, *J* = 4.0 Hz, 1H), 7.25-7.23 (m, 3H), 7.29 (t, *J* = 8.0 Hz, 2H), 7.15 (d, *J* = 8.0 Hz, 1H), 6.95 (d, *J* = 8.0 Hz, 1H), 6.86 (d, *J* = 8.0 Hz, 1H), 6.60 (s, 2H), 6.51 (q, *J* = 12.0 Hz, 2H), 6.46-6.42 (m, 2H), 5.94 (d, *J* = 20.0 Hz, 1H), 4.42 (t, *J* = 16.0 Hz, 1H), 4.36 (t, *J* = 8.0 Hz, 1H), 3.40-3.30 (m, 2H), 2.24-2.08 (m, 2H), 1.96-1.74 (m, 2H). **HRMS (TOF)** Calculated for [M+Na]^+^ = 919.2452, found = 919.2436. *In-situ* oxidation of this PTAD-DBCO-FITC precursor in DMF by equal equivalent NBS provided PTAD-DBCO-FITC (**8**, 10 mM in DMF, stored at −20 C°), which was used directly without further purification.

The red fluorescent probe PTAD-DBCO-Cy5 (**10**).

A solution of compound **PTAD-N_3_** (3.4 mg, 0.0128 mmol) and DBCO-Cy5 (5.0 mg, 0.0064 mmol) in dry CH3CN (1.0 mL) under nitrogen was stirred at room temperature overnight. The reaction mixture was purified by RP-HPLC (Separation was achieved using a Ultimate XB-C_18_ column, 21.2 × 250 mm 5 micron (Welch Materials *Inc.,* Shanghai, China) by gradient elution from 5% to 70% MeCN in water (constant 0.1% formic acid) over 15 min, isocratic elution with 70% MeCN from 15 to 30 min, and returned to initial conditions and equilibrated for 5 min) to give PTAD-DBCO-Cy5 precursor (6.0 mg, 0.0057 mmol, 90%, retention time 24 min) as blue solid, which was re-dissolved in anhydrous DMF (0.3 mL) and was treated equal equivalent NBS solution (20 mM in DMF) to prepare PTAD-DBCO-Cy5 stock solution (**10**, 10 mM in DMF, stored at −20 C°). **MS (ESI)** Calculated for [M-H]^-^ = 1035.5, found [M-H]^-^ = 1034.7. **HRMS (TOF)** Calculated for [M-Cl]^+^ = 1003.4977, found 1003.4964.

### Nucleotide i^6^A triphosphate synthesis

Nucleoside i^6^A (35 mg, 0.1 mmol) and proton sponge (58 mg, 0.25 mmol) were dissolved in trimethyl phosphate (0.7 mL) and was placed under the ice-water bath. phosphorus oxychloride (33.6 μL, 0.36 mmol) was slowly added and stirred at 0 °C for 12 hours. A solution of tributylamine (175 μL, 0.72 mmol) and tributyl ammonium pyrophosphate (642 mg) in DMF (2.0 mL) was slowly added and the reaction mixture was stirred at 0 °C for 30 minutes. The reaction was quenched by addition of 1.0 M aqueous TEAB (pH = 7.5, 15 mL). The mixture was diluted with H2O (5.0 mL) and subjected to HPLC purification. Separation was achieved using an Ultimate XB-C_18_ column, 21.2 × 250 mm 5 micron by gradient elution from 5% to 50% MeCN in water (constant 0.1% formic acid) over 25 minutes, isocratic elution with 50% MeCN from 25 to 30 minutes, and returned to initial conditions and equilibrated for 5 minutes to give **i^6^A-TP** (10.0 mg, as tributyl ammonium salt, retention time 20 minutes). **^1^H NMR** (400 MHz, methanol-*d*^4^) δ 8.40 (s, 2H), 8.30 (s, 1H), 6.13 (d, *J* = 4.0 Hz, 1H), 5.40 (t, *J* = 12.0 Hz, 2H), 4.73 (q, *J* = 16.0 Hz, 1H), 4.57-4.43 (m, 1H), 4.27-4.26 (m, 2H), 4.20-4.13 (m, 2H). The proton peak of methyl group in prenylated functionality was covered by tributyl ammonium salt in **^1^H NMR**. **^31^P NMR** (162 MHz, methanol-*d*^4^) δ −10.47 (d, *J* = 22.68 Hz, α-P), −11.46 (d, *J* = 22.68 Hz, γ-P), −24.00 (t, *J* = 43.74 Hz, β-P). **MS (ESI)** Calculated for [M-H] ^-^ = 574.1, found [M-H] ^-^ = 574.2.

### Profiling the dynamic of the Ene-ligation of i^6^A nucleoside with PTAD

Nucleoside i^6^A (10 mg, 0.03 mmol) were dissolved in CD_3_CN/ D_2_O (0.5 mL, v/ v = 4: 1, final concentration = 60 mM). 4-Phenyl-3H-1,2,4-triazole-3,5(4H)-dione (PTAD, 26 mg, 0.15 mmol, 3.0 equivalent) was then added to the solution (final concentration = 300 mM), and the reaction mixture was allowed to stand at room temperature. The **^1^H NMR** spectra were recorded by **^1^H NMR** (400 MHz, CD_3_CN/ D_2_O).

### Fluorescence labeling of i^6^A in the metabolites of eukaryotic cells

First culture the HeLa cells in a six-well plate to 70%-80%. Then add different concentrations of i^6^A (**4**) to DMEM medium (100 *μ*M, 200 *μ*M and 400 *μ*M of i^6^A). After 12 h of culture, collect all cells and extract total RNA.

RNA (i^6^A-incoporated) labelled by fluorescein PTAD-DBCO-FITC (**8**).

To the 1.5 mL Eppendorf tube containing PTAD-DBCO-FITC precursor (**7**, 10 *μ*L, 10 mM solution in DMF) was added *N*-bromo succinimide (1.0 *μ*L, 100 mM in DMF). The mixture was vortexed gently and formation of the light-red color was observed. The reagent was kept on ice and used for the i^6^A-incorporated RNA labelling immediately. The i^6^A-incoporated total RNA (10 μl, 20 ng/ μl in DEPC water) was incubated with PTAD-DBCO-FITC (**8**, 10 μl, 0.1 mM), shaken for 30 minutes at 0 °C. RNA was analyzed by gel electrophoresis on 1 % agarose. The electrophoresis conditions were as follows: power supply (DYY-6C power supply, Liuyi Biotechnology) was set to 130 V. 1% agarose gels were run at room temperature (25 °C) for 20 minutes and stained with or without GoodView™ (SBS Genentech, Beijing, China). A DNA size marker (1 kb DNA Ladder, TsingKe Biotech, TSJ102, Beijing, China) was used. Gels were analyzed by in-gel fluorescence measurements on a FluorChem^®^ FC3 imager (Alpha Innotech). Fluorescence was measured with a blue light excitation wavelength (475 nm) and a green filter emission (537 nm).

RNA (i^6^A-incoporated) labelled by fluorescein PTAD-DBCO-Cy5 (**10**).

To the 1.5 mL Eppendorf tube containing PTAD-DBCO-Cy5 precursor (**9**, 10 *μL,* 10 mM solution in DMF) was added *N*-bromo succinimide (1.0 *μ*L, 100 mM in DMF). The mixture was vortexed gently and formation of the light-red color was observed. The reagent was kept on ice and used for the i^6^A-incorporated RNA labelling immediately. The i^6^A-incoporated total RNA (10 μl, 20 ng/ μl in DEPC water) was incubated with PTAD-DBCO-Cy5 (**10**, 10 μl, 0.1 mM), shaken for 30 minutes at 0 °C. RNA was analyzed by gel electrophoresis on 1 % agarose. The electrophoresis conditions were as follows: power supply (DYY-6C power supply, Liuyi Biotechnology) was set to 130 V. 1% agarose gels were run at room temperature (25 °C) for 20 minutes and stained with or without GoodView™ (SBS Genentech, Beijing, China). A DNA size marker (1 kb DNA Ladder, TsingKe Biotech, TSJ102, Beijing, China) was used. Gels were analyzed by in-gel fluorescence measurements on a FluorChem^®^ FC3 imager (Alpha Innotech). Fluorescence was measured with a blue light excitation wavelength (475 nm) and a green filter emission (537 nm).

### Iodine-mediated cycloaddition of nucleoside i^6^A

The reactions were carried out in DMSO-*d*^6^ in order to facilitate *in-situ* NMR characterization without further purification. i^6^A (**4**, 0.15 mmol) was dissolved in DMSO-*d*^6^ (0.5 mL), and iodine (0.45 mmol in DMSO-*d*^6^) was added. The mixture was stirred at 37 °C for 5 minutes and was taken out for **^1^H NMR**.

### Plausible reaction pathway predication of Iodine-mediated cycloaddition of nucleoside i^6^A

All the calculations were performed with the Gaussian 16, Revision B.01 program. Geometry optimization of the model systems in the gas phase were carried out with the B3LYP/ BS1 DFT method augmented with the D3(BJ) version of Grimme’s empirical dispersion correction (BS1 denotes a basis set combining SDD for I and 6-31G (d, p) for other atoms). Frequency calculations were performed at the same level of theory to determine whether the optimized structures are minima (no imaginary frequencies) or saddle points (one imaginary frequency) on the potential energy surface, and to provide thermal corrections to the Gibbs free energies. The solvation energy corrections were computed at the B3LYP/ BS1 level with the SMD solvation model for DMSO on gas-phase optimized geometries. The single point energies were computed with M062X/ BS2 augmented with the D3 version of Grimme’s empirical dispersion correction (BS2 denotes a basis set combining SDD for I and Def2TZVP for other atoms). Gaussian 16 Citation: Gaussian 16, Revision B.01, M. J. Frisch, G. W. Trucks, H. B. Schlegel, G. E. Scuseria, M. A. Robb, J. R. Cheeseman, G. Scalmani, V. Barone, G. A. Petersson, H. Nakatsuji, X. Li, M. Caricato, A. V. Marenich, J. Bloino, B. G. Janesko, R. Gomperts, B. Mennucci, H. P. Hratchian, J. V. Ortiz, A. F. Izmaylov, J. L. Sonnenberg, D. Williams-Young, F. Ding, F. Lipparini, F. Egidi, J. Goings, B. Peng, A. Petrone, T. Henderson, D. Ranasinghe, V. G. Zakrzewski, J. Gao, N. Rega, G. Zheng, W. Liang, M. Hada, M. Ehara, K. Toyota, R. Fukuda, J. Hasegawa, M. Ishida, T. Nakajima, Y. Honda, O. Kitao, H. Nakai, T. Vreven, K. Throssell, J. A. Montgomery, Jr., J. E. Peralta, F. Ogliaro, M. J. Bearpark, J. J. Heyd, E. N. Brothers, K. N. Kudin, V. N. Staroverov, T. A. Keith, R. Kobayashi, J. Normand, K. Raghavachari, A. P. Rendell, J. C. Burant, S. S. Iyengar, J. Tomasi, M. Cossi, J. M. Millam, M. Klene, C. Adamo, R. Cammi, J. W. Ochterski, R. L. Martin, K. Morokuma, O. Farkas, J. B. Foresman, and D. J. Fox, Gaussian, *Inc.,* Wallingford CT, **2016**.

### Iodine-mediated cycloaddition of nucleoside i^6^A

The model reactions were carried out in DMSO-*d*^6^ in order to facilitate *in-situ* NMR characterization without further purification. i^6^A (0.15 mmol) was dissolved in DMSO-*d*^6^ (0.5 mL), and iodine (0.45 mmol, 3.0 equivalent) was added. The mixture was stirred at 37 °C for 5 min and was taken out for **^1^H NMR, ^13^C NMR, HPLC, UV** and **LC-MS** analysis.

### Plausible reaction pathway predication of Iodine-mediated cycloaddition of nucleoside i^6^A

All the calculations were performed with the Gaussian 16, Revision B.01 program. Geometry optimization of the model systems in the gas phase were carried out with the B3LYP/ BS1 DFT method augmented with the D3(BJ) version of Grimme’s empirical dispersion correction (BS1 denotes a basis set combining SDD for I and 6-31G (d, p) for other atoms). Frequency calculations were performed at the same level of theory to determine whether the optimized structures are minima (no imaginary frequencies) or saddle points (one imaginary frequency) on the potential energy surface, and to provide thermal corrections to the Gibbs free energies. The solvation energy corrections were computed at the B3LYP/ BS1 level with the SMD solvation model for DMSO on gas-phase optimized geometries. The single point energies were computed with M062X/ BS2 augmented with the D3 version of Grimme’s empirical dispersion correction (BS2 denotes a basis set combining SDD for I and Def2TZVP for other atoms).

### Profiling method illustrations of i^6^A residues in cell

#### The efficiency of i^6^ATP in T7 RNA polymerase transcription system was detected *in vitro*

The *EGFP* mRNA sequence was obtained by transcribing under the conditions of i^6^ATP: ATP ratio of 0: 1, 3: 7, 7: 3, 1: 0 through the *in vitro* transcription system, and then detected each group of *EGFP* RNA sequence through iodine (0.5 M) treatment. After processing, the *c*DNA sequence of *EGFP* in each group was obtained by reverse transcription. Next, the obtained cDNA sequence was used as a template, and the mutant *EGFP* gene sequence in each group was obtained by PCR amplification, and cloned into the T vector. Through the amplification of DH5*α E. coli.,* 50 clones were picked from each group and sequenced to detect the mutation. After the mutation site of *EGFP* gene, and analyze the insertion efficiency of i^6^ATP.

#### Comparative analysis of the recognition efficiency of T7 RNA polymerase and RNA polymerase II for i^6^ATP in eukaryotic cells

T7 RNA polymerase and T7-IRES2-EGFP expression system were expressed in HeLa cells, and RNA was extracted 48 hours after transfection, and then the mixture was treated with iodine to cause i^6^ATP insertion site mutation. Then, the mutant sequences of the two endogenously expressed genes of *GAPDH* and *Notch1* were obtained by reverse transcription, and the mutant gene sequence of *EGFP* catalyzed by *T7 RNA polymerase* transcription was obtained. Same as the previous detection process, by detecting the insertion efficiency of i^6^ATP in the endogenous highly expressed gene *GAPDH* and the oncogene *Notch1* and the insertion efficiency of i^6^ATP in *T7-EGFP.* Compare the recognition efficiency of i^6^ATP by eukaryotic RNA polymerase and the feasibility of using it as a real-time transcriptome marker.

#### Data availability

The genome sequence data that support the findings of this study are available in NCBI GenBank database under accession nos. NC_000012.12, and NC_000009.12. For the corresponding genomes, see SI **Table E1-3**. All other data generated or analyzed in this study are available within the Article and its Supplementary Information and Source Data. Source data are provided with this paper.

## Results and Discussion

### Prenyl-transformations and the development of new labeling reagents

Over the past few years, we endeavored to create chemical conversions of a wide range of inert (poly)prenylated functionalities.^21–22^ For example, we found that 4-phenyl-1,2,4-triazoline-3,5-dione (PTAD) reacts with polyprenylated macromolecules in a superfast manner under physiological conditions. Similar reactivity has also been analyzed with the other two reagents including Selectfluor and nitroso derivatives, both have great application potential to study the biological prenyl functionality. The dynamic properties of the prenyl functionality have been extensively explored using our methods (see Supplementary Information (SI), Fig. S1, S9-S13 for details). We also discovered the pseudo first-order kinetics for the reaction involved prenyl molecule β-caryophyllene with PTAD. The calculated half-life time of 5 min and kinetic constant of 0.956 mol^-1^•s^-1^ demonstrated the great application potential of our labeling reagents to study prenyl functionalized biomolecules. In addition, we also demonstrated that electron-donating groups on PTAD accelerated the process in a superfast manner. ^23^ Further stability tests showed that both PTAD-labeling reagents and the resulting coupling adduct are relatively inert to the external environment.

With these chemical tools in hand, we first synthesized the i^6^A nucleoside (**4** in **Fig. 2**) through a linear 4-step process in an overall 69% yield (SI, Scheme S1, Fig. S2-5), and started to test its compatibility with these labeling reagents. A rapid reaction was observed between i^6^A and PTAD (**5**) (**Fig. 2A**) with a clear color change in less than 5 min. The resulting products were monitored by in-situ NMR (**Fig. 2B**), confirming that this reaction can take place quickly with good targeting specificity in biocompatible conditions.

**Figure 2.**
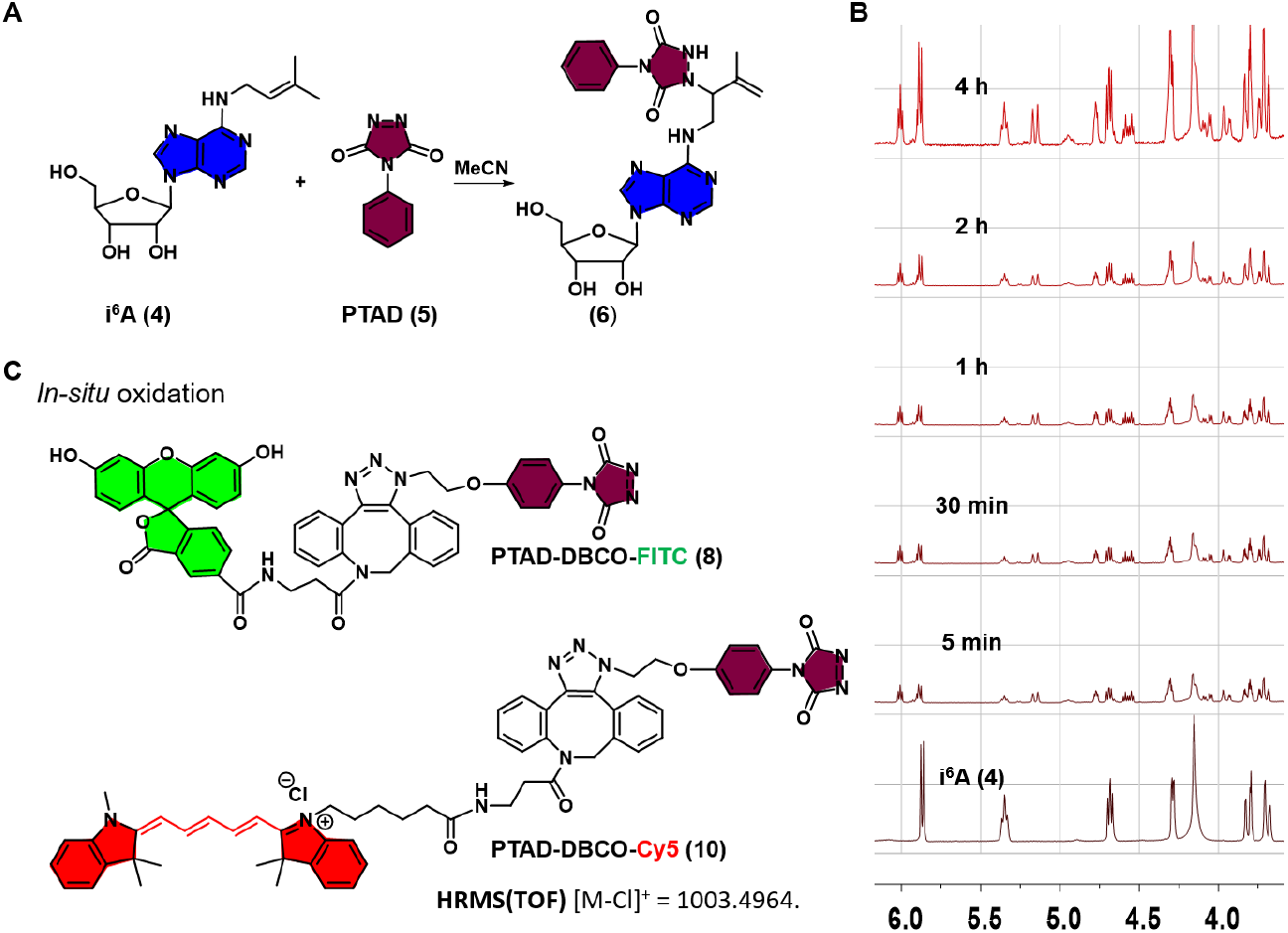
The reactions of i^6^A (**4**) with various functionalities. **(A)** Reaction of i^6^A (**4**) with PTAD (**5**) at r.t. **(B)** Proton NMR was used to monitor the reaction of i^6^A (**4**, 0.03 mmol, 30 mM) with PTAD (0.15 mmol) in CD_3_CN/D_2_O (v/v, 1/1) at r.t. The rapid reaction resulted in approximately 50% conversion within 5 min (peak (5.5 ppm, triplet) is assigned to alkenyl site in prenyl group of i^6^A). From top to bottom: 4 h, 2 h, 1 h, 30 min, 5 min and 0 min. **(C)** Chemical structure of the green fluorescent probe PTAD-DBCO-FITC (**8**), and chemical structure of the red fluorescent probe PTAD-DBCO-Cy5 (**10**).

To further expand this method and take advantage of the fluorescence labeling using this chemistry, we constructed two fluorescent probes derived from the structures of PTAD. (**8** and **10**, **Fig. 2C**, and SI, Scheme S2-3 and Fig. S6-8). The strain-promoted click reaction of azido PTAD with FITC-DBCO (SI, Scheme S2 and Fig. S6-7) or Cy5-DBCO (SI, Scheme S3 and Fig. S8) proceeded smoothly to provide fluorescent precursors, followed by the *in-situ* oxidation with *N*-bromosuccinimide (NBS) to yield fluorescent probes **8** and **10**. In the meantime, we synthesized the i^6^A triphosphates (i^6^A-TP) for its enzymatic incorporation into RNA strands. Based on previous studies, i^6^A-TP was obtained in a linear 2-step process with 30% of isolated yields (SI, Scheme S4, Fig. S14-15). All the data were consistent with the literature results.^27–28^ The peaks in the phosphorus NMR spectra at −10.5, −11.5 and −24.0 ppm were assigned to α, γ and β phosphorus atoms respectively, each with a *J* value of 26.8 Hz.

### Fluorescence labeling of i^6^A in the metabolites of eukaryotic cells

With the two fluorescent labeling reagents and i^6^A-TP building block in hands, we asked whether i^6^A nucleoside can be incorporated by RNA polymerases in the process of transcription within eukaryotic cells, and if so can they be recognized and fluorescently labeled (**Fig. 3**). To investigate this, we cultured HeLa cells to 70%-80% confluence in a 6-well plate, followed by adding different concentrations of i^6^A (**4**, 100 μM, 200 μM, 400 μM) in DMEM. Subsequently, the CMV-T7 and T7-IRES2-EGFP plasmids, as well as their conjugate units (SI, Table S1-3 and Fig. S16-23) were transfected respectively in HeLa cells for 36 hours before the measurement was taken for the expression of EGFP protein by fluorescence microscopy. As shown in **Figure 3A**, the expression of green fluorescent protein can only be detected in the presence of CMV-T7 and T7-IRES2-EGFP or their integration counterparts, which indicated the successful formation of specific transcripts of the target genes by T7 RNA polymerase in eukaryotic cells.

**Figure 3.**
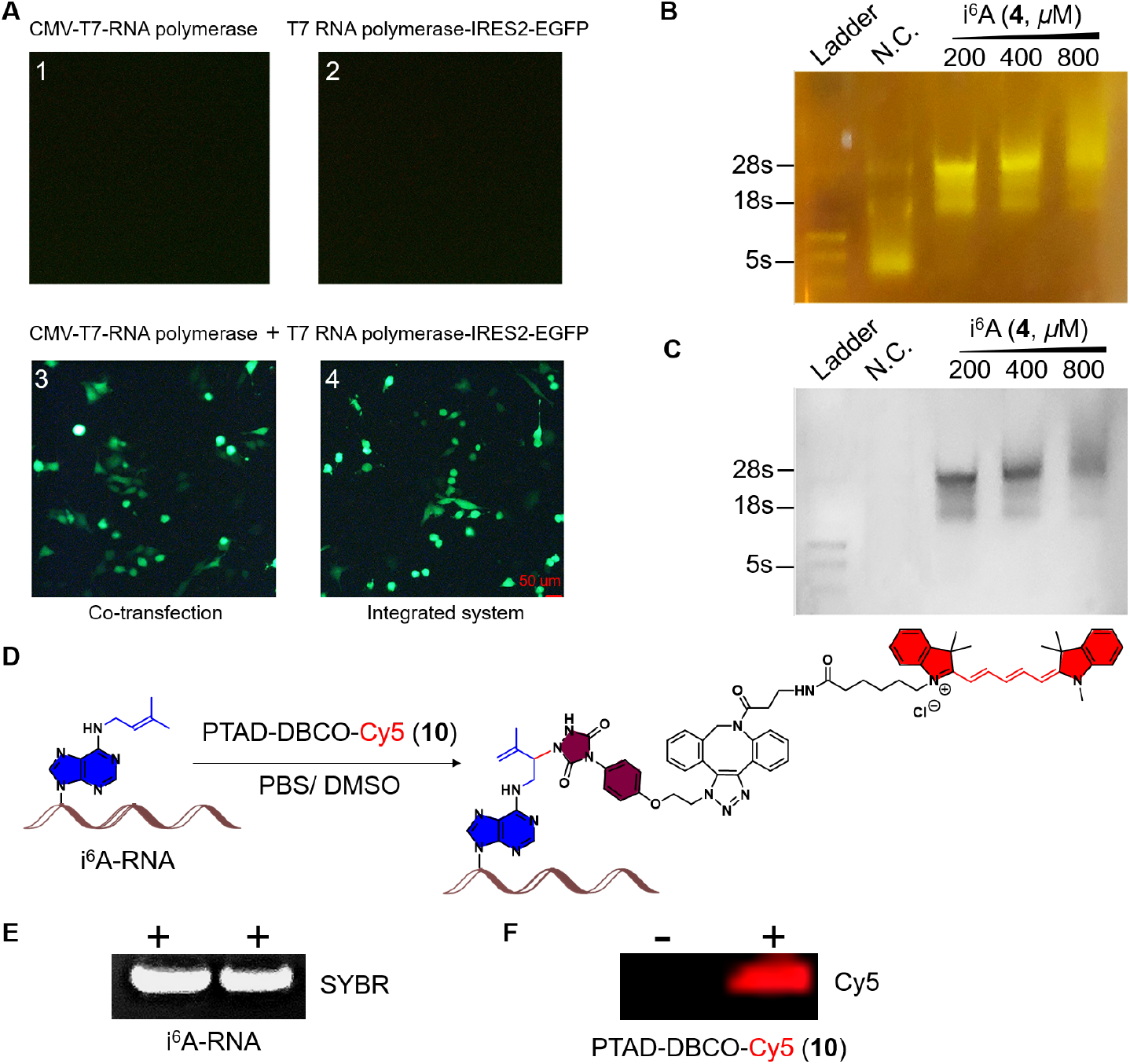
Investigation of the synthesized i^6^A (**4**)-incorporated RNAs in eukaryotic cells and its fluorescent labeling. (A) Expression level of EGFP green fluorescent protein initiated by the T7 RNA polymerase transcription system in HeLa cells. A1-A4 represent the transfection of CMV-T7 RNA polymerase, T7 RNA polymerase-IRES2-EGFP, CMV-T7 RNA polymerase with T7 RNA polymerase-IRES2-EGFP, and integrated CMV-T7 RNA polymerase-T7 RNA polymerase-IRES2-EGFP respectively. Only cotransfection or integrated systems of CMV-T7 RNA polymerase and T7 RNA polymerase-IRES2-EGFP displayed green fluorescent signals. (B) Detection of the green fluorescent signal in i^6^A-incorporated RNAs using PTAD-DBCO-FITC (**8**, 0.1 mM). (C) Goldview dye staining of i^6^A-incorporated RNAs. (D) Fluorescent labeling of i^6^A-incorporated RNAs using the well-designed PTAD-DBCO-Cy5 (**10**, 0.1 mM). (E) Western blotting assays of RNAs containing i^6^A *via* reverse transcription. (F) Red fluorescent imaging of PTAD-DBCO-Cy5 (**10**)-labeled i^6^A-RNAs in a gel. Bar scale: 50 μM.

For the labeling experiment, HeLa cells were harvested and the total RNAs were extracted 12 hours after the transfection, which were subsequently incubated with our PTAD-DBCO-FITC (**8**) or PTAD-DBCO-Cy5 (**10**) respectively. After excess washing, the purified RNAs were evaluated for the incorporation of i^6^A residues, and the overall PTAD-DBCO-FITC (**8**) fluorescent labeling efficiency was determined by agarose gel electrophoresis (SI, Fig. S25-27). Both the modified and the wild-type RNAs appeared to have similar stability with i^6^A incorporated as observed in Goldview nucleic acid dye staining (**Fig. 3B-C**). In addition, the rRNA bands representing 5s and 18s were significantly shifted upward compared with the wild-type one. By contrast, no significant change was observed in the rRNA band at 28s. Notably, the fluorescence intensity of the RNAs containing i^6^A decreased when the level of i^6^A reached the cytotoxic concentration of 800 μM (**Fig. 3B-C**, and Fig. S25-27). No fluorescence signal was detected in the control group in comparision to a distinct fluorescence signal observed in the total RNAs incorporated with i^6^A and treated with PTAD-DBCO-Cy5 (**13**) probe (**Fig. 3D-F**). Our assays indicated that i^6^A nucleosides could be recognized and used in transcription by RNA polymerase and the resulting RNAs are detectable by our fluorescent probes.

### Development of iodine-mediated oxidation and reverse transcription (IMORT) method for i^6^A profiling

We further examined the prenyl modification on RNAs ^24–26,29^ via the addition of I_2_, an oxidative reagent utilized broadly in the cycloaddition reaction of a prenyl group.^30–32^ As expected, the reaction of i^6^A with optimized concentration of iodine reagent proceeded very smoothly to yield the full conversion of i^6^A in less than 5 min (**Fig. 4A-B** and Fig. S28-S39 in SI). The NMR and mass spectrometry results confirmed the spontaneous formation of the cycloaddition adducts **3a** or **3b** (Fig. S28-S38).

**Figure 4.**
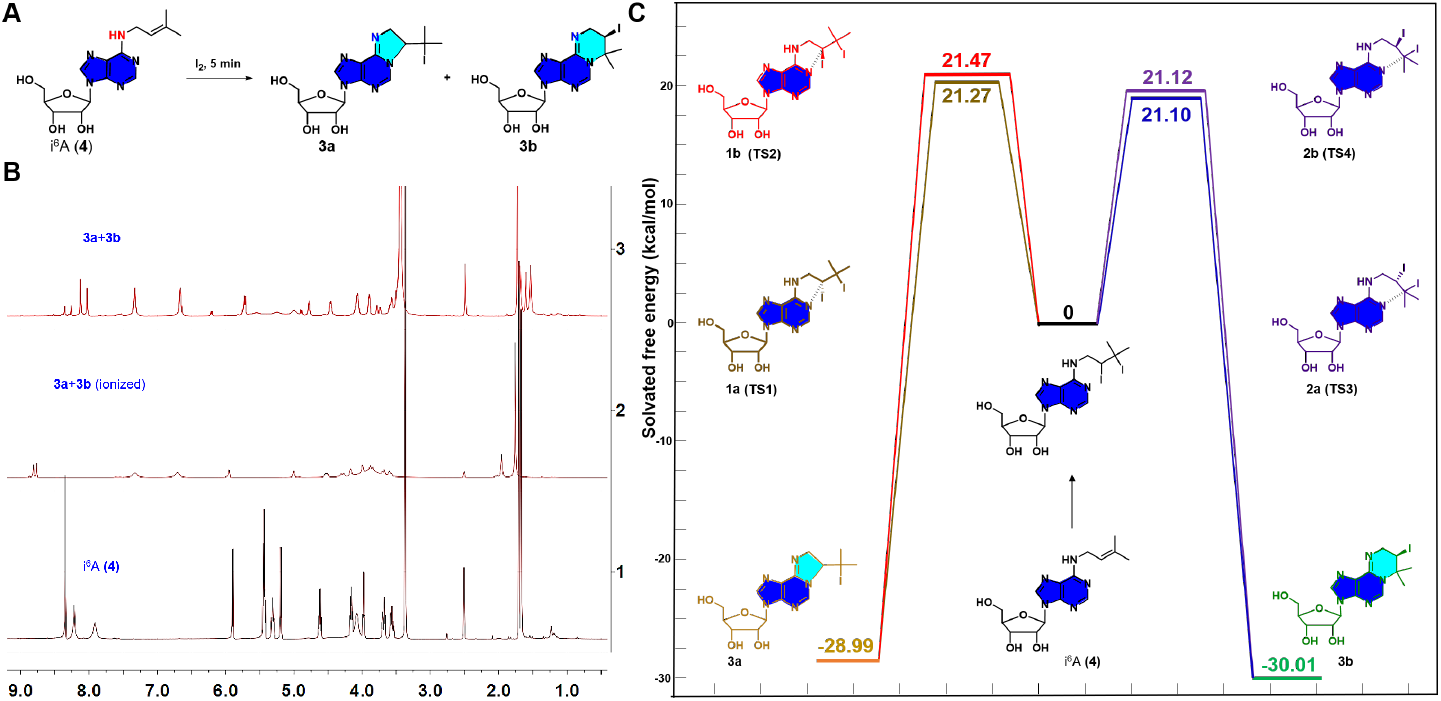
Investigation of the chemical transformations of prenylated nucleoside i^6^A (**4**). **(A)** Reaction of i^6^A (**4**, 0.15 mmol) with iodine (0.45 mmol) at r.t. for 5 min. **(B)** Comparative analysis using proton NMR for the reaction of i^6^A (**4**) and I_2_ with and without sodium persulfate. **(C)** DFT calculations for the possible product of the reaction of i^6^A (**4**) with I_2_. All the calculations were performed using the Gaussian 16, Revision B.01 program.

DFT calculations were performed to elucidate the reaction pathway and mechanism. The addition of I_2_ resulted in the spontaneous attachment of di-iodinyl atoms to the double bond of the prenyl group with four potential isomers formed because of the chirality of the products (**1a-1b, 2a-2b**) (**Fig. 4C** and Fig. S40 for details). Two plausible reaction pathways have been identified thus far involving the formation of either five-membered or six-membered intermediates through a rearrangement process. Between the two products, compound **3b** has been demonstrated to be more abundant based on the extensive NMR studies,^26^ even though **Fig. 4C** showed very small difference among the activation barriers of the four isomers. As a result, the iodine treatment followed by quenching with sodium persulfate converted i^6^A from an H-bond donor to a cyclic H-bond acceptor, which might change the enzyme recognition and cause base mutation during reverse transcription (RT) process, indicating the potential application for the detection of the prenyl functionality in endogenous RNAs at a single-base resolution.

### Profiling of i^6^A residues in cell

We subsequently applied this IMORT method to detect i^6^A modification sites in transcriptome wide studies. *EGFP* gene (see SI, Fig. S42-44) was chosen as RNA substrate to model the incorporation of i^6^A in cellular RNAs. In the course of cellular metabolism, living HeLa cells were fed with 30% and 70% i^6^A nucleoside in two groups respectively following the optimized protocol. The resulting EGFP RNA from each experimental group was further treated with I_2_ (0.5 M). Each EGFP RNA was then reverse transcribed (see SI, Fig. S41, 45-47 for details) into *c*DNA followed by PCR amplification. Each amplified EGFP amplicon was next cloned into the T-vector and transformed into DH5α E. coli. A total of 50 clones were picked from each group, followed by high-throughput sequencing and comparing with the native control to determine the sites of mutation on EGFP gene and to analyze the insertion efficiency of i^6^A. As anticipated, adenosine with prenyl modification displays normal base pairing with thymidine (A≡T). By contrast, the addition of iodine induced the transformation of i^6^A to its cyclic form, thereby diminishing the normal base pairing specificity and resulting in a mixture of complementary G/C/U (**Fig. 5A**). The base-mutation rates could be further measured by the sequencing data. It turns out that in the presence of 30% of i^6^ATP, the resulting A bases were significantly changed (**Figs. 5B1** and **5B3**), whereas G/C/U displayed statistically minimal effect (**Figs. 5B2** and **5B4**). By contrast, the I_2_-treated i^6^A shows a significant mutation rate, as shown in **Fig. 5C1** and **5C2** (also see **Extended Data Table E4** in SI for details), which demonstrated that our method could be employed to detect the i^6^A sites in RNAs.

**Figure 5.**
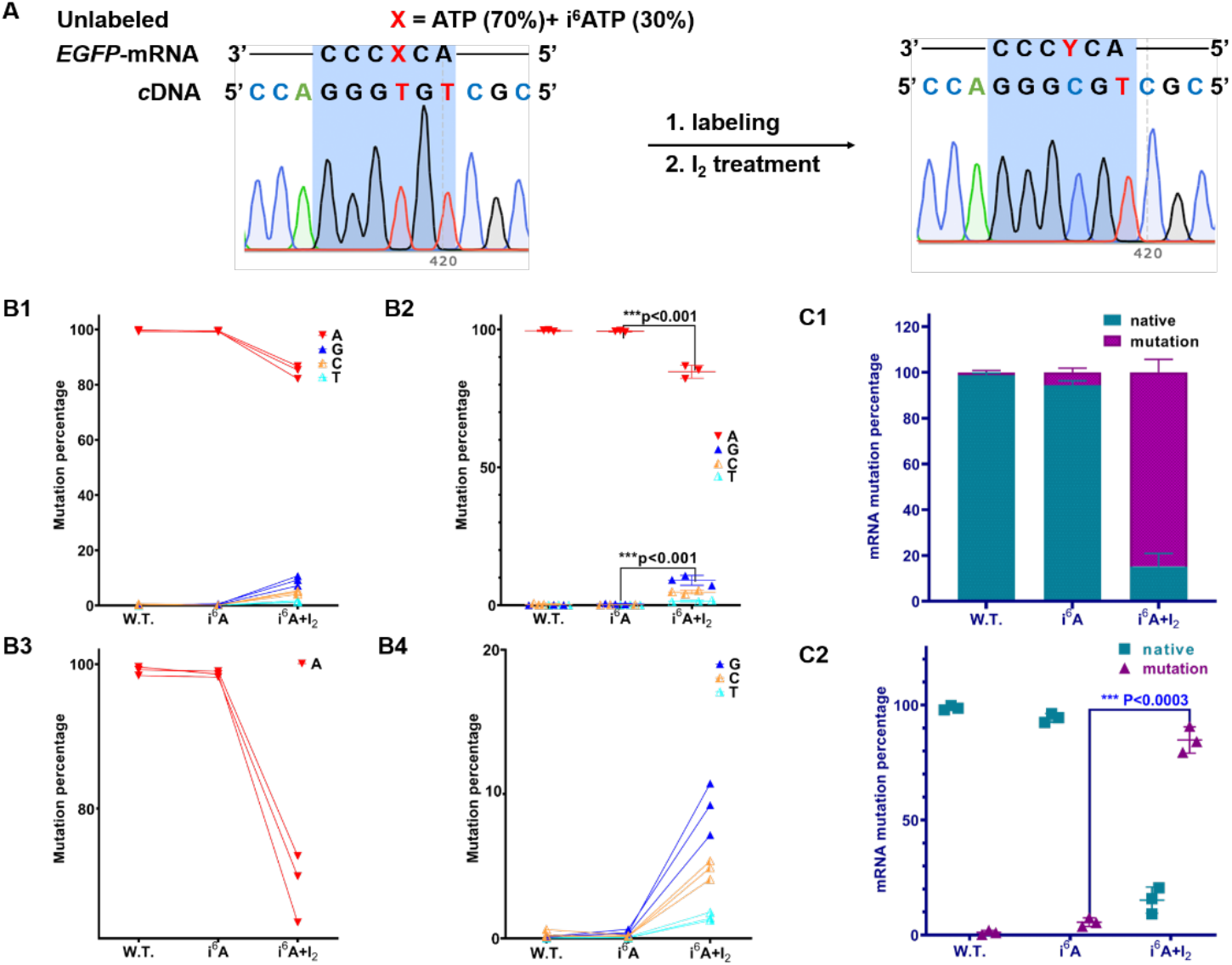
Detection of i^6^A sites in RNA at a single-base resolution. **(A)** Detection scheme for i^6^A modification sites in RNA by employing i^6^ATP in *in-vitro* studies. Schematic of i^6^A insertion at a specific RNA site and chemical-induced mutation with I_2_ treatment. **(B)** Insertion of i^6^ATP at a specific site of transcribed RNA followed by chemical mutation after I_2_ treatment. A comparative analysis of i^6^A-incorporated RNA with and without iodine treatment showed that I_2_ treatment is a powerful tool for detecting i^6^A in nucleic acids. **(C)** Determination of mRNA labeling efficiency by i^6^ATP under in vitro transcription conditions.

## Conclusion

In summary, we used our recently developed Ene-ligation bioorthogonal chemistry and new PTAD-based fluorescence probes for the direct labeling of the i^6^A containing RNAs in cell. We synthesized the i^6^A triphosphate building block and demonstrated that it can be recognized and incorporated by eukaryotic RNA polymerase through intracellular transcription with adjustable i^6^A concentrations, resulting in the i^6^A-modified RNA that can be directly labeled by our PTAD-based fluorescent dye. We also found that iodine-mediated cycloaddition reactions of the prenyl group occurs in a superfast manner, and converts i^6^A from a hydrogen-bond acceptor into a donor. Based on this reactivity, we developed a new iodine-mediated oxidation and reverse transcription sequencing method to profile the i^6^A residues in cellular RNAs. The biological functions of prenyl-modifications in both RNA and protein have been increasingly appreciated. For example, the expression levels of prenylated 2’-5’-oligoadenylate synthetase 1 (OAS1) has been closely associated with the severeness of hospitalized SARS-CoV-2 patients, indicating critical roles that prenylation could play in human antiviral defense system.^33^. Our new chemical biology methods developed in this work will provide unique toolset for detecting, profiling and monitoring the i^6^A residues in both normal and diseased cell environments, shedding light on more systematic functional studies of this natural modification.

## Acknowledgements

Rui Wang thanks the starting fund from Huazhong University of Science and Technology for financial support. “One-Hundred-Talents” Youth Program of Hubei Province is appreciated. Jia Sheng thanks the funding support from US National Science Foundation (CHE-1845486). Jason Chin is high appreciated for generous gift of the plasmid. We thank Prof. Sydney Hecht and Prof. Maxim Royzen for valuable comments. We also thank Prof. Zhipeng Zhou from Huazhong Agricultural University for valuable comments about RNA sequencing data.

## Author contributions

R. W. and J.S., L.W. designed the project. S.W., Y.Y.L, H.L.Z., Y.F.G.,and J.J.Z. performed compound isolation and structural elucidation under the supervision of R.W., J.S., and S. W., H.L.Z. performed the chemical synthesis of PTAD-based probes. H.L.Z, Y.Y.L, and Y.F.G conducted bioassays. Y.Y.L, and S.W. conducted NMR spectra measurements. H.L.Z, Y.Y.L, Y.F.G., Y.Y.Z., and L.W. conducted EGFP expression and purification, and some bioinformatic studies. R.W., and J.S. wrote the manuscript, with input from all co-authors.

## Supplementary information

The online version contains supplementary material available at NAR online.

## Correspondence and requests for materials

should be addressed to Rui Wang (Huazhong University of Science and Technology) or Jia Sheng (University at Albany SUNY).

