## Supplemental Information for "Bioorthogonal in-cell Labeling and Profiling of *N*^6^-isopentenyladenosine (i^6^A) Modified RNA"

### 1. Materials

All chemicals were purchased from Sigma-Aldrich, Acros, Inno-chem, Macklin Inc, Energy Chemical and were used without further purification. Extra dry solvents, such as 1,4-dioxane, DMF, and THF, were obtained from Inno-chem in sealed bottles over 3 or 4 Å molecular sieves and stored under dried nitrogen. Organic solvents were obtained from Sinopharm Chemical Reagent Co., *Ltd* (Shanghai, China) and used in reactions, column chromatography and recrystallizations. Milli-Q ultrapure water (resistivity, 18 mΩ) purified through Millipore Milli-Q Advantage A1 purification system was used for all bioconjugation reactions. The reactions were monitored by thin-layer chromatography (TLC) analysis using silica gel (60-Å pore size, F254, Yantai Chemical Industry Research Institute) plates. Compounds were visualized by UV irradiation (λ = 254 nm) and/or spraying TLC stain such as a $KMnO_4$ solution followed by electronic heating. Flash chromatography columns were performed on silica gel (60-Å pore size, 230−400 mesh). HPLC purification were performed using EasyChrom-1000 system with NU3000 serials UV/Vis. detector (Hanbon Sci. & Tech., Jiangsu, China) using an Ultimate® XB-$C_{18}$ column, 21.2 × 250 mm 5 micron (Welch Materials Inc., Shanghai, China). Separation was achieved by gradient elution from 5% to 70% acetonitrile in water (constant 0.1% formic acid) over 20 min, isocratic elution with 70% acetonitrile from 20 to 25 min, and returned to initial conditions and equilibrated for 5 min. The LC chromatograms were recorded by monitoring absorption at 254 nm and 220 nm. $^{1}$H and $^{13}$C NMR spectra were recorded at room temperature on a Bruker spectrometer (AM-600 or AM-400) operating at 600/400 and 150/100 MHz, respectively. Chemical shifts are given in parts per million, and 1H and 13C {1H} NMR spectra were referenced using the solvent signal as an internal standard. The following abbreviations are used for the proton spectra multiplicities: s: singlet, d: doublet, t: triplet, m: multiplet, br: broad. Coupling constants (J) are reported in Hertz (Hz). HRMS (TOF) were obtained from the Bruker Micro TOF II Spectrometer using Electro Spray Ionization (ESI). MS (ESI) was obtained from the Expression L (Beijing Bohui Innovation Biotechnology Co., *Ltd*). LC-MS analysis was obtained from the Bruker Orbitrap LC/MS

(Q Exactive™) at *Huazhong University of Science and Technology Analytical and Testing Center*. UV–visible absorbance measurements were performed UV–visible spectrometer (Lambda365, PerkinElmer, German). Confocal Imaging was performed using a LSM 780 confocal microscope (Zeiss) with a 20× objective at 16-bit depth under non-saturating conditions. EGFP were imaged with a 480 nm (excitation) and a 510 nm (emission) and false-colored green.

Materials. Streptococcus pyogenes (product # M0646), T7 Endonuclease I (product # M0302) Ribonucleotide solution mix (NTPs) and deoxy-ribonucleoside triphosphates (dNTPs) were purchased from New England Biolabs (USA). Transcript Aid T7 High Yield Transcription kit (product # K0441) and Glycogen (product # R0561) were purchased from Thermo Fisher Scientific. Pyrobest™ DNA Polymerase and PrimeSTAR HS DNA Polymerase were purchased from TaKaRa Shuzo Co. Ltd. (Tokyo, Japan). DNA Clean & Concentrator™-5 kit (product # D4014) was purchased from Zymo Research Corp. The DNeasy Blood & Tissue Kit was purchased from QIAGEN. The oligonucleotides at HPLC purity were obtained from TaKaRa company (Dalian, China). The nucleic acid stains Super GelRed (NO.: S-2001) was bought from US Everbright Inc. (Suzhou, China). Thiazolyl Blue Tetrazolium Bromide (MTT, CAS # 298-93-1) were purchased from Sigma-Aldrich Inc. (Shanghai, China). DPBS (CAS # 63995-75-5) was purchased from TCI (Shanghai) Development Co., Ltd.

### 2. Preliminary rearrangement investigations.

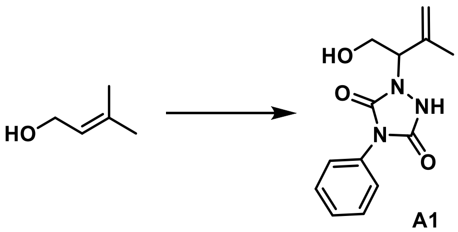

**A1**

To a stirred solution of prenyl alcohol (8.6 mg, 0.1 mmol) was added 4-phenyl-3H-1, 2, 4-triazole-3, 5(4H)-dione (PTAD, 17 mg, 0.1 mmol) in acetonitrile (0.6 mL). The mixture continued for 12 hours. Solvent was removed and the crude product was purified by flash chromatography (eluent: PE: ethyl acetate = 10:1) to afford yellowish oil product **A1** (20 mg, 0.077 mmol) in 82% yield.

**$^1$H NMR** (400 MHz, $CDCl_3$) δ 7.50-7.25 (m, 5H), 5.02 (s, 1H), 4.92 (s, 1H), 4.64-4.61 (m, 1H), 4.00-3.95 (m, 1H), 3.88-3.84 (m, 1H), 1.75 (s, 3H). **MS (ESI)** Calculated for $[M+H]^+$ = 262.1, found 262.0.

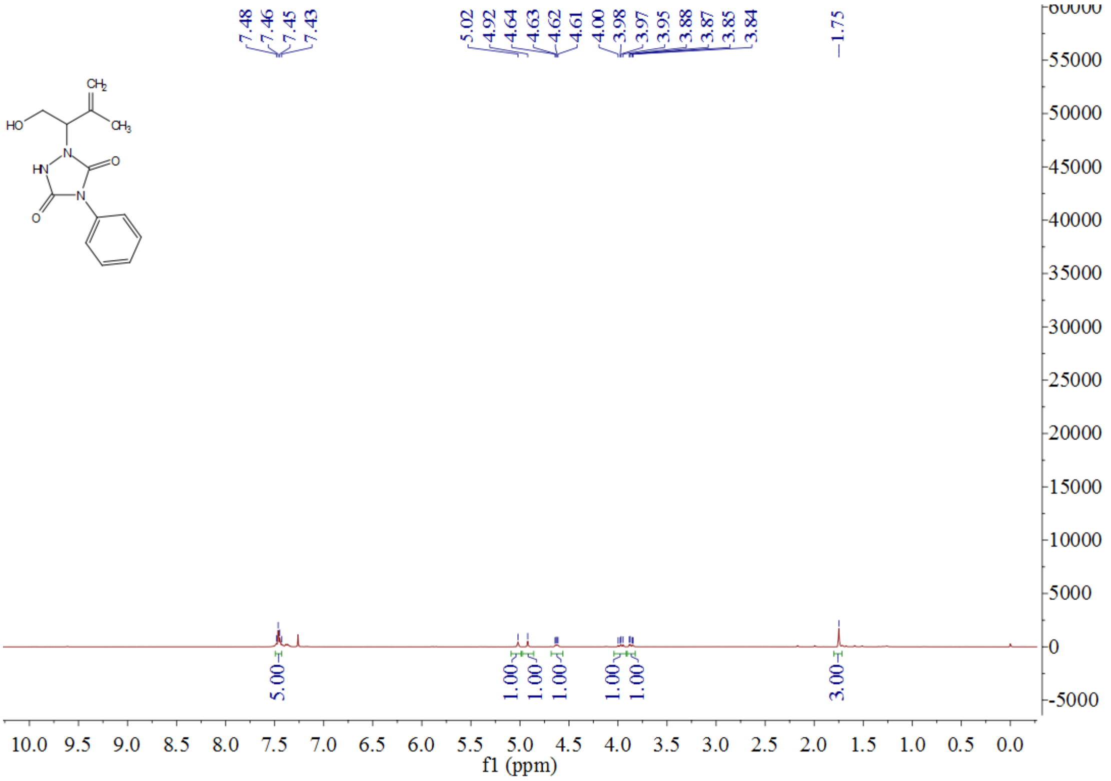

**Figure S1. $^1$H NMR** of crude product (**A1**) in $CDCl_3$, 400 MHz.

### 3. Synthesis nucleoside i⁶A (4).

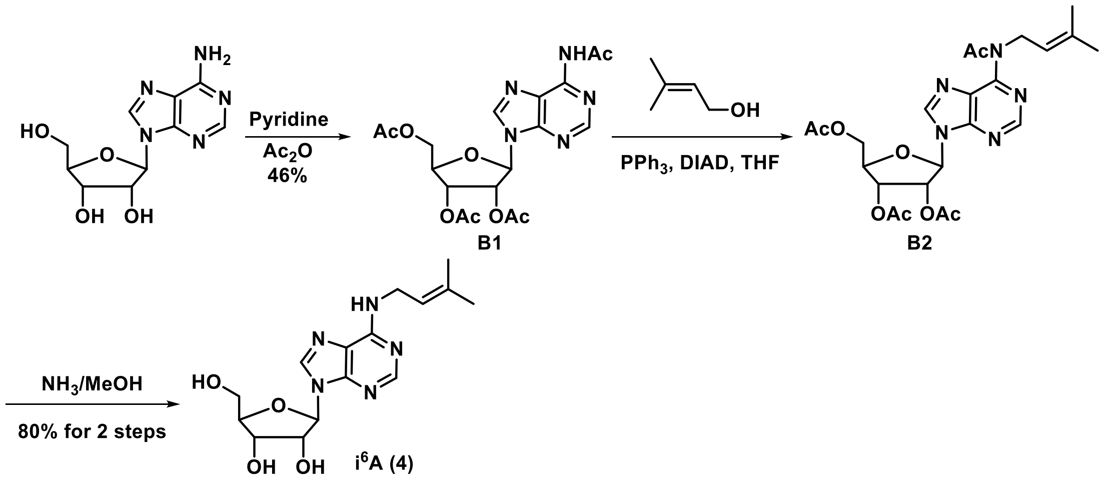

**Scheme S1**. Synthesis of i⁶A (**4**).

#### Synthesis of $N^6$-acetyl-2’, 3’, 5’-tri-*O*-acetyladenosine (B1).

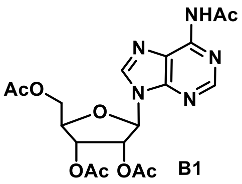
B1

A mixture of adenosine (2.0 g, 7.48 mmol), pyridine (15 mL), and $Ac_2O$ (7 mL, 74.2 mmol) was stirred at room temperature overnight. The resulting clear solution was refluxed at 60 °C overnight. The reaction was cooled down and quenched by EtOH. Then the reaction mixture was co-evaporated with addition of excess of EtOH to remove pyridine completely. The resultant foam was dissolved in MeOH (20 mL) and imidazole (0.4 g, 5.88 mmol) was added and the solution was stirred at room temperature. After 8 hours, the solution was diluted with ethyl acetate (50 mL) and washed by brine (5 × 50 mL). Then the organic solvent was dried over $Na_2SO_4$, filtered and concentrated. The residue was purified by silica gel ($CH_2Cl_2$: MeOH = 30: 1) to give **B1** (1.48 g, 3.4 mmol, 46%) as yellowish solid. **$^1$H NMR** (400 MHz, $CDCl_3$) δ 9.16 (s, 1H), 8.70 (s, 1H), 8.23 (s, 1H), 6.23 (d, *J* = 4.0 Hz, 1H), 5.97 (t, *J* = 8.0 Hz, 1H), 5.68 (t, *J* = 12.0 Hz, 1H), 4.48-4.44 (m, 2H), 4.41-4.36 (m, 1H), 2.63 (s, 3H), 2.16 (s, 3H), 2.12 (s, 3H), 2.09 (s, 3H). **$^{13}$C NMR** (101 MHz, $CDCl_3$) δ 170.80, 170.36, 169.60, 169.39, 152.66, 151.04, 149.52, 141.47, 122.23, 86.45, 80.40, 73.09, 70.61, 63.03, 25.75, 20.75, 20.54, 20.39. **MS (ESI)** Calculated for $[M+Na]^+$ = 458.1, found 458.0.

**Synthesis of *N*[6]-acetyl-2', 3', 5'-tri-*O*-acetyl-*N*[6]-(3-methylbut-2-enyl) adenosine (B2).**

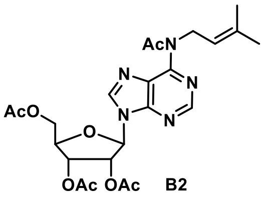

A mixture of **B1** (1.3 g, 3 mmol), triphenylphosphine ($PPh_3$, 1.2 g, 4.5 mmol), and 3-methyl-2-buten-1-ol (387 mg, 4.5 mmol) in THF (5.0 mL) was stirred at r.t. until a homogeneous solution was formed. Di-isopropyl azodicarboxylate (DIAD, 0.9 g, 4.5 mmol) was added in one portion. The reaction was monitored by TLC. A second addition of triphenylphosphine (4.5 mmol), alcohol and DIAD was made to achieve complete conversion of starting material **B1** after 20 hours. After 5 hours the mixture was evaporated and the residue was purified by column chromatography ($CH_2Cl_2$: MeOH = 50: 1) to give crude product **B2** (2.33 g, 4.63 mmol) as white solid, which was taken to the next step directly.

**Synthesis of** *N*[6]-(3-methylbut-2-enyl)-adenosine **(4,** i[6]A).

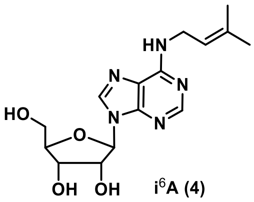

Compound **B2** (2.33 g, 4.63 mmol) was dissolved in 7.0 M $NH_3$ in MeOH solution (1.0 mL, 7.0 mmol) and the solution was stirred for 48 hours. The volatiles were evaporated under *vacuo* and the residue was purified by silica gel chromatography (PE: ethyl acetate = 1: 1) to give nucleoside i[6]A (**4**, 800 mg, 2.38 mmol, 80% over 2 steps) as white solid.

**$^1$H NMR** (400 MHz, DMSO-*d*[6]) δ 8.35 (s, 1H), 8.22 (s, 1H), 7.91 (s, 1H), 5.89 (d, *J* = 4.0 Hz, 1H), 5.45-5.42 (m, 2H), 5.31 (t, *J* = 16.0 Hz, 1H), 5.19 (d, *J* = 8.0 Hz, 1H), 4.62 (q, *J* = 16.0 Hz, 1H), 4.16 (q, *J* = 12.0 Hz, 1H), 3.98 (q, *J* = 8.0 Hz, 1H), 3.71-3.66 (m, 1H), 3.59 - 3.53 (m, 1H), 1.69 (d, *J* = 12.0 Hz, 6H). **$^{13}$C NMR** (101 MHz, DMSO-*d*[6]) δ 172.17, 154.80, 152.79, 140.14, 133.81, 122.43, 120.29, 88.45, 86.38, 73.97, 71.13, 62.14, 38.13, 25.84, 22.91, 18.29. **HRMS (TOF)** Calculated for $[M+H]^+$ = 336.1663, found 336.1690.

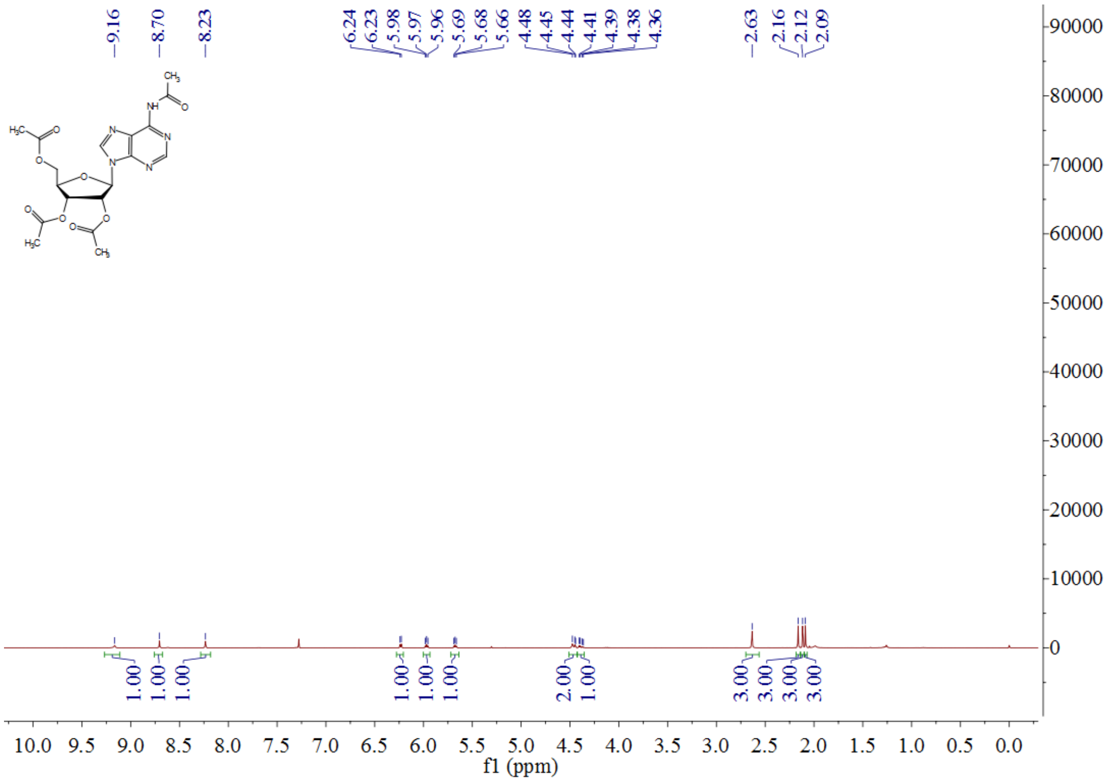

**Figure S2**. **[1]H NMR** spectra of compound (**B1**) in $CDCl_3$, 400 MHz.

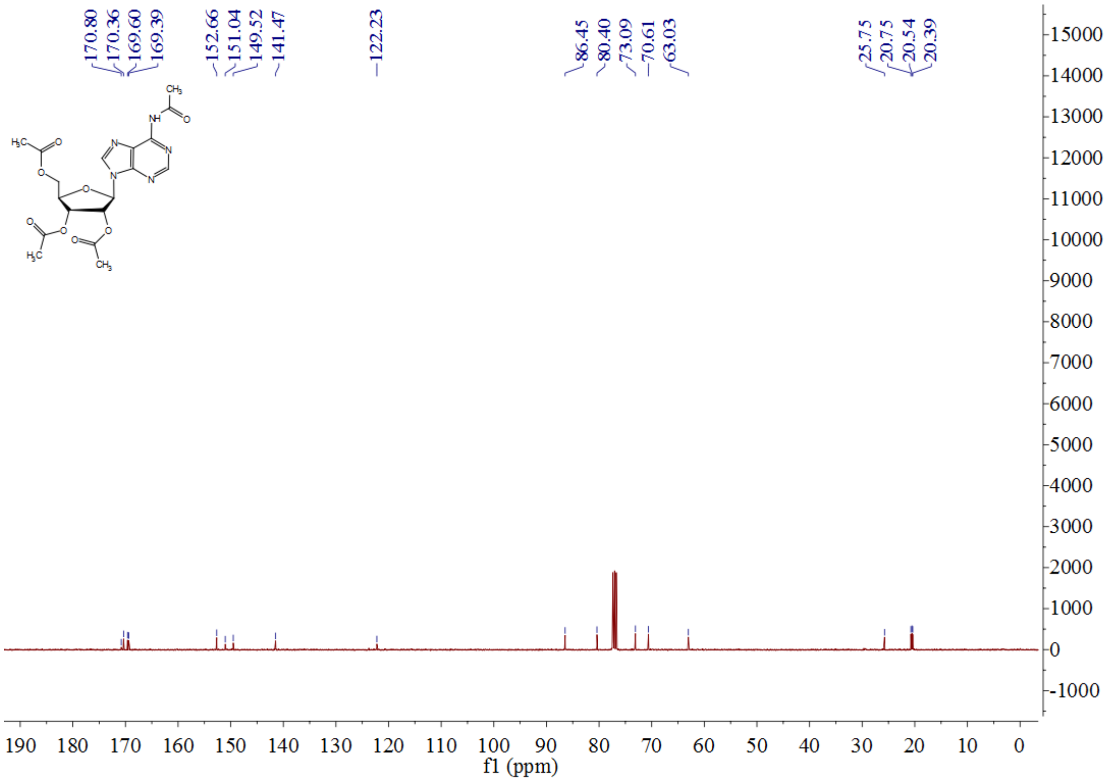

**Figure S3**. **[13]C NMR** spectra of compound (**B1**) in $CDCl_3$, 101 MHz.

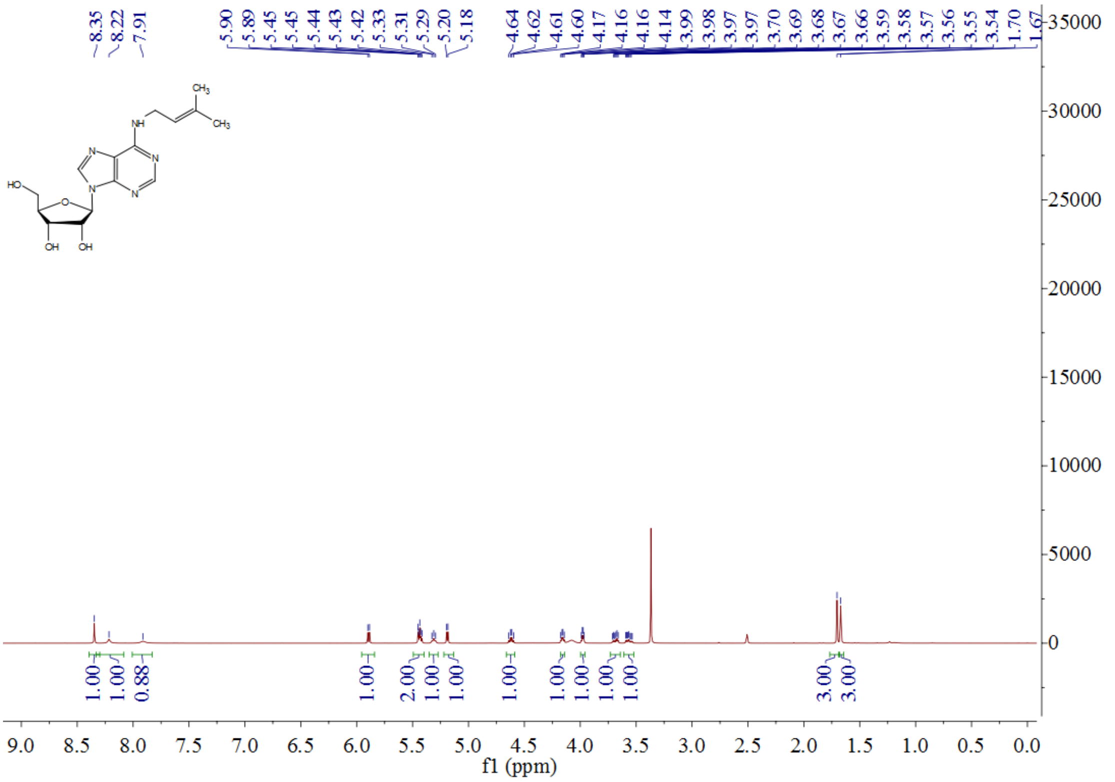

**Figure S4.** $^{1}$**H NMR** spectra of i$^{6}$A (**4**) in DMSO-$d_6$, 400 MHz.

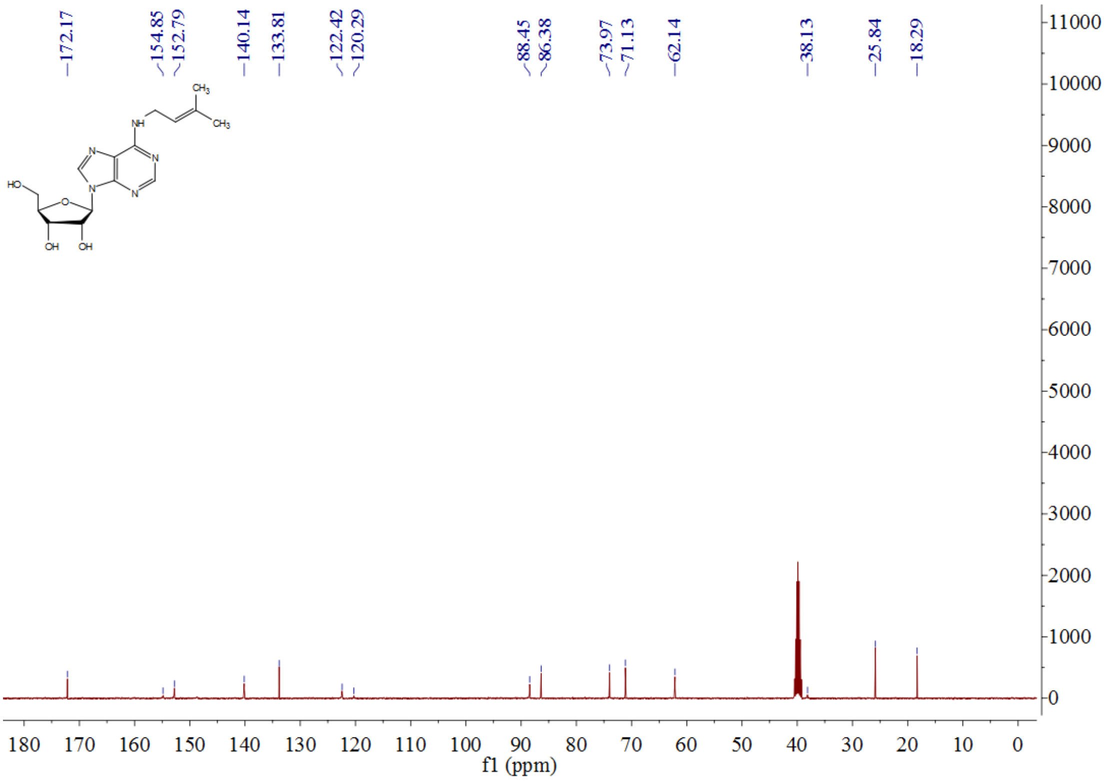

**Figure S5.** $^{13}$**C NMR** spectra of i$^{6}$A (**4**) in DMSO-$d_6$, 101 MHz.

### 4. Synthesis of fluorescent probes.

#### Probe PTAD-DBCO-FITC (8).

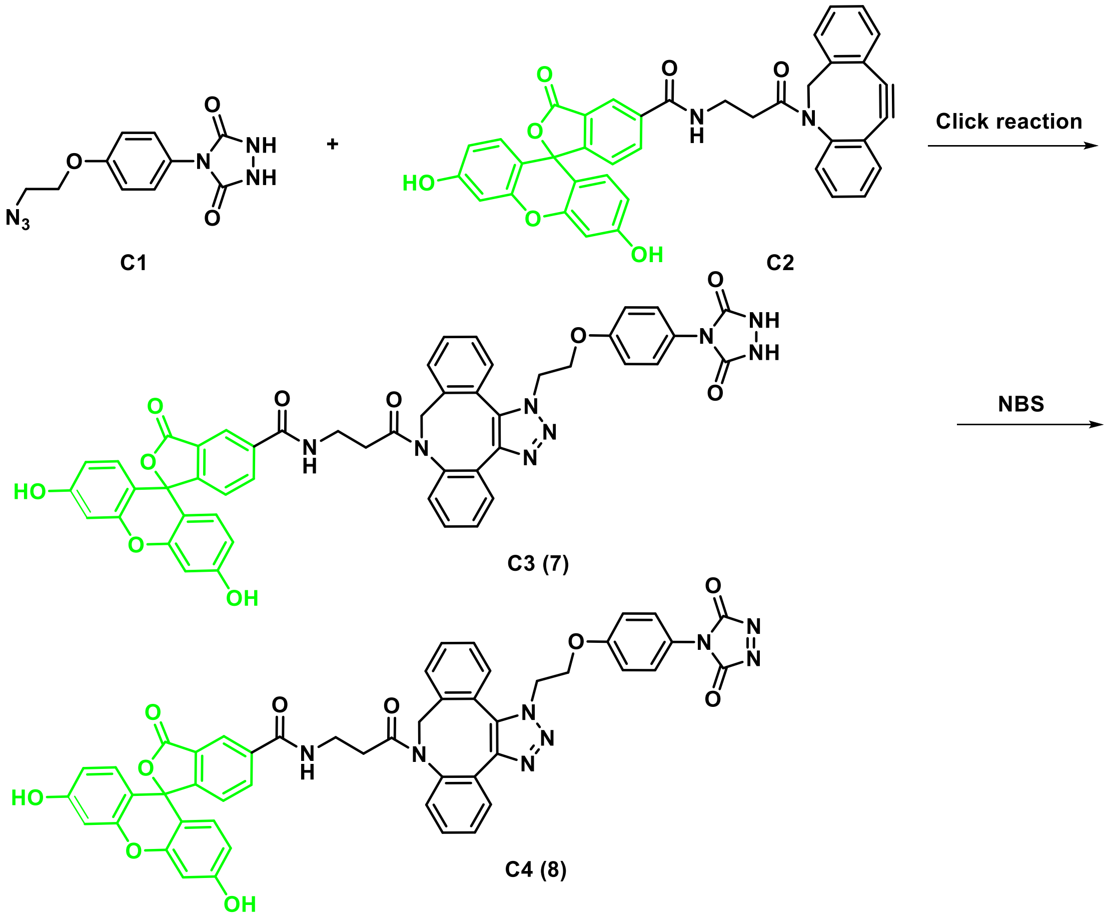

**Scheme S2.** Synthetic route of **PTAD-DBCO-FITC (8)**.

A solution of compound **C1** (10 mg, 0.04 mmol) and **FITC-DBCO** (**C2**, 5 mg, 0.01 mmol) in dry DMF (1.0 mL) under nitrogen was stirred at room temperature overnight. The reaction mixture was purified by RP-HPLC (separation was achieved using an Ultimate XB-$C_{18}$ column, 21.2 × 250 mm 5 micron (Welch Materials *Inc.*, Shanghai, China) by gradient elution from 5% to 70% acetonitrile in water (constant 0.1% formic acid) over 15 min, isocratic elution with 70% acetonitrile from 15 to 30 minutes, and returned to initial conditions and equilibrated for 5 minutes to give **PTAD-DBCO-FITC** (**7**, 1.0 mg, 0.001 mmol, 11%).

**$^1$H NMR** (400 MHz, methanol-$d^4$) δ 8.23 (s, 1H), 7.99 (d, $J$ = 8.0 Hz, 1H), 7.54 (d, $J$ = 8.0 Hz, 2H), 7.45 (s, 2H), 7.29 (d, $J$ = 4.0 Hz, 1H), 7.25-7.23 (m, 3H), 7.29 (t, $J$ = 8.0 Hz, 2H),

7.15 (d, *J* = 8.0 Hz, 1H), 6.95 (d, *J* = 8.0 Hz, 1H), 6.86 (d, *J* = 8.0 Hz, 1H), 6.60 (s, 2H), 6.51 (q, *J* = 12.0 Hz, 2H), 6.46-6.42 (m, 2H), 5.94 (d, *J* = 20.0 Hz, 1H), 4.42 (t, *J* = 16.0 Hz, 1H), 4.36 (t, *J* = 8.0 Hz, 1H), 3.40-3.30 (m, 2H), 2.24-2.08 (m, 2H), 1.96-1.74 (m, 2H). **MS (ESI)** Calculated for $[M+H]^+$ = 897.3, found 897.2. **HRMS (TOF)** Calculated $[M+H]^+$ = 897.2633, found $[M+H]^+$ = 897.2616; **HRMS (TOF)** Calculated $[M+Na]^+$ = 919.2452, found 919.2436. **PTAD-DBCO-FITC (8)** was obtained by *in-situ* oxidation using NBS (DMF solution) at room temperature for 5 minutes.

**Probe PTAD-DBCO-Cy5 (10).**

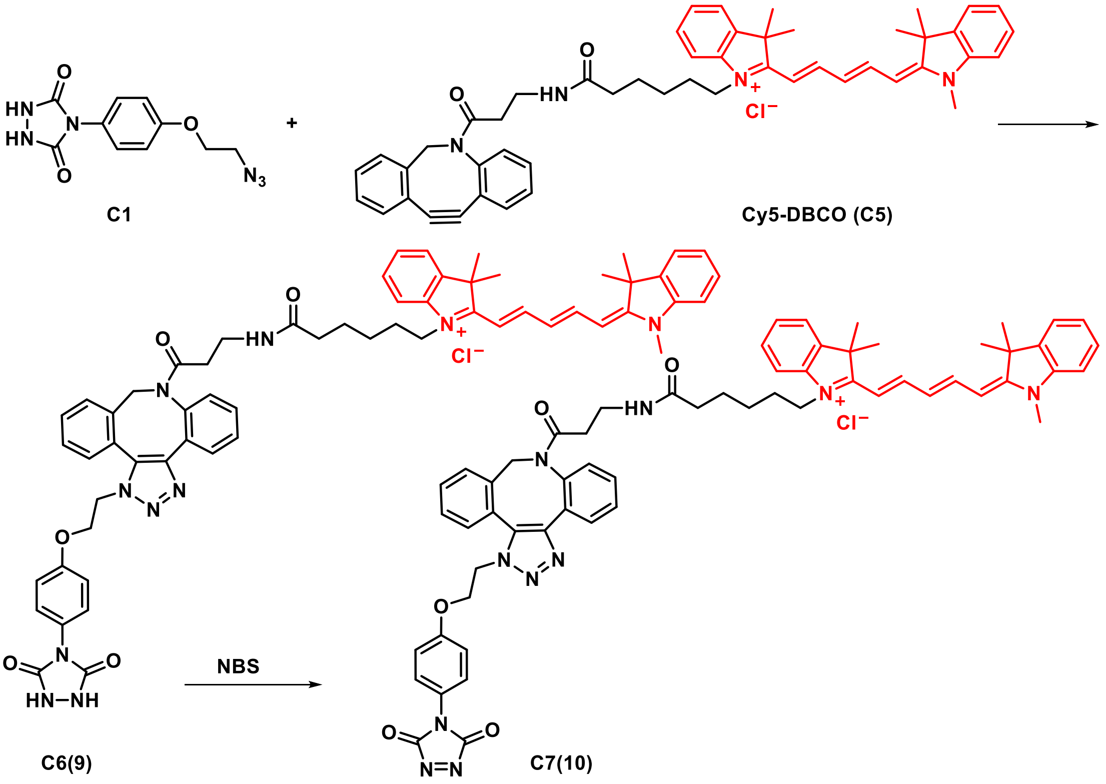

**Scheme S3.** Synthetic route of **PTAD-DBCO-Cy5** (**10**).

A solution of compound **C1** (1.7 mg, 0.0064 mmol) and **Cy5-DBCO** (**C5**, 2.5 mg, 0.0032 mmol) in dry $CH_3CN$ (1.0 mL) under nitrogen was stirred at room temperature overnight. The reaction mixture was purified by RP-HPLC (Separation was achieved using a Ultimate XB-$C_{18}$ column, 21.2 x 250 mm 5 micron (Welch Materials *Inc.*, Shanghai, China) by gradient elution from 5% to 70% acetonitrile in water (constant 0.1% formic acid) over 15 minutes, isocratic elution with 70% acetonitrile from 15 to

30 minutes, and returned to initial conditions and equilibrated for 5 minutes) to give compound **9** (3.0 mg, 0.0028 mmol, 90%) as blue solid, which was dissolved in anhydrous DMF (0.14 mL) and was treated equal equivalent *N*-bromo succinimide (NBS) solution (20 mM in DMF) to prepare **PTAD-DBCO-Cy5** stock solution (**10**, 10 mM in DMF, stored in -20 °C). **MS (ESI)** Calculated for $[M-H]^-$ = 1035.5, found 1035.8. **HRMS (TOF)** for **PTAD-DBCO-Cy5** precursor (**9**), Calculated for $[C_{60}H_{63}N_{10}O_5]^+$ = 1003.4977 as $[M-Cl]^+$ (**10**), found 1003.4964.

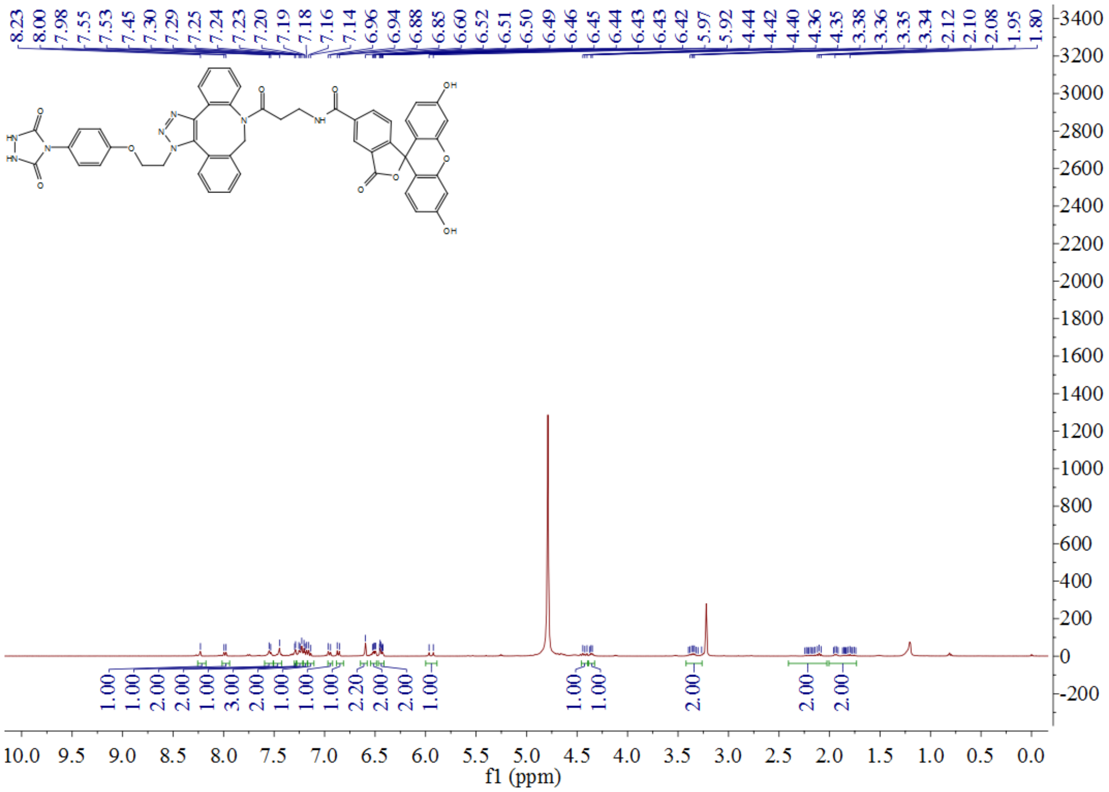

**Figure S6. $^1$H NMR** spectra of **PTAD-DBCO-FITC** precursor (**8**).

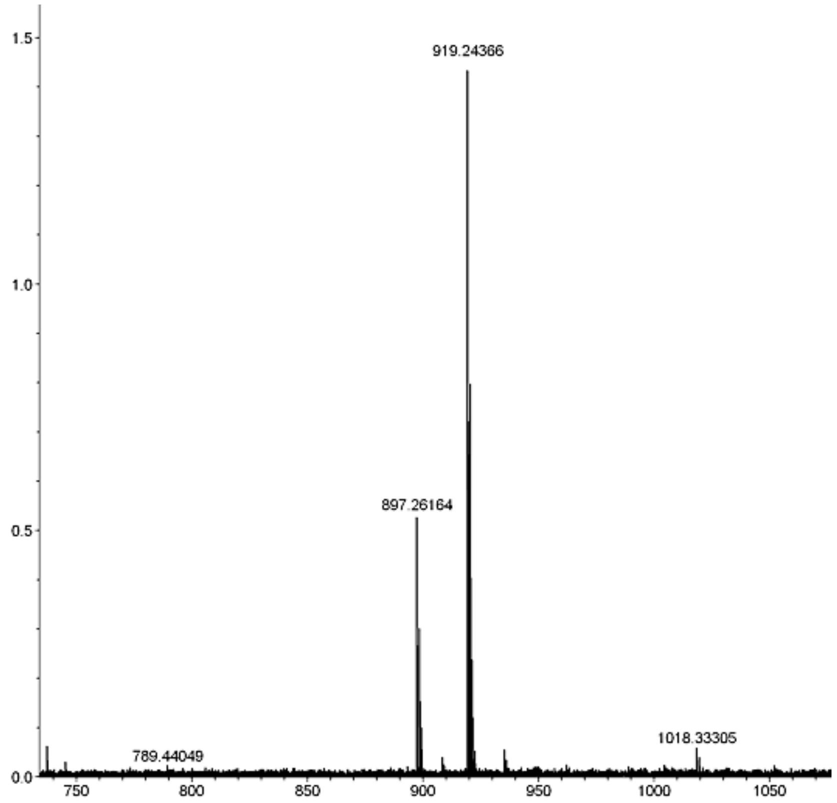

**Figure S7. HRMS (TOF)** spectra of **PTAD-DBCO-FITC** precursor (**8**).

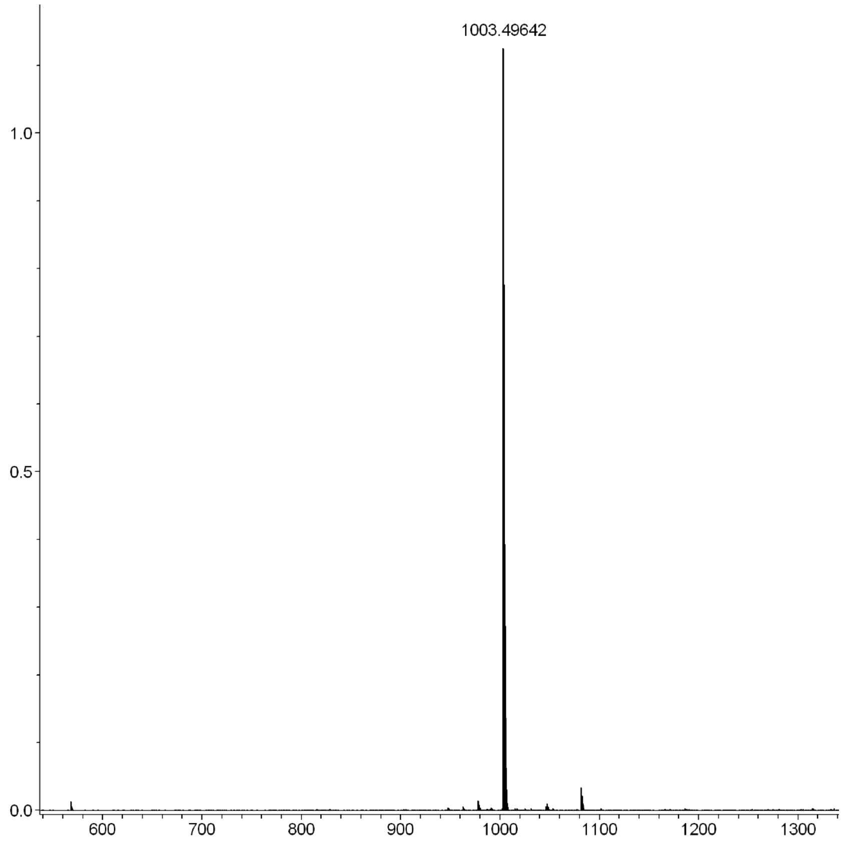

**Figure S8. HRMS (TOF)** spectra of **PTAD-DBCO-FITC** precursor (**10**).

### 5. Investigations of reaction properties of i[6]A (4) with PTAD (5).

#### Reaction of i[6]A (4) with PTAD (5).

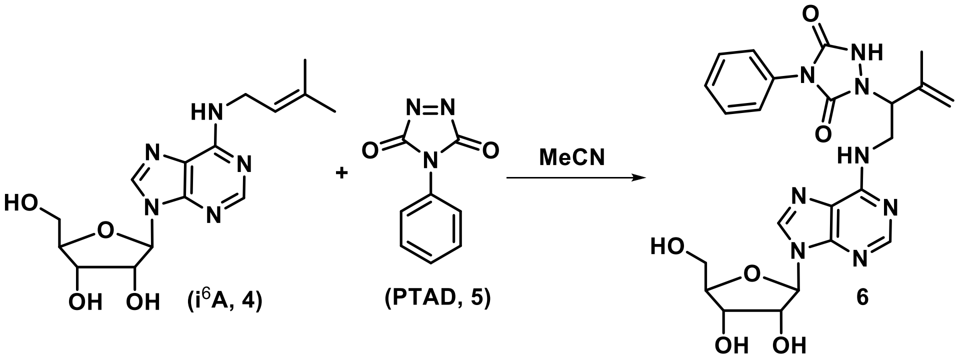

To a solution of nucleoside i[6]A (**4**, 33 mg, 0.1 mmol) in acetonitrile (1.0 mL) was added 4-phenyl-1, 2, 4-triazoline-3, 5-dione (**5**, PTAD, 34 mg, 0.2 mmol) and stirred at room temperature until the red color of PTAD disappeared. Then the organic solvents were removed. The residue was purified by silica gel column chromatography ($CH_2Cl_2$: MeOH = 10: 1) to afford adduct **6** (20 mg, 0.039 mmol, 39%) as white solid.

**$^1$H NMR** (400 MHz, methanol-*d*$^4$) δ 8.26 (t, *J* = 12.0 Hz, 2H), 7.51 (d, *J* = 4.0 Hz, 1H), 7.42-7.33 (m, 4H), 7.24-7.19 (m, 2H), 5.95 (q, *J* = 8.0 Hz, 1H), 5.18 (d, *J* = 12.0 Hz, 2H), 5.07 (t, *J* = 16.0 Hz, 1H), 4.76-4.70 (m, 1H), 4.32 (q, *J* = 8.0 Hz, 1H), 4.18-4.11 (m, 3H), 3.88 (q, *J* = 16.0 Hz, 1H), 3.76-3.72 (m, 1H), 1.90 (s, 3H). **$^{13}$C NMR** (151 MHz, $CD_3OD$) δ 154.92, 153.72, 152.02, 140.41, 131.48, 128.75, 128.59, 128.58, 127.86, 126.03, 124.44, 120.04, 113.87, 89.95 (d, *J* = 21.1 Hz), 86.73, 74.06 (d, *J* = 27.2 Hz), 71.29 (d, *J* = 18.1 Hz), 62.10 (d, *J* = 15.1 Hz), 59.06, 48.18, 20.15 (d, *J* = 4.5 Hz). **MS (ESI)** Calculated for $[M+Na]^+$ = 533.2, found = 533.0. **HRMS (TOF)** Calculated for $C_{23}H_{27}N_8O_6$ $[M+H]^+$ = 511.2054, found 511.2041.

#### Dynamic investigations of i[6]A (4) with PTAD (5).

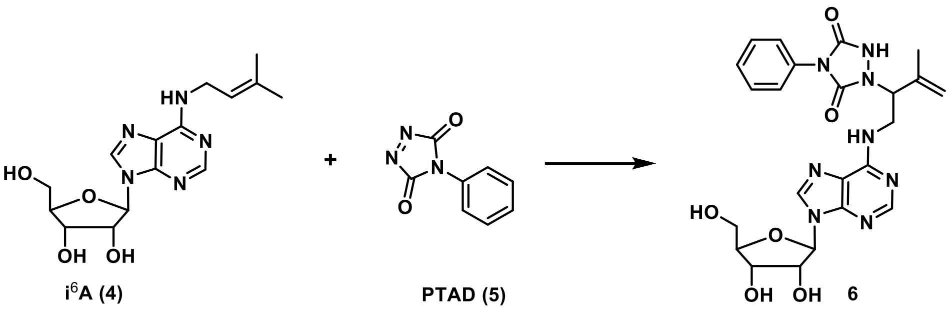

Nucleoside i$^6$A (**4**, 10 mg, 0.03 mmol) were dissolved in $CD_3CN$/ $D_2O$ (1.0 mL, v/ v = 1: 1, final concentration = 30 mM). PTAD (**5**, 26 mg, 0.15 mmol) was then added to the solution (final concentration = 150 mM), and the reaction mixture was allowed to stand at room temperature. The **$^1$H NMR** spectra were recorded by proton NMR (400 MHz).

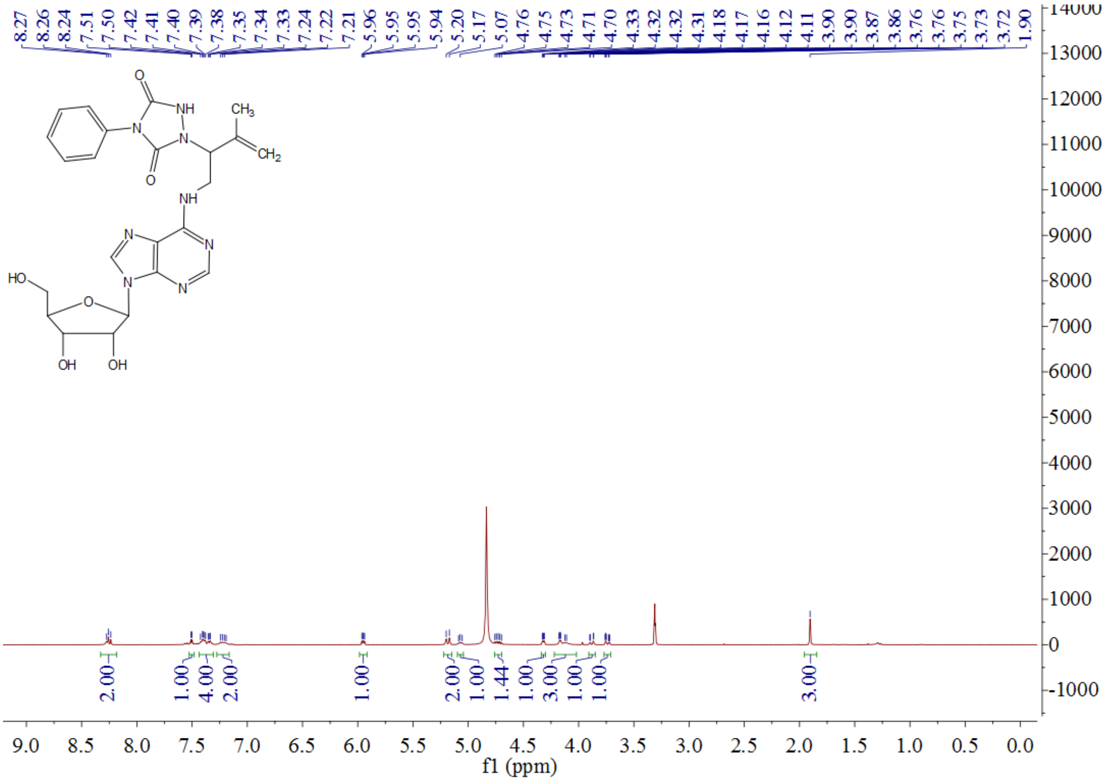

**Figure S9. $^1$H NMR** spectra of compound (**6**).

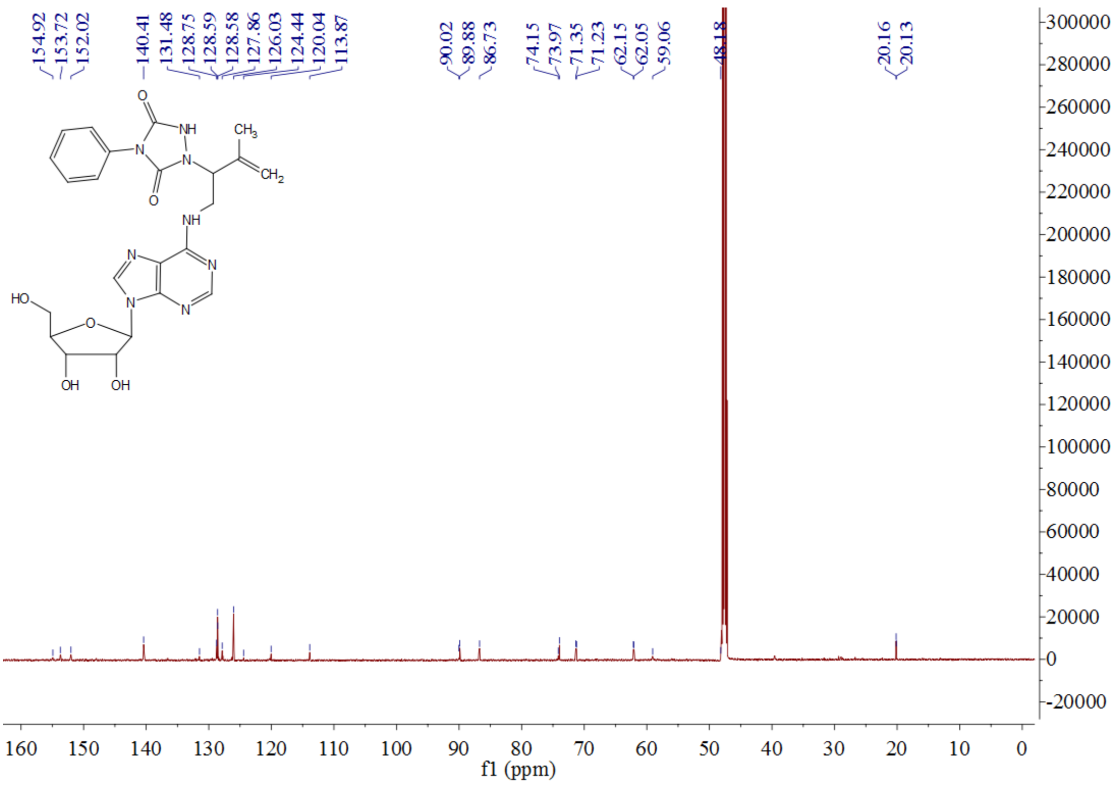

**Figure S10.** **$^{13}$C NMR** spectra of the adduct **(6).**

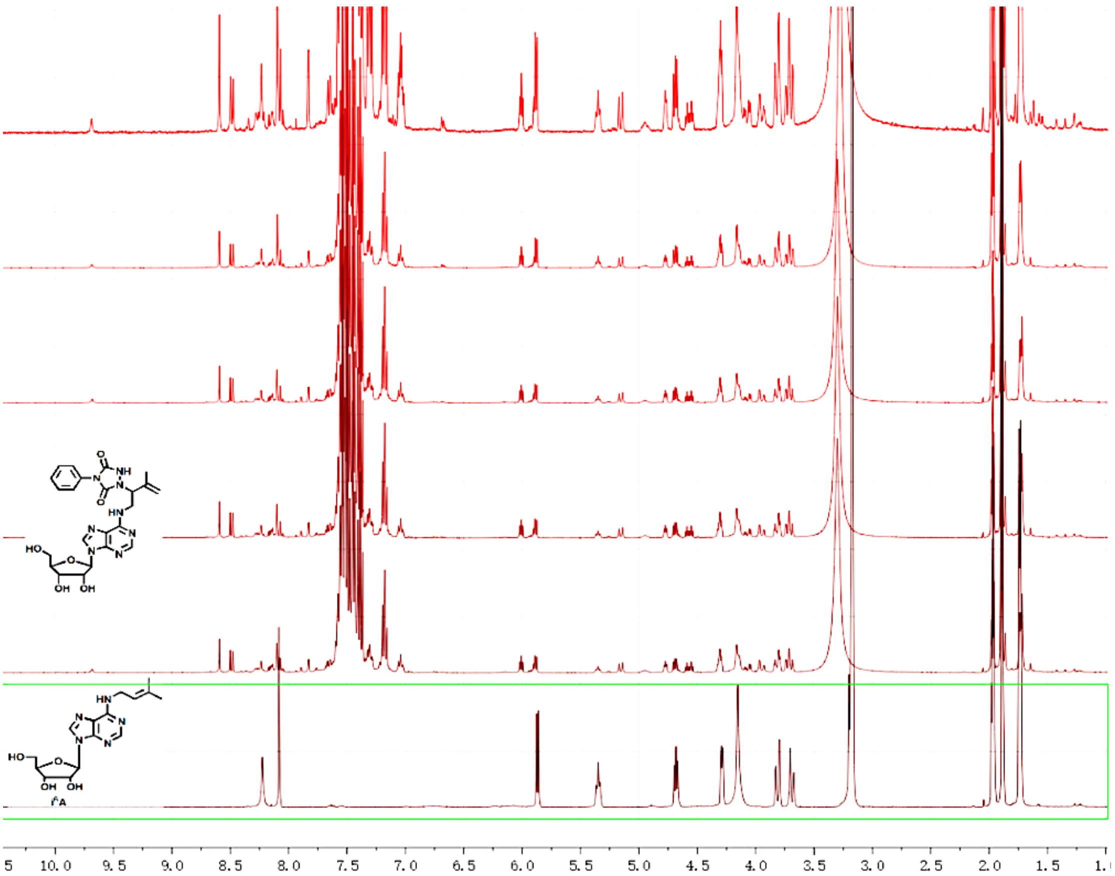

**Figure S11.** Dynamic investigations of the reaction of i$^6$A (**4**) with PTAD (**5**) by **$^1$H NMR**.

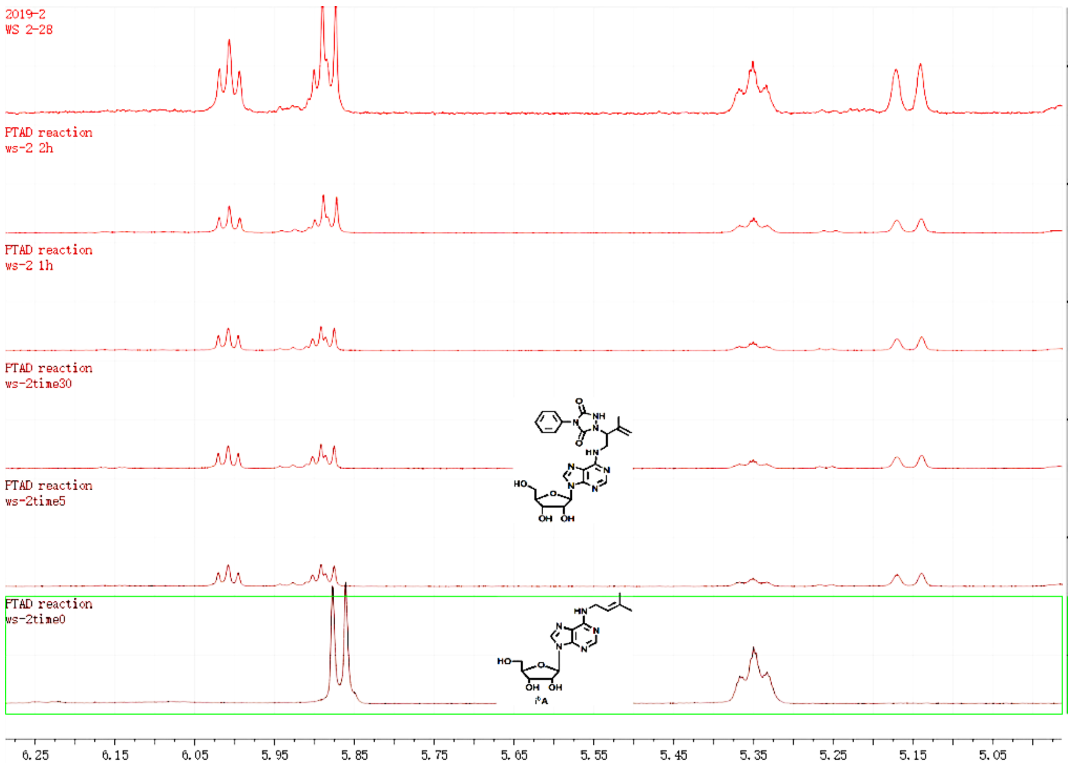

**Figure S12.** Investigations dynamic property of the reaction of i[6]A (**4**) with PTAD (**5**). The reaction was run in an NMR tube. A solution of i[6]A (**4**, 20 mg) was dissolved in DMSO-$d^6$ (0.6 mL). To this solution was added PTAD (**5**, 20 mg), the reaction was monitored by **$^1$H NMR**. **$^1$H NMR** shown that the i[6]A (**4**) was almost consumed in 60 minutes.

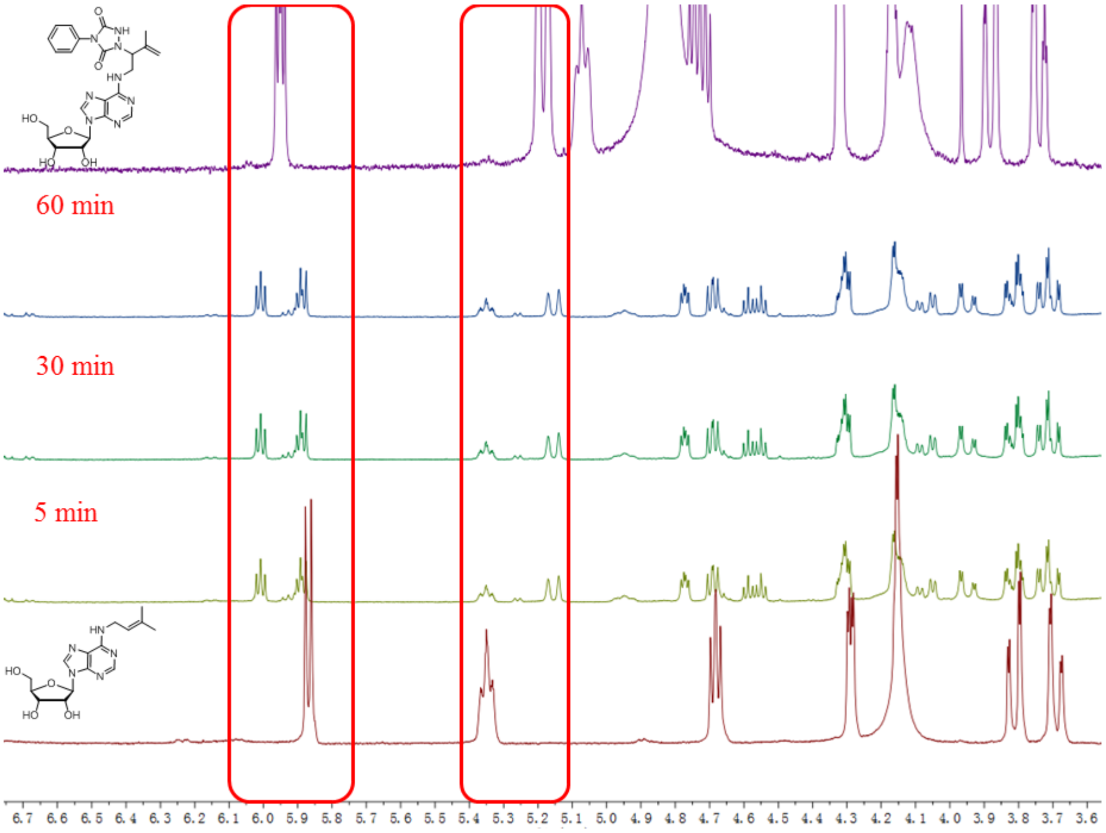

**Figure S13.** The investigations of i[6]A (**4**) with PTAD (**5**) via **$^1$H NMR** experiment.

### 6. Synthesis of nucleotide i$^6$ATP.

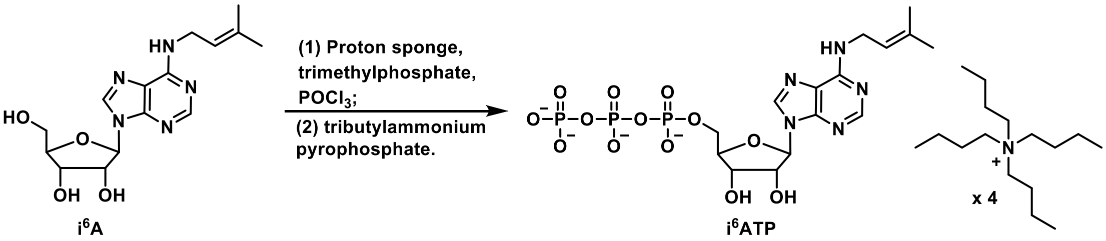

**Scheme S4.** Synthetic route of i$^6$A **(4)** triphosphate.

Nucleoside i$^6$A (**4**, 35 mg, 0.1 mmol) and proton sponge (58 mg, 0.25 mmol) were dissolved in trimethyl phosphate (0.7 mL) and was placed under the ice-water bath. Phosphorus oxychloride ($POCl_3$, 33.6 μL, 0.36 mmol) was slowly added and stirred at 0 °C for 12 hours. A solution of tributylamine (175 μL, 0.72 mmol) and tributyl ammonium pyrophosphate (642 mg) in DMF (2.0 mL) was slowly added and the reaction mixture was stirred at 0 °C for 30 minutes. The reaction was quenched by addition of 1.0 M aqueous triethylammonium bicarbonate (TEAB, pH = 7.5, 15 mL). The mixture was diluted with $H_2O$ (5.0 mL) and subjected to HPLC purification. Separation was achieved using an Ultimate XB-$C_{18}$ column, 21.2 × 250 mm 5 micron by gradient elution from 5% to 50% acetonitrile in water (constant 0.1% formic acid) over 25 minutes, isocratic elution with 50% acetonitrile from 25 to 30 minutes, and returned to initial conditions and equilibrated for 5 minutes to give i$^6$ATP (10.0 mg, as tributyl ammonium salt).

**$^1$H NMR** (400 MHz, methanol-$d^4$) δ 8.40 (s, 2H), 8.30 (s, 1H), 6.13 (d, $J$ = 4.0 Hz, 1H), 5.40 (t, $J$ = 12.0 Hz, 2H), 4.73 (q, $J$ = 16.0 Hz, 1H), 4.57-4.43 (m, 1H), 4.27-4.26 (m, 2H), 4.20-4.13 (m, 2H). **$^{31}$P NMR** (162 MHz, methanol-$d^4$) δ -10.47 (d, $J$ = 22.68 Hz, α-P), -11.46 (d, $J$ = 22.68 Hz, γ-P), -24.00 (t, $J$ = 22.68 Hz, β-P). **MS(ESI)** Calculated for $[M-H]^-$ = 574.1, found 574.2.

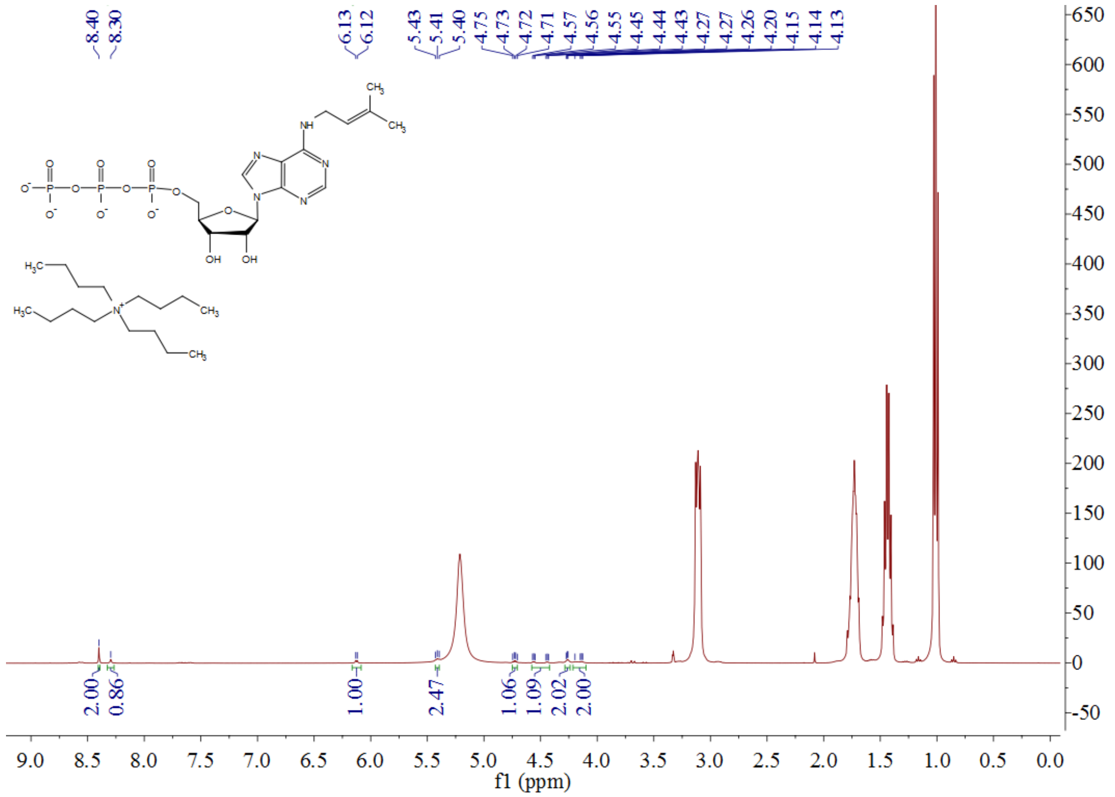

**Figure S14.** **$^{1}$H NMR** spectra of i$^{6}$ATP in $CD_3OD$, 400 MHz.

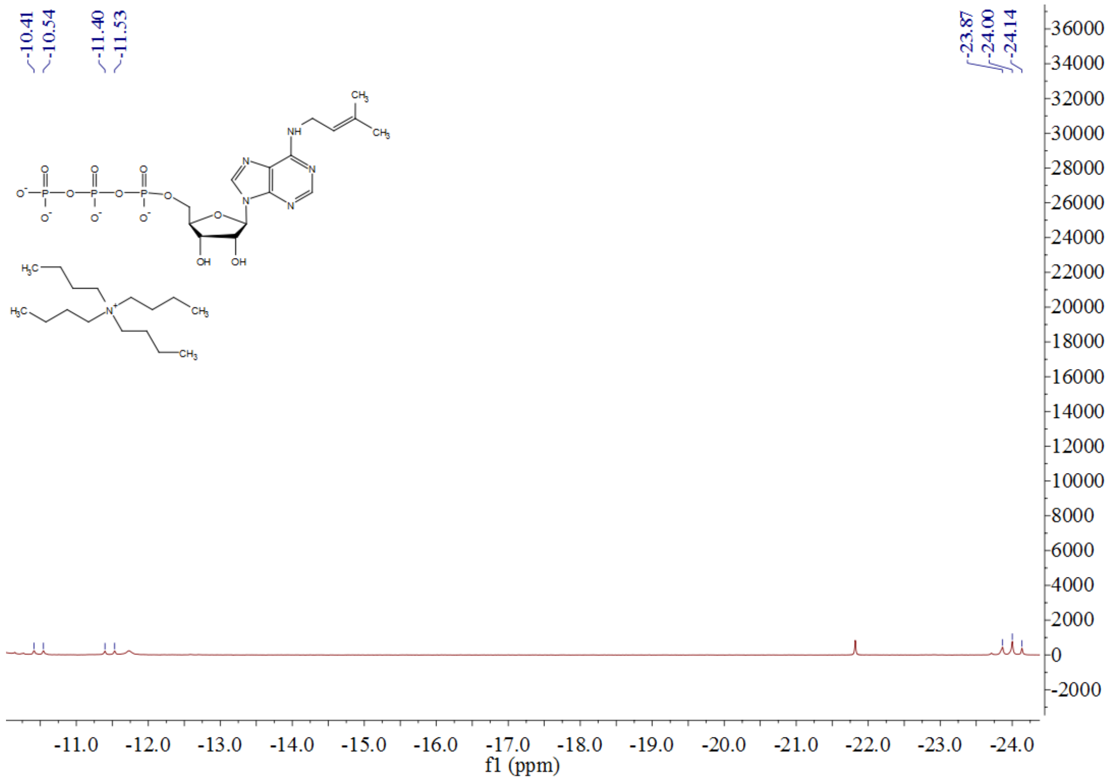

**Figure S15.** **$^{31}$P NMR** spectra of i$^{6}$ATP in $CD_3OD$, 162 MHz.

### 7. Profiling of i$^6$A-incoporated RNA.

#### 7.1 Test the ability of T7 RNA polymerase to recognize modified nucleotides (i$^6$ATP) at specific sites.

***Step 1**. Construction of T7 RNA polymerase-mediated EGFP transcription system in vitro.*

In view of the high specificity of *T7 RNA polymerase*, T7 promoter as a strong promoter can efficiently guide the expression of downstream genes. Here, the T7 promoter sequence is used to guide the transcription of *EGFP in vitro*.

T7 promoter EGFP Gene

5'-TAATACGACTCACTATAGGGATGGTGAGCAAGGGCGAGGA....TGGACGAGCTGTACAAG -3'

**Figure S16.** Illustration of the construction of transcription system of chimeric *T7P-EGFP* gene *in vitro*.

*EGFP* gene sequence was amplified from *pEGFP-C1* plasmids. Then T7 promoter sequence was linked to the 5'-end by fusion PCR. The primers are summarized as follow:

| Name | Sequence (5'-3') |
|---|---|
| T7P-GFP-F | TAATACGACTCACTATAGGGATGGTGAGCAAGGGCGAGGAGCTG |
| T7P-GFP-R | CTACTTGTACAGCTCGTCCATGCCG |

**Table S1.** Sequences for the construction of chimeric *T7P-EGFP*.

***Step 2**. Detection of T7 RNA polymerase catalyzed RNA polymerization process in vitro.*

Using the *T7P-EGFP* constructed above as a template and rNTP as substrates, the reaction was carried out at 37 $^oC$ for 2 hours which was catalyzed by T7 RNA polymerase. After the reaction, the template DNA *T7P-EGFP* was degraded with DNase I. Finally, the RNA was precipitated and purified, and the transcription effect of the RNA was detected by agarose gel electrophoresis. The *T7P-EGFP* template sequence and HeLa genomic RNA were used as the control group.

The results show that through the above *T7 RNA polymerase* transcription system *in vitro*, the transcription of the target gene has been successfully achieved.

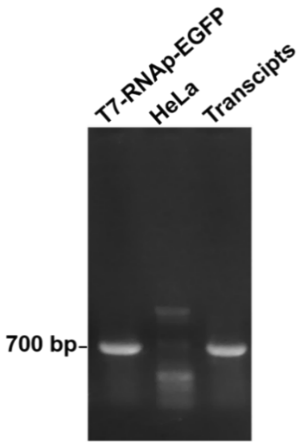

**Figure S17.** Detection of the transcripts of *T7P-EGFP*. Lane 1: *T7P-EGFP* template. Lane 2: HeLa genome. *T7-RNAp-EGFP* represents *T7P-EGFP* herein.

Firstly, the *EGFP* gene sequence containing the T7 promoter was obtained by PCR amplification, and then the *T7P-EGFP* gene sequence was used as a template to perform *in vitro* reverse transcription by *T7 RNA polymerase*. The HeLa cell genome was used as a control template, and after the transcription was completed, it was detected by agarose gel electrophoresis. Based on the above assays, it has been demonstrated that using *T7P-EGFP* as a template the *in vitro* transcription of *EGFP* RNA sequence.

**7.2 Optimization of the labeling conditions and labeling ability detection of T7 RNA polymerase-promoted specific RNA synthesis using i$^6$ATP as the substrate.**

Preliminarily experiments have unveiled that the recognition efficiency of *T7 RNA polymerase* for i$^6$ATP is significantly higher than that of K4. Next, i$^6$ATP will be used as a substrate to optimize the labeling conditions when i$^6$ATP is incorporated on the target RNA using the *T7 RNA polymerase* transcription system, and further test the labeling efficiency.

***Step 1.*** *Detection of fidelity of chimeric T7P-EGFP towards i$^6$ATP substrate.*

EGFP-RNA was transcript with T7 transcription kit (*Takara*) *in vitro*. The system was kept at 37 °C for 2 hours. 1-4 extended to 6 hours. Then the templets were digested with *DNase I*, 37 °C for 30 minutes. After that, the RNA was precipitated with 75% alcohol at -20 °C overnight. Then the RNA was washed with 75% alcohol and re-dissolved with RNase free DEPC water (20 μl) and detected by 1% agarose gel electrophoresis. RNA concentration was determined with a Nanodrop One microvolume UV-Vis. spectrophotometer.

| **System** | **Volume** |
|---|---|
| *T7P-EGFP* gene | 1 μg |
| T7 RNA polymease | 1 μl |
| Transcription buffer in *vitro* | 2 μl |
| Lane 2 (rNTP mixture, 100 mM) | 2 μl |
| Lane 3 (A\C\G\UTP, 100 mM) | 0.5 μl × 4 |
| Line 4 ($i^6$ATP, 100 mM) | 0.5 μl× 1 |
| Line 5 ($i^6$ATP, 100 mM) | 0.5 μl × 2 |
| Line 6 ($i^6$ATP, 100 mM) | 0.5 μl × 3 |
| Line 7 ($i^6$ATP, 100 mM) | 0.5 μl × 4 |
| DEPC water | total 20 μl |

**Table S2.** Protocol of transcription system *in vitro*.

As depicted, it is demonstrated that *T7 RNA polymerase* could recognize $i^6$ATP, although the recognition efficiency is low comparing with rATP.

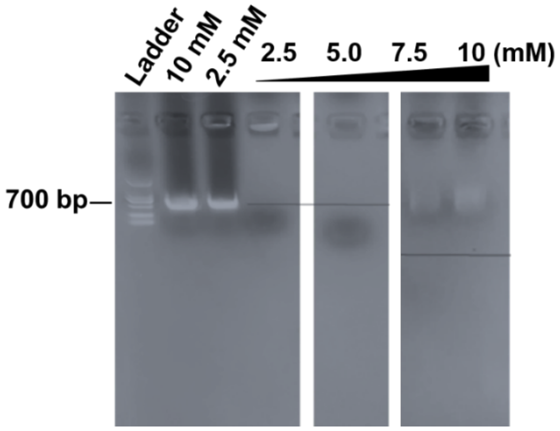

**Figure S18.** Transcription system in vitro. Lane 1: ladder. Lane 2-3 were positive control assays. Lane 2 (rNTP mixtures, 2.0 μl, 100 mM, separately). Lane 3 (A/C/G/UTP, 0.5 μl, 100 mM, separately). Lane 5-8 (C/G/UTP, 0.5 μl, 100 mM, separately) and varied concentrations of $i^6$ATP (0.5, 1.0, 1.5 and 2.0 μl, 100 mM). 1% agarose gel electrophoresis was used to monitor the transcripts.

The construction systems are shown below:

| System | Volume |
|---|---|
| *T7P-EGFP* gene | 1.0 μg |
| T7 RNA polymerase | 1.0 μl |
| Transcription buffer *in vitro* | 2.0 μl |
| NTP (100 mM) | 2.0 μl |
| $i^6$ATP (100 mM) | 4.0 μl |
| DEPC water | total 20.0 μl |

**Table S3.** Protocol for transcription.

Then the templets were digested with *DNase I*, 37 °C for 30 minutes. Subsequently, the RNA was precipitated with 75% alcohol at -20 °C overnight. Then the RNA was washed with 75% alcohol and re-dissolved with DEPC water (20 μl). The RNA will be directly used for the next step.

***Step 2.*** *Optimization of the transcription efficiency with $i^6$ATP.*

In order to improve the transcription efficiency of T7P-*EGFP* under the condition of $i^6$ATP addition, the above conditions were optimized.

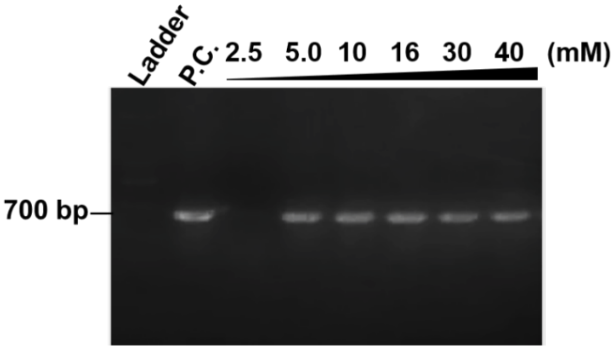

**Figure S19.** Optimization of the transcription efficiency in vitro. P. C. represents positive control assay, rNTP mixtures are A/C/U/GTP (0.5 μl, 100 mM, separately). All experiments were used mixtures with rNTP (A/C/U/GTP, 0.5 μl, 100 mM, separately) and varied concentrations of $i^6$ATP: 0.5, 1.0, 2.0, 4.0, 6.0, 8.0 μl (100 mM). Transcripts were run via 1% agarose gel electrophoresis.

The results showed that the transcription efficiency of *T7P-EGFP* is low in the presence of low concentration $i^6$ATP in experimental group 1, which is consistent with the previous results. When the same concentration of adenosine triphosphate (ATP) is added, the transcription efficiency of *T7P-EGFP* is significantly improved, and the transcription efficiency of *T7P-EGFP* decreases with the increase of the concentration of i$^6$ATP, but it is not obvious.

**7.3 Detection of the labeling efficiency using optimized condition with i$^6$ATP targeting mRNA of T7P-EGFP.**

Through the above experiments, it has been demonstrated that i$^6$ATP can be recognized by *T7 RNA polymerase*, and subsequently the *in vitro* transcription efficiency of *EGFP* mRNA in the presence of i$^6$ATP has been optimized. Next, the efficiency of i$^6$ATP transcription into *EGFP* needs to be further verified, and the prenyl group on i$^6$A (**4**) is specifically labeled by the well-designed fluorescent probe **PTAD-DBCO-FITC** (**8**).

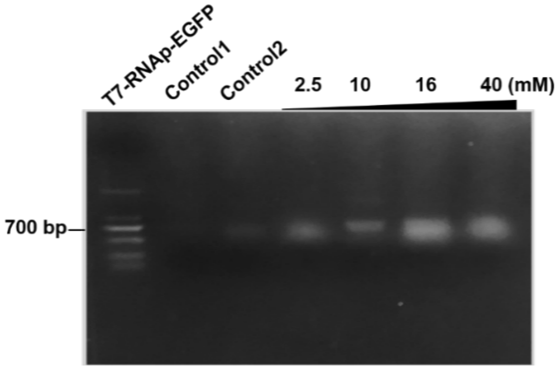

**Figure S20**. Optimization of the transcription efficiency *in vitro*. P. C. represents Positive Control assay, rNTP mixtures are A/C/U/GTP (0.5 µl, 100 mM, separately). All experiments were used mixtures with rNTP (A/C/U/GTP, 0.5 µl, 100 mM, separately) and varied concentrations of i$^6$ATP: 0.5, 1.0, 2.0, 4.0, 6.0, 8.0 µl (100 mM). Transcripts were run *via* 1% agarose gel electrophoresis. *Control1* is a negative control without template, and Control2 is a negative control without substrate. *T7-RNAp-EGFP* represents *T7P-EGFP* herein.

The experimental results showed that the control group could not detect the fluorescence signal of **PTAD-DBCO-FITC** (**8**), and with the increase of i$^6$ATP concentration, the fluorescence intensity of RNA band gradually increased, which indicated that the content of i$^6$A (**4**) in RNA gradually increased. The labeling efficiency of RNA gradually increases.

**7.4 Construction of T7-RNA polymerase eukaryotic expression system.**

In order to realize the detection of RNA labeling efficiency by use of i$^6$A (**4**) in eukaryotic cells, the eukaryotic expression system of *T7 RNA polymerase* was first constructed. By detecting the insertion level of $i^6A$ (**4**) in the downstream target gene mRNA guided by the T7 promoter, the target RNA can be labeled by the addition of $i^6A$ (**4**) in eukaryotic cells. If T7 RNA polymerase in eukaryotic cells can recognize $i^6A$ (**4**), the change in the transcription profile of the cell at a specific time point can be detected by controlling the detection time when $i^6A$ (**4**) is added.

At first stage, detection of the recognition of $i^6A$ (**4**) by T7 RNA polymerase in the eukaryotic cells and the RNA labeling efficiency.

***Step 1.*** *Construction map of T7-RNA polymerase eukaryotic expression system.*

In order to realize the simultaneous expression of *T7 RNA polymerase* and its target gene in the same cells, the *T7 RNA polymerase* and *EGFP* gene expression system were constructed into the same eukaryotic expression system. As shown below, *T7 RNA polymerase* is transcribed and expressed by the strong eukaryotic promoter CMV, and the *EGFP* gene sequence is guided by the T7 promoter. In addition, because T7 RNA polymerase transcription produces mRNA lacking the ribosome recognition sequence in eukaryotic cells, the translation process cannot be realized in eukaryotic cells. Therefore, the 5'-end of the *EGFP* gene is connected to an *IRES2* sequence, and the *IRES2* transcription sequence can guide the ribosome to downstream genes. Translation process to realize the expression process of *EGFP* in eukaryotic cells such as tumor cells. In addition, in order to detect the expression of *EGFP* and T7 RNA polymerase and the relationship between the two, respectively, an expression system for *EGFP* and T7 RNA polymerase was constructed.

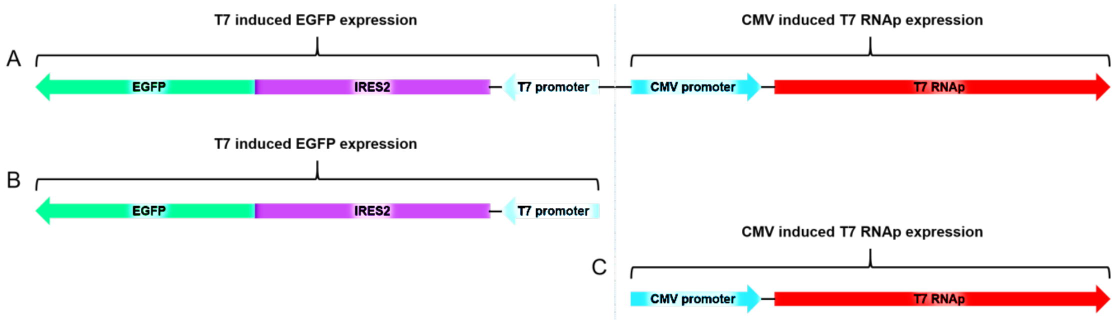

**Figure S21.** The structure of the intracellular transcription system catalyzed by T7 RNA polymerase. *Top A*, the T7 RNA polymerase transcription system guided by the CMV promoter and the *EGFP*

transcription system guided by the T7 promoter are constructed in the same vector. *Bottom B and C*, the T7 RNA polymerase transcription system guided by the CMV promoter and the *EGFP* transcription system guided by the T7 promoter were constructed into two transfection systems respectively.

***Step 2.*** *Detection of expression of T7 RNA polymerase and EGFP in HeLa cells.*

Next, the T7 RNA polymerase constructed above and the expression level of the *EGFP* eukaryotic expression system mediated by T7 RNA polymerase in tumor cells were further tested. To this end, the above system is transfected into HeLa cells through a cation-mediated transfection reagent. After 36 hours of transfection, the expression of green fluorescent protein was first detected by fluorescence microscope. Then the cells were lysed by RIPA, the total cell protein was extracted, and the expression of T7 RNA polymerase and *EGFP* were detected by Western-blotting.

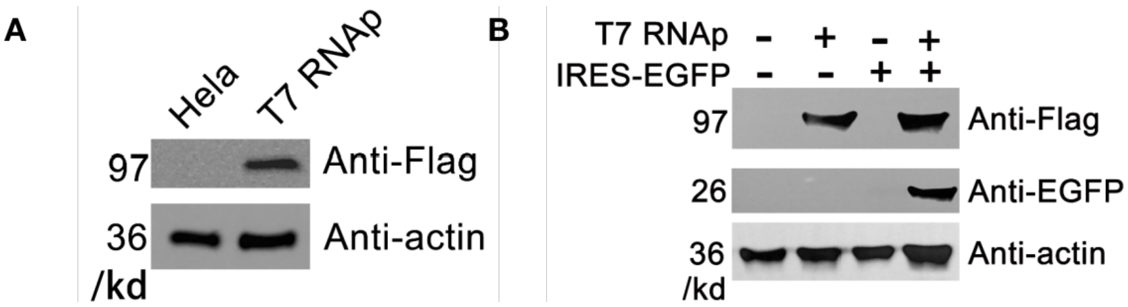

**Figure S22**. Western-blotting detection the protein expression of T7 RNA polymerase and *EGFP*. **Fig. S22A**. Transfect the *CMV-T7 RNA polymerase* transcription system into HeLa cells, and detection of the protein expression of *T7 RNA polymerase* after 36 hours. The T7 RNA polymerase has a FLAG tag. **Fig. S22B**. HeLa cells were transfected with *CMV-T7 RNA polymerase* and *T7* RNA polymerase-*IRES2-EGFP* transcription system, and *CMV-T7 RNA polymerase* and *T7 RNA polymerase-IRES2-EGFP* combined system, respectively. The expression of *EGFP* was detected 36 hours later. *T7 RNAp* represents *T7 RNA polymerase.*

The results showed that *T7 RNA polymerase* successfully achieved expression in HeLa cells. Moreover, the expression of *EGFP* is *T7 RNA polymerase* dependent (**Fig. S22A**), and *EGFP* expression can only be detected in cells that also express *T7 RNA polymerase* (**Fig. S22B**). Therefore, the above experiments show that the *EGFP* eukaryotic expression system mediated by *T7 RNA polymerase* has been successfully constructed.

### 7.5 Detection of T7 RNA polymerase transcription in eukaryotic cells by fluorescent expression of EGFP.

In the above experiment, the protein expression of *T7 RNA polymerase* and *EGFP* has been detected by Western-blotting. Next, the expression of *EGFP* was further detected by fluorescence microscope. As in the previous step, the *CMV-T7 RNA polymerase* and *T7 RNA polymerase-IRES2-EGFP* transcription systems were transfected into HeLa cells, and the two systems were transfected together. In addition, the *CMV-T7 RNA polymerase* and *T7 RNA polymerase-IRES2-EGFP* integration systems were separately transfected into an experimental group. 36 hours after transfection, the expression of *EGFP* fluorescent protein was detected by fluorescence microscope. The experimental results are consistent with the western-blot detection results, that is, the expression of *EGFP* is T7 polymerase-dependent. No matter in the *CMV-T7 RNA polymerase* and *T7 RNA polymerase-IRES2-EGFP* single transfection group or the two integrations group, only two systems existed concurrently, the expression of green fluorescent protein can only be detected.

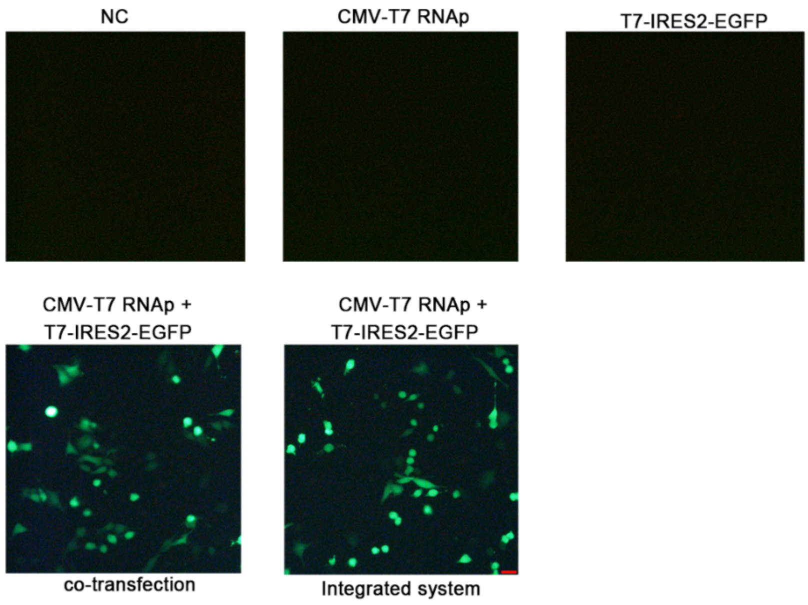

**Figure S23**. Detection of *T7 RNA polymerase* transcription system catalyzing the transcription and expression of *EGFP* fluorescent protein in HeLa cells. The *CMV-T7 RNA polymerase* and *T7 RNA polymerase-IRES2-EGFP* systems were transfected into HeLa cells, and the two systems were transfected together and the integrated system was transfected. After 36 hours of the transfection, the expression of *EGFP* green fluorescent protein was detected under a fluorescence microscope. Scale bar 50 μM. *T7 RNAp* represents T7 RNA polymerase herein. Scale bar: 50 um.

The above results demonstrated that the system constructed herein has successfully achieved the *T7 RNA polymerase* specifically directing the transcription of target genes in eukaryotic cells.

#### 7.6 Detection of i$^6$A-incoporated RNA labeling ability in eukaryotic cells.

Through the above experiments, we have successfully constructed a T7 RNA polymeras*e* recognition system for i$^6$A (**4**), and realized that i$^6$A (**4**) participates in the transcription of the target gene. In addition, a T7 RNA polymerase-guided eukaryotic transcription system was constructed to realize the eukaryotic transcription process of target genes guided by *T7 RNA polymerase*. Next, we will further test whether i$^6$A (**4**) can be recognized and participate in the transcription process in eukaryotic cells.

***Step 1****. Cytotoxic assays of i$^6$A (**4**) towards HeLa cells.*

In order to better detect the involvement of i$^6$A (**4**) in the transcription process of cellular RNA, we also tested the effect of i$^6$A (**4**) on cytotoxicity and optimized the concentration of i$^6$A (**4**) to treat cells.

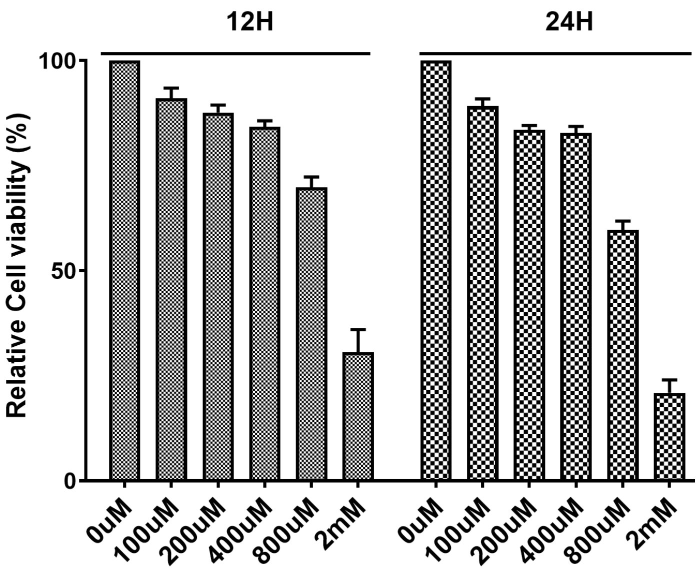

**Figure S24.** Cytotoxic assays using various concentration of i$^6$A (**4**) in 12 hours or 24 hours.

The experimental results show that the i$^6$A (**4**) concentration within 400 μM has little effect on cell viability. When the concentration reaches 800 μM, cell viability will be significantly inhibited. When the concentration reaches 2.0 mM, i$^6$A (**4**) has a significant inhibitory effect on cells.

***Step 2**. Exploration of whether i⁶A (**4**) can be recognized by RNA polymerase in eukaryotic cells.*

Next, we will further test whether $i^6A$ (**4**) can participate in the RNA transcription process in eukaryotic cells. First culture the HeLa cells in a six-well plate to 70% - 80%. Then add different concentrations of $i^6A$ (**4**) to DMEM medium (100 µM, 200 µM and 400 µM of $i^6A$). After 12 hours of culture, collect all cells and extract total RNA.

**RNA ($i^6A$-incoporated) labelled by fluorescein PTAD-DBCO-FITC (8)**.

To the 1.5 mL Eppendorf tube containing PTAD-DBCO-FITC precursor (**7**, 10 µL, 10 mM solution in DMF) was added *N*-bromo succinimide (1.0 µL, 100 mM in DMF). The mixture was vortexed gently and formation of the light-red color was observed. The reagent was kept on ice and used for the $i^6A$-incorporated RNA labelling immediately. Labeling protocol is the same stated above.

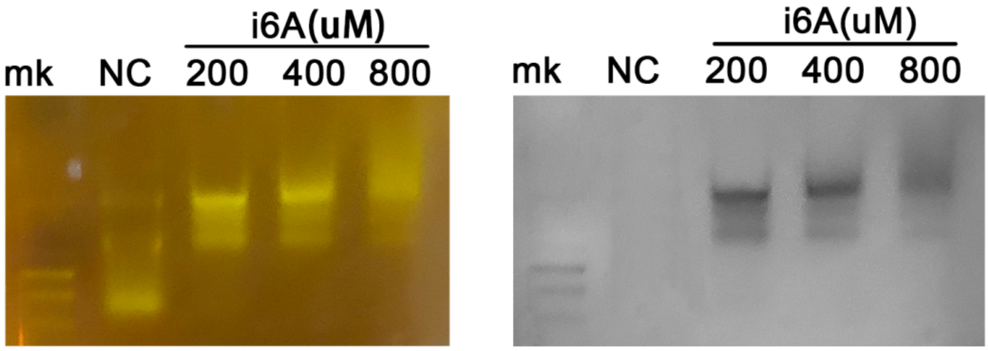

**Figure S25.** Detection of $i^6A$'s involvement in RNA transcription in eukaryotic cells. After culturing the cells with DMEM medium containing 200 µM, 400 µM, and 800 mM of $i^6A$ (**4**) in HeLa cells for 24 hours, the total RNA was extracted and then labeled with **PTAD-DBCO-FITC (8)** to detect the fluorescence intensity of **PTAD-DBCO-FITC (8)**.

After labeling process, remove excess **PTAD-DBCO-FITC (11)** dye to determine whether $i^6A$ (**4**) can be transcribed into RNA efficiently.

The analysis of the results showed that under the conditions of GoldView™ nucleic acid dye staining, it was detected that the total RNA image in the cell with addition of $i^6A$ (**4**) was better than that of the wild type. In addition, its 5s, 18s RNA bands moved up significantly compared with the wild type, while the 28s band did not change significantly.

Through **PTAD-DBCO-FITC (8)** dye staining, it was found that no fluorescence signal was detected in the control group, but significant fluorescence signal was detected in the total RNA of the cells in the presence of i$^6$A (**4**), and the addition of i$^6$A (**4**, 800 µM) resulted in a weaker RNA fluorescence signal, which may be due to i$^6$A (**4**, 800 µM) has certain cytotoxicity to cells. Through this experiment, we have demonstrated that i$^6$A (**4**) can be recognized in eukaryotic cells and participate in the RNA transcription process.

#### 7.7 Labeling of i$^6$A-incoporated RNA *in vivo*.

HeLa cells was obtained from ATCC. HeLa cells were cultured in DMED medium with 10% fetal calf serum. Cells were maintained in a humidified incubator at 5% $CO_2$ and 37 °C. To label RNA *in vivo*, HeLa cells were cultured in 6-well plates to 70%-80%. Then different concentration i$^6$A (**4**) was added to DMEM medium (i$^6$A: 100 µM, 200 µM, 400 µM). After 12 hours, all of the cells were collected and total RNA was extracted with total RNA extraction kit (*Takara*). The RNA was further labeled with **PTAD-DBCO-FITC (8)** as described. Then it will be detected by agarose gel electrophoresis.

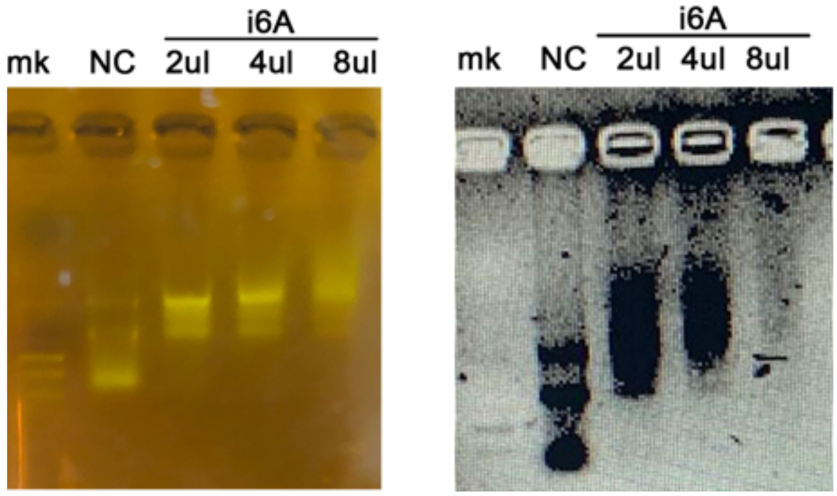

**Figure S26.** Gel electrophoresis results of the transcripts containing i$^6$A (**4**).

### 8. Procedure for labeling of RNA (i$^6$A-incoporated) using fluorescent PTAD-DBCO-Cy5 (10).

To the 1.5 mL Eppendorf tube containing PTAD-DBCO-Cy5 precursor (**9**, 10 $\mu$L, 10 mM solution in DMF) was added *N*-bromo succinimide (1.0 $\mu$L, 100 mM in DMF). The mixture was vortexed gently and formation of the light-red color was observed. The reagent was kept on ice and used for the i$^6$A-incorporated RNA labelling immediately. The i$^6$A-incoporated total RNA (10 µl, 20 ng/ µl in DEPC water) was incubated with PTAD-DBCO-Cy5 (**10**, 10 µl, 0.1 mM), shaken for 30 minutes at 0 °C. RNA was analyzed by gel electrophoresis on 1 % agarose. The electrophoresis conditions were as follows: power supply (DYY-6C power supply, Liuyi Biotechnology) was set to 130 V. 1% agarose gels were run at room temperature (25 °C) for 20 minutes and stained with or without GoodView™ (SBS Genentech, Beijing, China). A DNA size marker (1 kb DNA Ladder, TsingKe Biotech, TSJ102, Beijing, China) was used. Gels were analyzed by in-gel fluorescence measurements on a FluorChem® FC3 imager (Alpha Innotech). Fluorescence was measured with a blue light excitation wavelength (475 nm) and a green filter emission (537 nm).

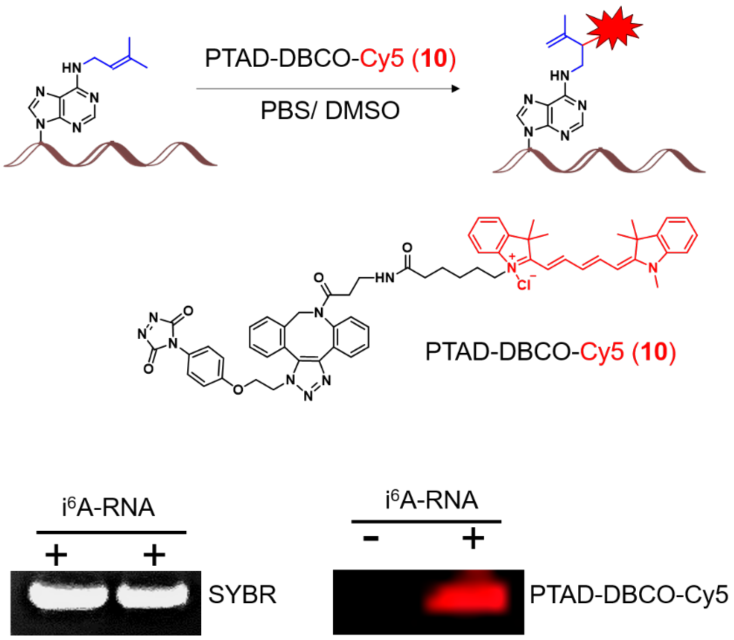

**Figure S27**. Directly fluorescent labeling of prenylated tRNA from *E. coli* using PTAD-DBCO-Cy5 (**10**).

### 9. Investigation of $I_2$-mediated cycloaddition of i⁶A (4).

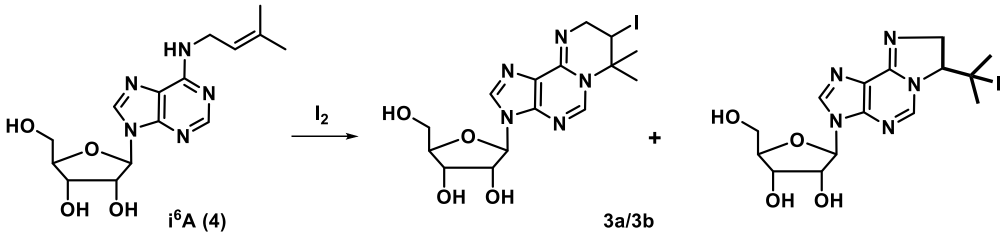

Model reactions of $i^6A$ nucleoside (**4**). The model reactions were carried out in DMSO-$d^6$ in order to facilitate *in-situ* NMR characterization without further purification. $i^6A$ (**4**, 0.15 mmol) was dissolved in DMSO-$d^6$ (0.5 mL), and iodine (0.45 mmol) was added. The mixture was stirred at 37 °C for 5 minutes and was taken out for **$^1$H NMR**.

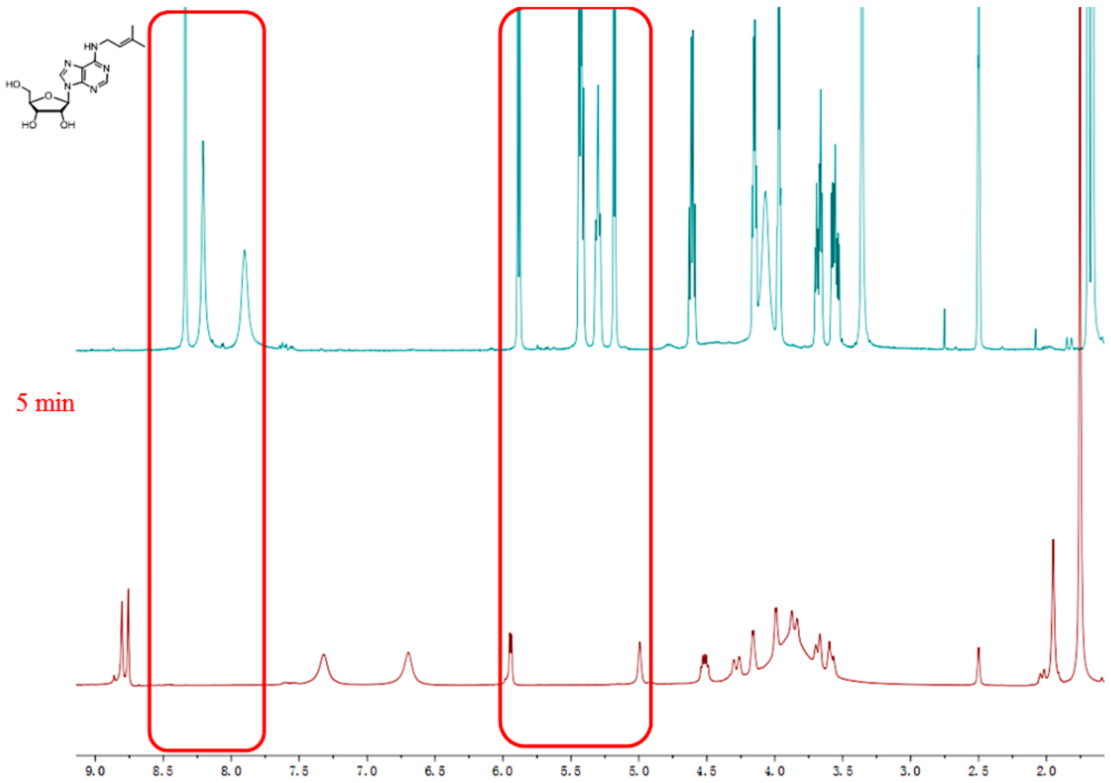

**Figure S28. $^1$H NMR** comparison of $i^6A$ (**4**) and $i^6A$ (**4**) with $I_2$.

#### 9.1 HPLC and mass investigations.

The nucleoside $i^6A$ (**4**, 0.15 mmol) was dissolved in DMSO (1.0 mL), and iodine (0.45 mmol) was added. The mixture was stirred at 37 °C for 1 hour. Afterwards, saturated $Na_2S_2O_3$ in $H_2O$ was titrated into the solution in order to remove the excess iodine, and then $Na_2CO_3$ in $H_2O$ was added. The resultant mixture was further stirred at 37 °C for 1 hour and then subjected to HPLC. Separation was achieved using an Ultimate XB-$C_{18}$ column, 4.6 × 150 mm 5 micron (Welch Materials *Inc.,* Shanghai, China) by gradient elution at 0.5 mL/ min from 5% to 90% acetonitrile in water (constant 0.1% formic acid) over 15 minutes, isocratic elution with 90% acetonitrile from 15 to 18 minutes, and returned to initial conditions and equilibrated for 5 minutes).

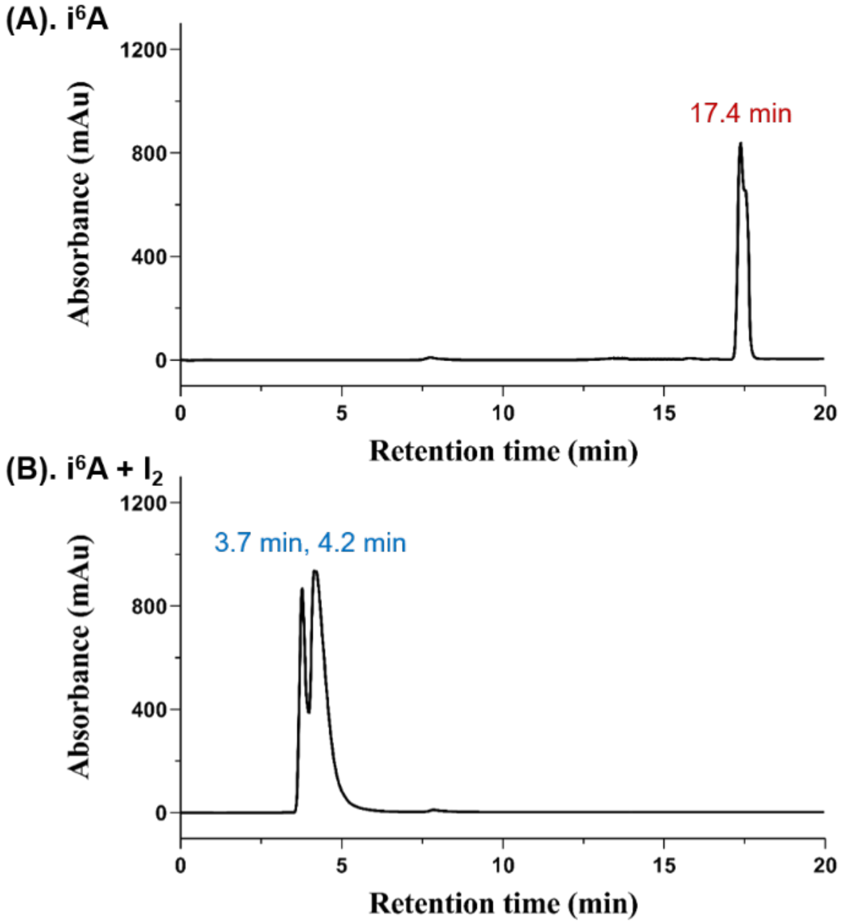

**Figure S29.** HPLC comparison of $i^6A$ (**4**) and the adduct (**3a/ 3b**) of $i^6A$ (**4**) with PTAD (**5**).

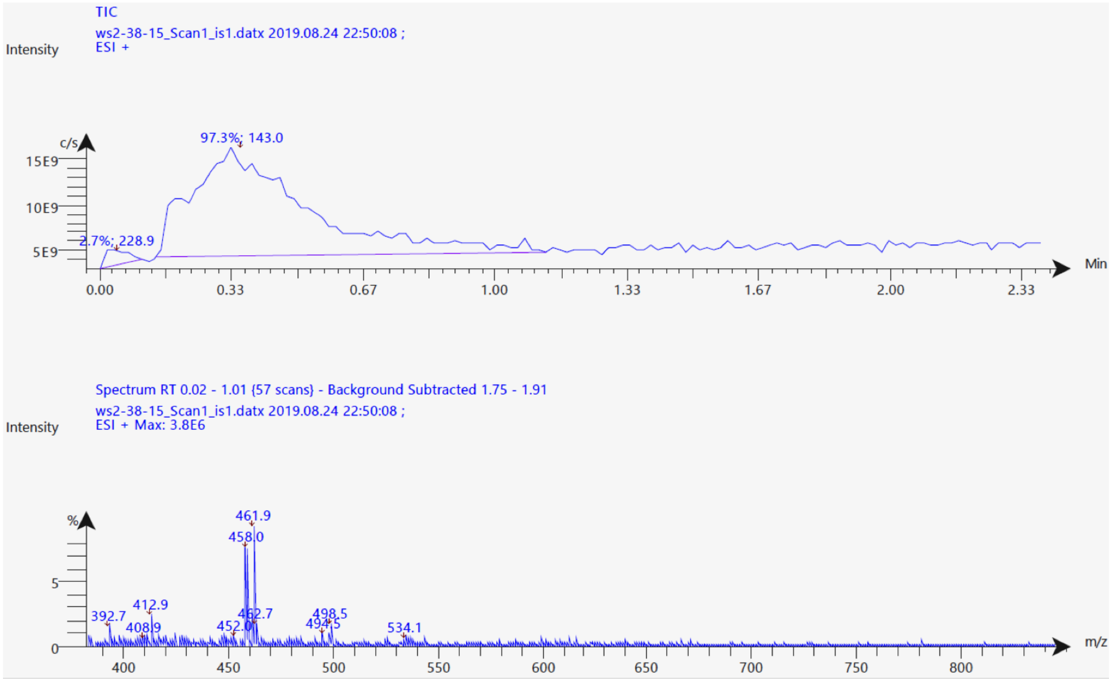

**Figure S30.** HPLC comparison of i$^6$A (**4**) and the adduct (**3a/ 3b**) of i$^6$A (**4**) with PTAD (**5**). **MS (ESI)** $[M+H]^+$ = 461.9.

Acquisition Paramet

| | | | | | |
|---|---|---|---|---|---|
| Source Type | ESI | Ion Polarity | Positive | Set Nebulizer | 0.3 Bar |
| Focus | Not active | | | Set Dry Heater | 200 °C |
| Scan Begin | 50 m/z | Set Capillary | 4500 V | Set Dry Gas | 4.0 l/min |
| Scan End | 3000 m/z | Set End Plate Offset | -500 V | Set Divert Valve | Waste |

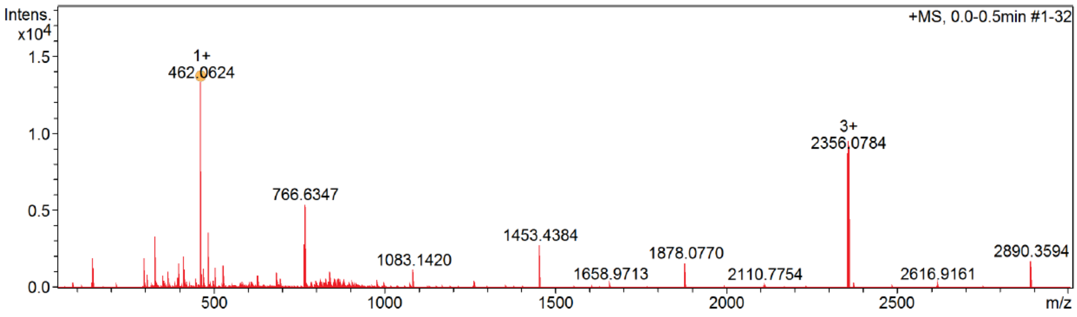

| Meas. m/z | # | Ion Formula | m/z | err [ppm] | Mean err [ppm] | rdb | N-Rule | e¡¥ Conf | mSigma | Std I | Std Mean m/z | Std I VarNorm | Std m/z Diff | Std Comb Dev |
|---|---|---|---|---|---|---|---|---|---|---|---|---|---|---|
| 462.062358 | 1 | C15H21IN5O4 | 462.063279 | 2.0 | -0.3 | 7.5 | ok | even | 1.4 | 2.0 | n.a. | n.a. | n.a. | n.a. |

**Figure S31. HRMS (TOF)** Calculated for $[M+H]^+$= 462.0638, found 462.0624.

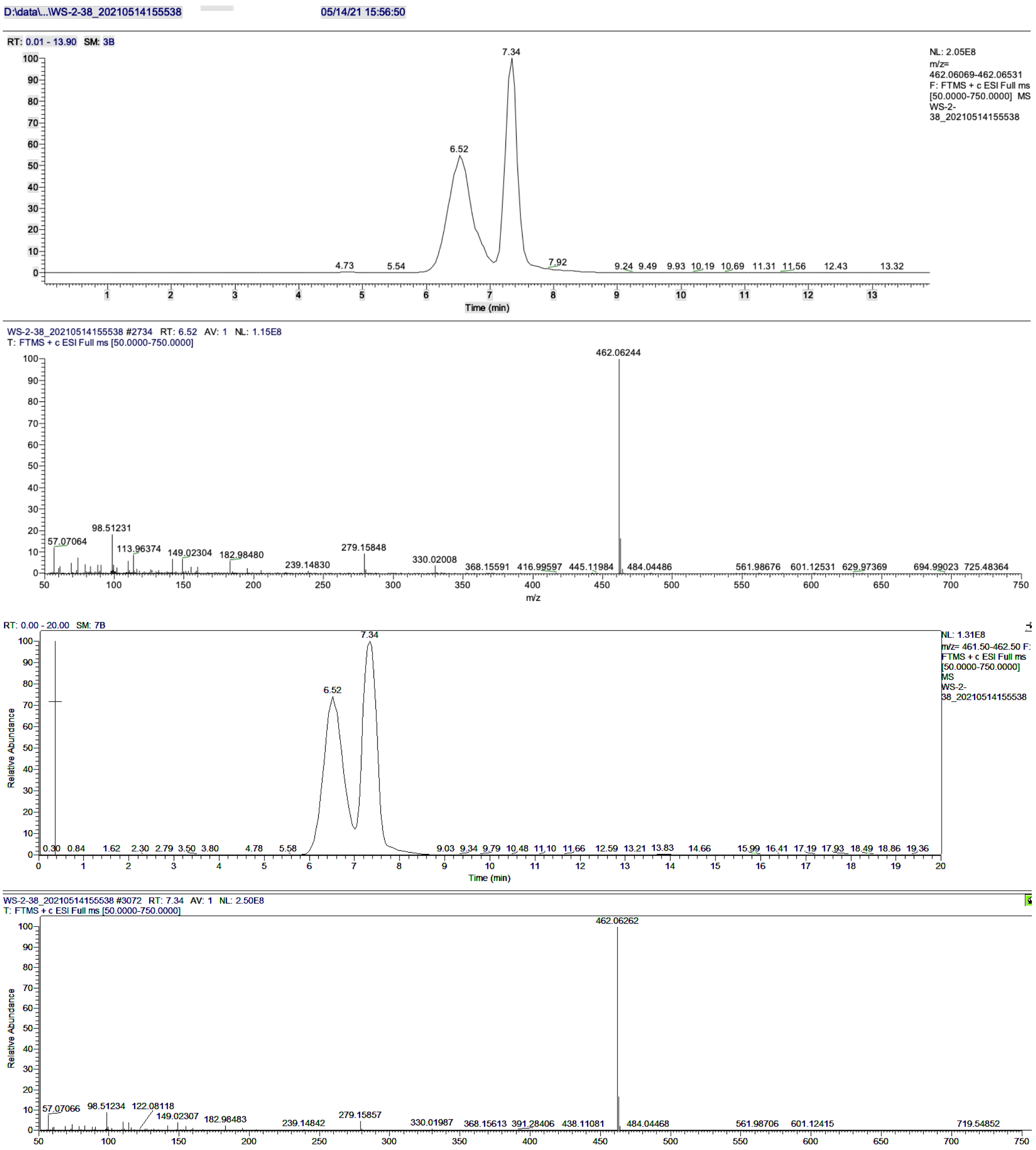

**Figure S32.** LC-MS analysis of cyclo adduct (**3a/ 3b**). $[M+H]^+$ = 462.06244 (6.52 min) or 462.06262 (7.34 min).

### 9.2 NMR investigation.

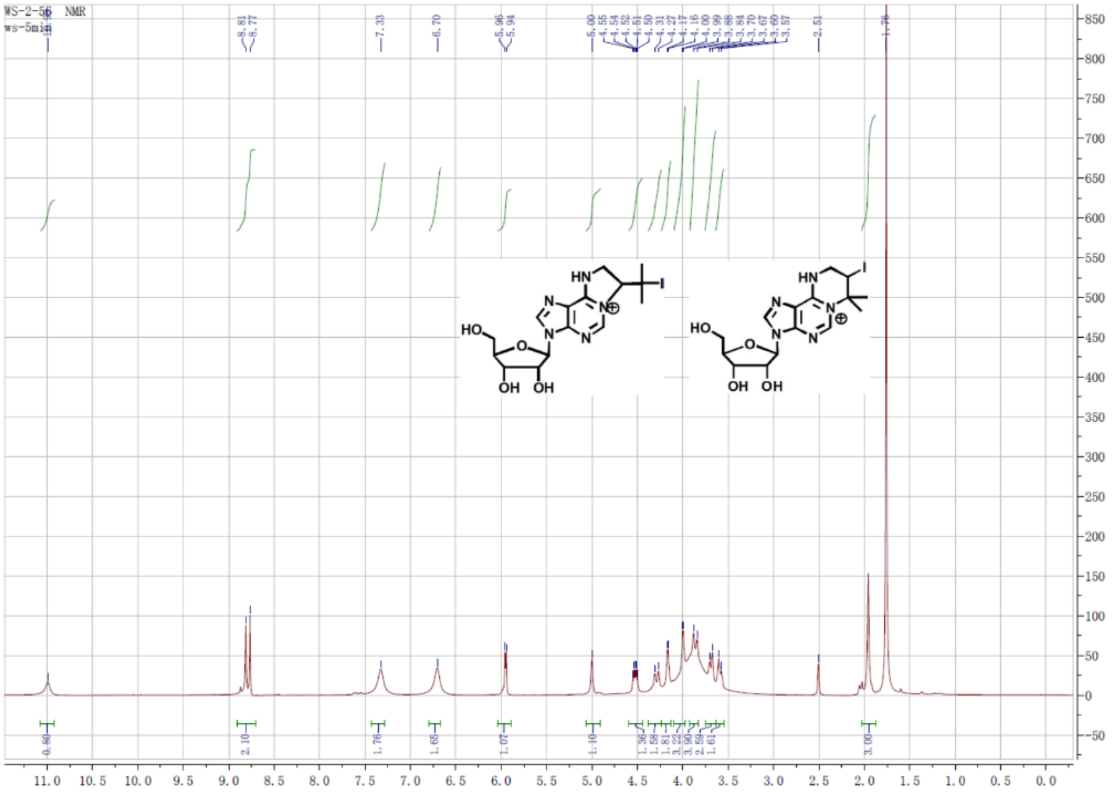

**Figure S33. $^{1}$H NMR** of crude product of i$^{6}$A (**4**) reacted with $I_2$ in DMSO-*d*$^{6}$.

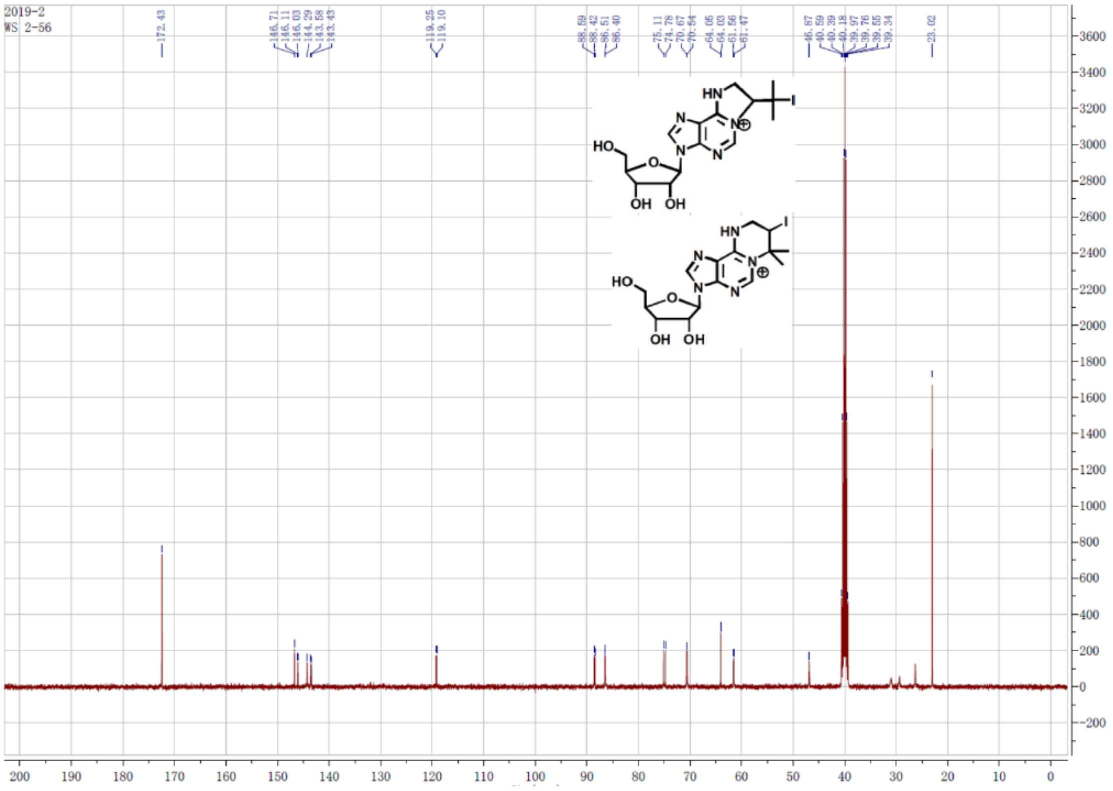

**Figure S34. $^{13}$C NMR** of crude product of i$^{6}$A (**4**) reacted with $I_2$ in DMSO-*d*$^{6}$.

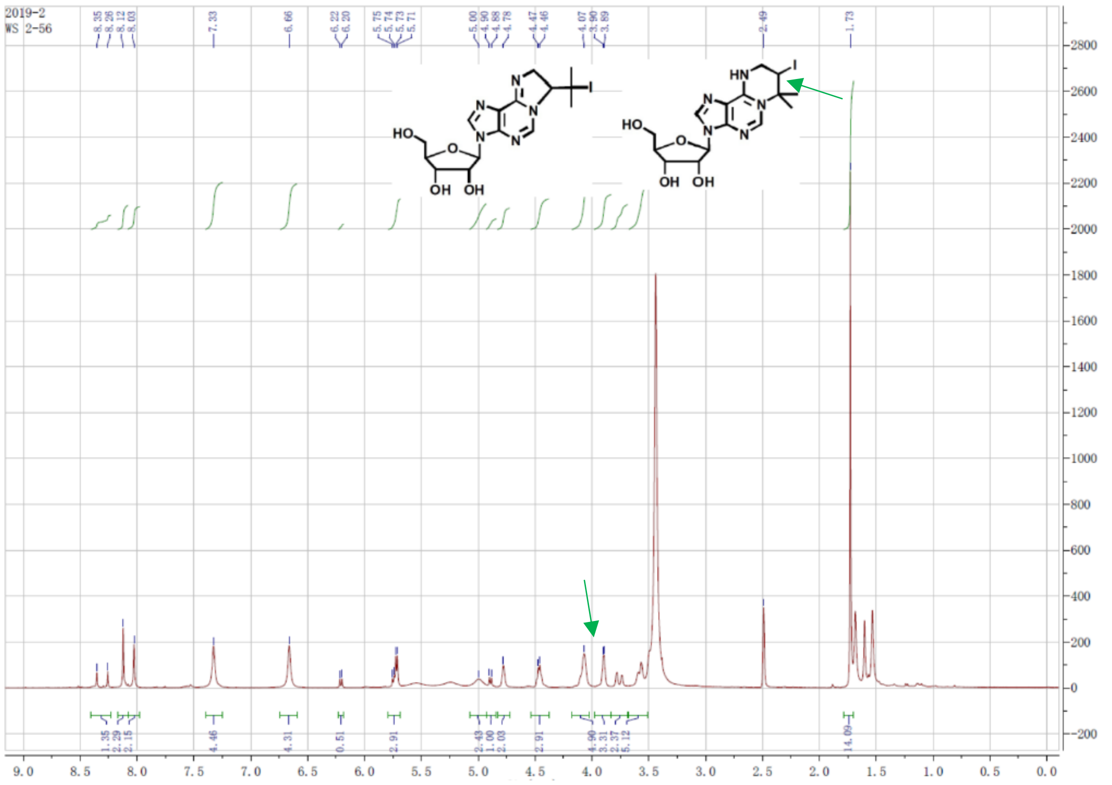

**Figure S35.** **$^{1}$H NMR** of crude product (**3a**/ **3b**) of i$^{6}$A (**4**) reacted with $I_2$ after $Na_2S_2O_3$ workup in DMSO-*d$^{6}$*.

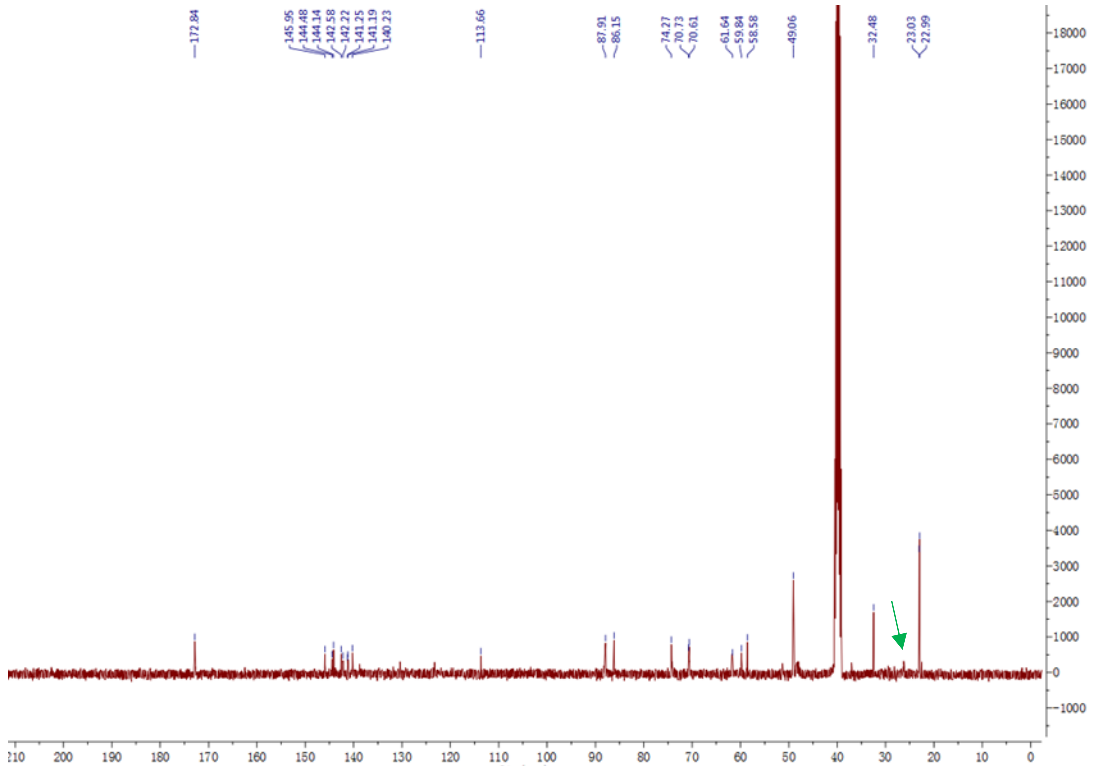

**Figure S36.** **$^{13}$C NMR** of crude product (**3a**/ **3b**) of i$^{6}$A (**4**) reacted with $I_2$ after $Na_2S_2O_3$ workup in DMSO-*d$^{6}$*.

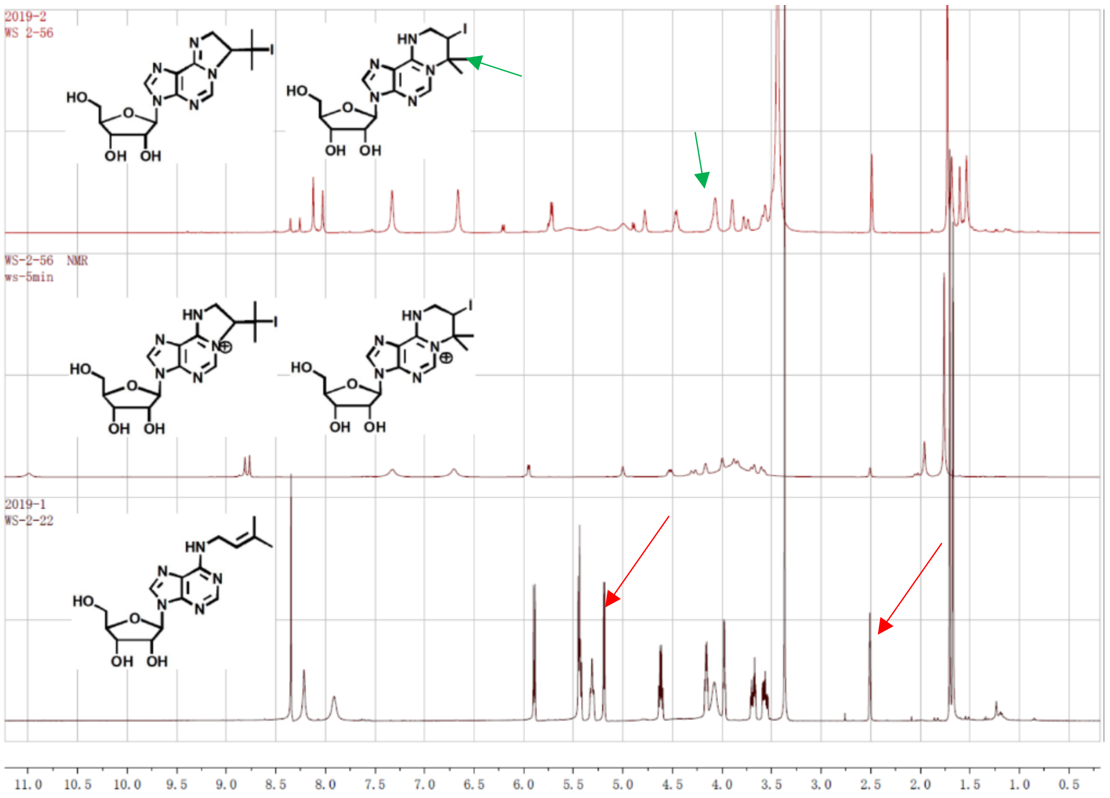

**Figure S37.** $^{1}$**H NMR** of compound **3a** and **3b** comparation of i$^{6}$A (**4**) reacted with $I_2$.

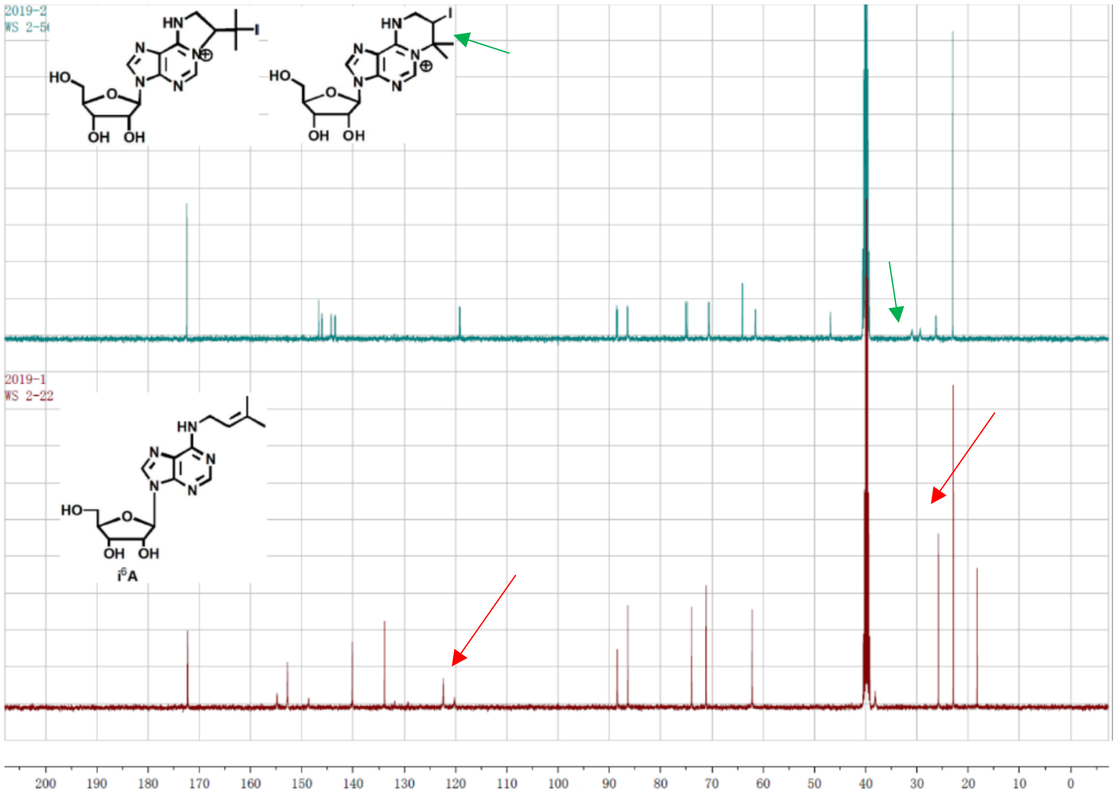

**Figure S38.** $^{13}$**C NMR** comparation of i$^{6}$A (**4**) reacted with $I_2$.

#### 9.3 UV spectra analysis.

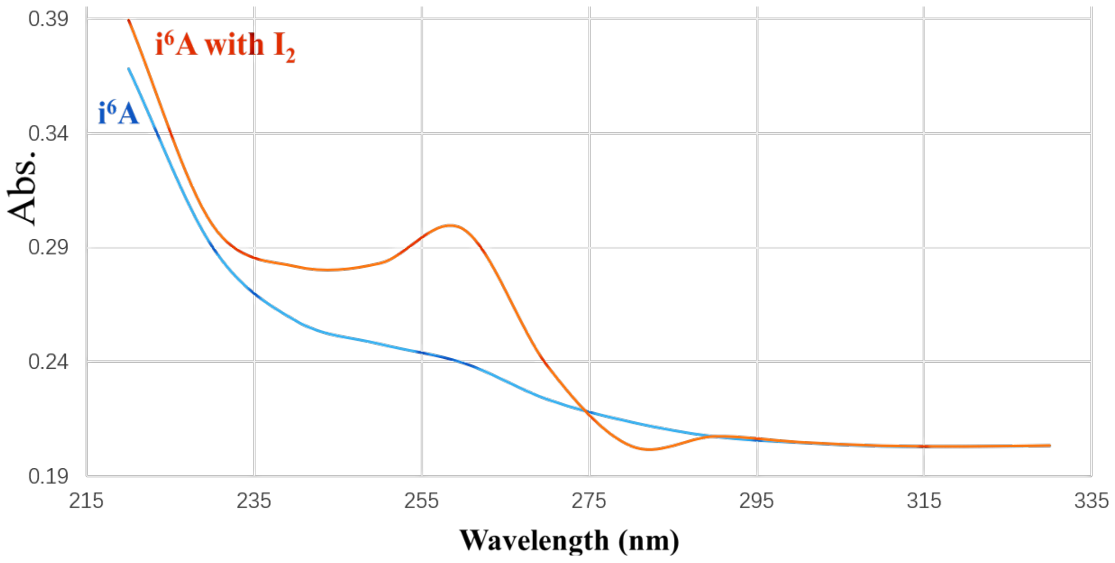

**Figure S39.** UV spectroscopy comparations of i$^6$A (**4**) with cyclo adduct (**3a/ 3b**). Condition: i$^6$A (**4**, 0.1 M) was dissolved in a mixture of acetonitrile and water (v/ v, 1: 1) and was analyzed by Ultraviolet Visible Spectrophotometer (*Perkin Elmer, Lambda 365*).

#### 9.4 DFT calculations of the reaction of i$^6$A (4) with iodine.

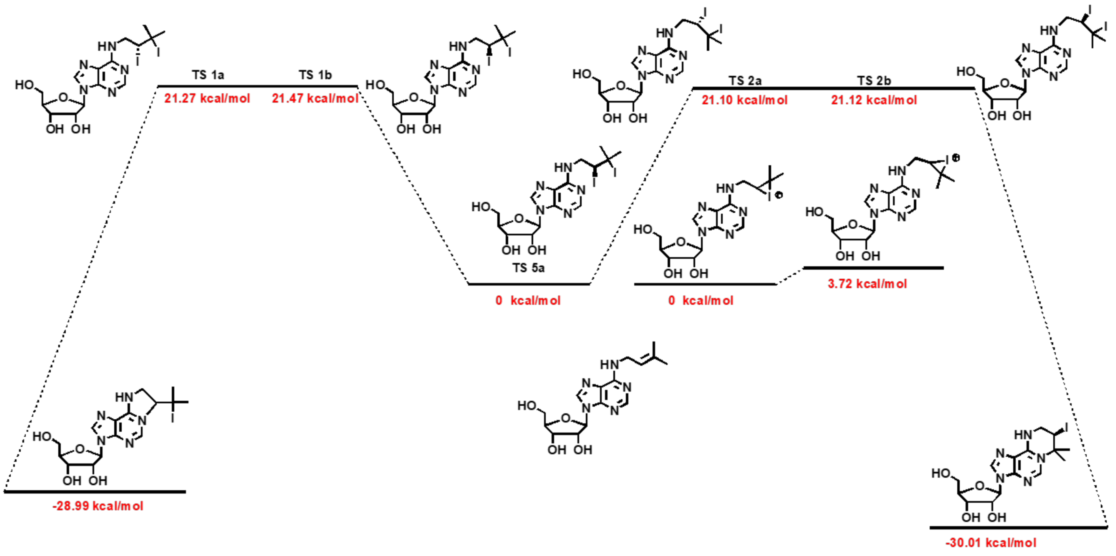

**Figure S40.** DFT calculations of the reaction of i$^6$A (**4**) with iodine. Deiodination process was calculated to be very fast. In contrast, the iodine epoxide intermediate was elusive.

### Computational Experiments

All the calculations were performed with the Gaussian 16, Revision B.01 program.

Geometry optimization of the model systems in the gas phase were carried out with the B3LYP/ BS1 DFT method augmented with the D3(BJ) version of Grimme's empirical dispersion correction (BS1 denotes a basis set combining SDD for I and 6-31G (d, p) for other atoms). Frequency calculations were performed at the same level of theory to determine whether the optimized structures are minima (no imaginary frequencies) or saddle points (one imaginary frequency) on the potential energy surface, and to provide thermal corrections to the Gibbs free energies. The solvation energy corrections were computed at the B3LYP/ BS1 level with the SMD solvation model for DMSO on gas-phase optimized geometries. The single point energies were computed with M062X/ BS2 augmented with the D3 version of Grimme's empirical dispersion correction (BS2 denotes a basis set combining SDD for I and Def2TZVP for other atoms).

Gaussian 16 Citation: Gaussian 16, Revision B.01, M. J. Frisch, G. W. Trucks, H. B. Schlegel, G. E. Scuseria, M. A. Robb, J. R. Cheeseman, G. Scalmani, V. Barone, G. A. Petersson, H. Nakatsuji, X. Li, M. Caricato, A. V. Marenich, J. Bloino, B. G. Janesko, R. Gomperts, B. Mennucci, H. P. Hratchian, J. V. Ortiz, A. F. Izmaylov, J. L. Sonnenberg, D. Williams-Young, F. Ding, F. Lipparini, F. Egidi, J. Goings, B. Peng, A. Petrone, T. Henderson, D. Ranasinghe, V. G. Zakrzewski, J. Gao, N. Rega, G. Zheng, W. Liang, M. Hada, M. Ehara, K. Toyota, R. Fukuda, J. Hasegawa, M. Ishida, T. Nakajima, Y. Honda, O. Kitao, H. Nakai, T. Vreven, K. Throssell, J. A. Montgomery, Jr., J. E. Peralta, F. Ogliaro, M. J. Bearpark, J. J. Heyd, E. N. Brothers, K. N. Kudin, V. N. Staroverov, T. A. Keith, R. Kobayashi, J. Normand, K. Raghavachari, A. P. Rendell, J. C. Burant, S. S. Iyengar, J. Tomasi, M. Cossi, J. M. Millam, M. Klene, C. Adamo, R. Cammi, J. W. Ochterski, R. L. Martin, K. Morokuma, O. Farkas, J. B. Foresman, and D. J. Fox, *Gaussian, Inc.,* Wallingford CT, **2016**.

### 10 Detection of $i^6$A modifications through mutant assay of $I_2$-mediated *EGFP* gene with $i^6$ATP.

General strategy: the structure of nucleoside $i^6$A (**4**) changes under the action of $I_2$. After the structural change, $i^6$A (**4**) no longer base-pairs with T or U. According to this principle, *c*DNA is obtained by reverse transcription of RNA containing $i^6$ATP, and the target gene sequence can be obtained by reverse transcription experiment using *c*DNA as a template. Then the target gene sequence is sequenced and compared with the natural sequence of the target gene. After in-depth analysis, the position information of the $i^6$A insertion site and the insertion efficiency can be identified.

#### 10.1 The efficiency of $i^6$ATP in T7 RNA polymerase transcription system was detected *in vitro*.

The *EGFP* mRNA sequence was obtained by transcribing under the conditions of $i^6$ATP: ATP ratio of 0: 1, 3: 7, 7: 3, 1: 0 through the *in vitro* transcription system, and then detected each group of *EGFP* RNA sequence through $I_2$ (concentration) treatment. After processing, the *c*DNA sequence of *EGFP* in each group was obtained by reverse transcription. Next, the obtained cDNA sequence was used as a template, and the mutant *EGFP* gene sequence in each group was obtained by PCR amplification, and cloned into the T vector. Through the amplification of DH5*α E. coli.,* 50 clones were picked from each group and sequenced to detect the mutation. After the mutation site of *EGFP* gene, and analyze the insertion efficiency of $i^6$ATP.

#### 10.2 Comparative analysis of the recognition efficiency of T7 RNA polymerase and RNA polymerase II for $i^6$ATP in eukaryotic cells.

*T7 RNA polymerase* and *T7-IRES2-EGFP* expression system were expressed in HeLa cells, and RNA was extracted 48 hours after transfection, and then the mixture was treated with $I_2$ to cause $i^6$ATP insertion site mutation. Then, the mutant sequences of the two endogenously expressed genes of *GAPDH* and *Notch1* were obtained by reverse transcription, and the mutant gene sequence of *EGFP* catalyzed by *T7 RNA polymerase* transcription was obtained. Same as the previous detection process, by detecting the insertion efficiency of $i^6$ATP in the endogenous highly expressed gene

*GAPDH* and the oncogene *Notch1* and the insertion efficiency of i$^6$ATP in *T7-EGFP*. Compare the recognition efficiency of i$^6$ATP by eukaryotic RNA polymerase and the feasibility of using it as a real-time transcriptome marker.

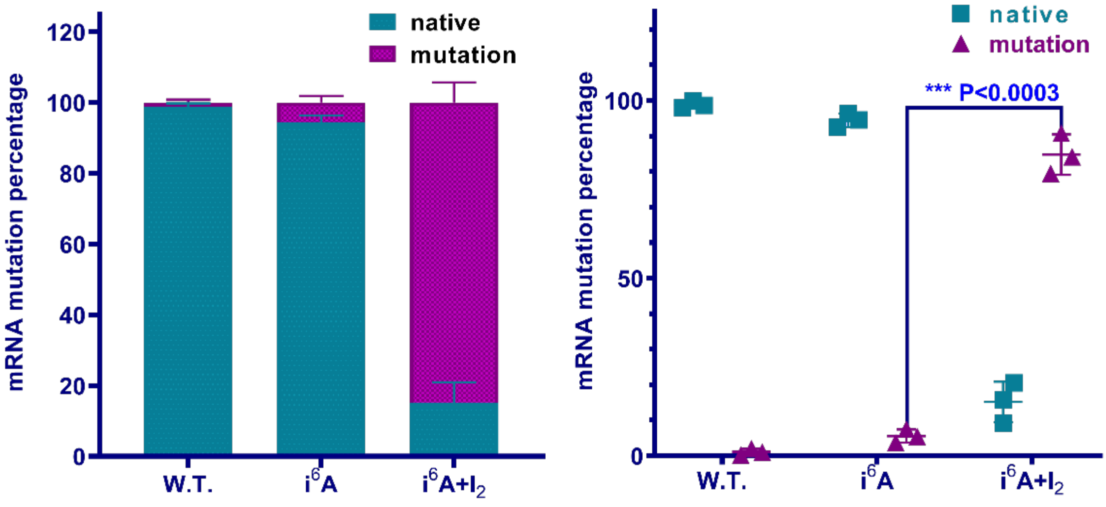

**Figure S41.** Results of mutant assays of i$^6$A (**4**)-incorporated RNA.

### 16. References.

(1) M. Habibian, X. Shu, Q. Dai, T. Wu, I. R. Bothwell, Y.-N. Yue, Z.-Z. Zhang, J. Cao, Q.-L. Fei, M.-K. Luo, C. He, and J.-Z. Liu. *N$^6$-allyladenosine: a new small molecule for RNA labeling identified by mutation assay*. *J. Am. Chem. Soc. 139*, 17213-17216 (**2017**).

(2) X. Shu, J. Cao, M.-H. Cheng, S.-Y. Xiang, M.-S. Gao, T. Li, X.-E. Ying, F.-Q. Wang, Y.-N. Yue, Z.-K. Lu, Q. Dai, X.-L. Cui, L.-J. Ma, Y.-Z. Wang, C. He, X.-H. Feng and J.-Z. Liu. *A metabolic labeling method detects m$^6$A transcriptome-wide at single base resolution*. *Nat. Chem. Bio. 16*, 887-895 (**2020**).

### Extended Data

#### 1. Analysis of RNA polymerase.

Sequences of T7 RNA polymerase.

```
         10         20         30         40         50
MNTINIAKND FSDIELAAIP FNTLADHYGE RLAREQLALE HESYEMGEAR
         60         70         80         90        100
FRKMFERQLK AGEVADNAAA KPLITTLLPK MIARINDWFE EVKAKRGKRP
        110        120        130        140        150
TAFQFLQEIK PEAVAYITIK TTLACLTSAD NTTVQAVASA IGRAIEDEAR
        160        170        180        190        200
FGRIRDLEAK HFKKNVEEQL NKRVGHVYKK AFMQVVEADM LSKGLLGGEA
        210        220        230        240        250
WSSWHKEDSI HVGVRCIEML IESTGMVSLH RQNAGVVGQD SETIELAPEY
        260        270        280        290        300
AEAIATRAGA LAGISPMFQP CVVPPKPWTG ITGGGYWANG RRPLALVRTH
        310        320        330        340        350
SKKALMRYED VYMPEVYKAI NIAQNTAWKI NKKVLAVANV ITKWKHCPVE
        360        370        380        390        400
DIPAIEREEL PMKPEDIDMN PEALTAWKRA AAAVYRKDKA RKSRRISLEF
        410        420        430        440        450
MLEQANKFAN HKAIWFPYNM DWRGRVYAVS MFNPQGNDMT KGLLTLAKGK
        460        470        480        490        500
PIGKEGYYWL KIHGANCAGV DKVPFPERIK FIEENHENIM ACAKSPLENT
        510        520        530        540        550
WWAEQDSPFC FLAFCFEYAG VQHHGLSYNC SLPLAFDGSC SGIQHFSAML
        560        570        580        590        600
RDEVGGRAVN LLPSETVQDI YGIVAKKVNE ILQADAINGT DNEVVTVTDE
        610        620        630        640        650
NTGEISEKVK LGTKALAGQW LAYGVTRSVT KRSVMTLAYG SKEFGFRQQV
        660        670        680        690        700
LEDTIQPAID SGKGLMFTQP NQAAGYMAKL IWESVSVTVV AAVEAMNWLK
        710        720        730        740        750
SAAKLLAAEV KDKKTGEILR KRCAVHWVTP DGFPVWQEYK KPIQTRLNLM
        760        770        780        790        800
FLGQFRLQPT INTNKDSEID AHKQESGIAP NFVHSQDGSH LRKTVVWAHE
        810        820        830        840        850
KYGIESFALI HDSFGTIPAD AANLFKAVRE TMVDTYESCD VLADFYDQFA
        860        870        880 883
DQLHESQLDK MPALPAKGNL NLRDILESDF AFA
```

Structural analysis of T7 RNA polymerase.

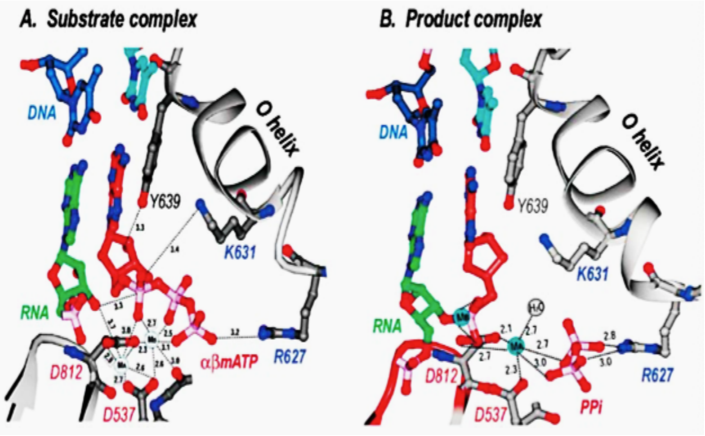

**Figure S42**. Crystal structure T7 RNA polymerase (PDB code: 1s76 and 1s77).

The crystal structure reported by Steinz *et al.* have indicated that the core AA→**Y639** play critical role in process of the recognition of substrate elongation into the nucleic acid sequence. Besides, **K631** and **R627** may also be important.

Crystal analysis of the T7 RNA polymerase.

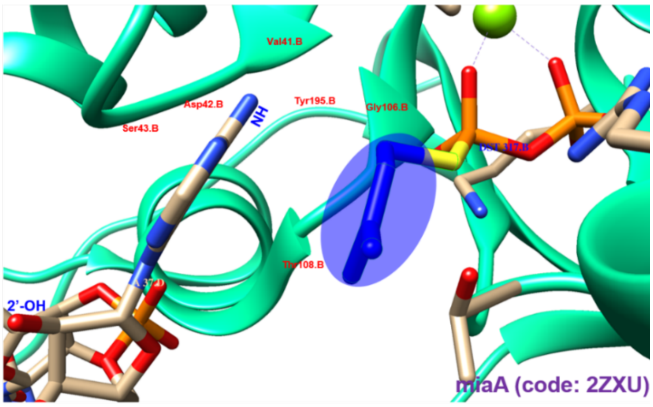

**Figure S43.** Crystal structure of MiaA.

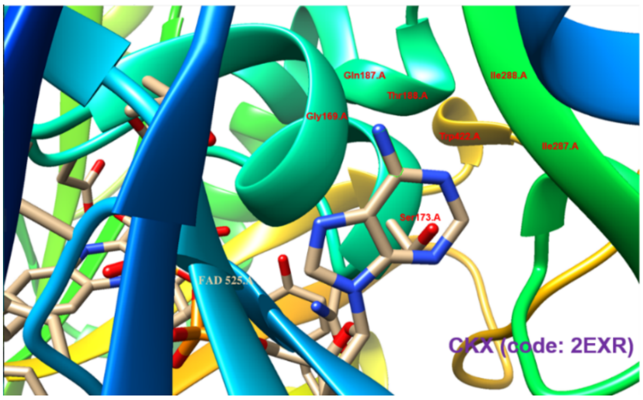

**Figure S44.** Crystal structure of CKX.

### 2. Original sequences data of templated EGFP gene.

```
Templated EGPF sequence used here
1          10          20          30          40          50
ACAATGACTG ATTACGATTC GAGCTCGGTA CCCGGGGATC CTCTAGAGAT
TTAATACGAC TCACTATAGG GATGGTGAGC AAGGGCGAGG AGCTGTTCAC100
CGGGGTGGTG CCCATCCTGG TCGAGCTGGA CGGCGACGTA AACGGCCACA
AGTTCAGCGT GTCCGGCGAG GGCGAGGGCG ATGCCACCTA CGGCAAGCTG200
ACCCTGAAGT TCATCTGCAC CACCGGCAAG CTGCCCGTGC CCTGGCCCAC
CCTCGTGACC ACCCTGACCT ACGGCGTGCA GTGCTTCAGC CGCTACCCCG300
ACCACATGAA GCAGCACGAC TTCTTCAAGT CCGCCATGCC CGAAGGCTAC
GTCCAGGAGC GCACCATCTT CTTCAAGGAC GACGGCAACT ACAAGACCCG400
CGCCGAGGTG AAGTTCGAGG GCGACACCCT GGTGAACCGC ATCGAGCTGA
AGGGCATCGA CCTCAAGGAG GACGGCAACA TGCTGGGGCA CAAGCTGGAG50[illegible]
TACAACTACA ACAGCCACAA CGTCTATATC ATGGCCGACA AGCAGAAGAA
CGGCATCAAG GTGAACTTCA AGATCCGCCA CAACATCGAG GACGGCAGCG600
TGCAGCTCGC CGACCACTAC CAGCAGAACA CCCCCATCGG CGACGGCCCC
GTGCTGCTGC CCGACAACCA CTACCTGAGC ACCCAGTCCG CCCTGAGCAA700
AGACCCCAAC GAGAAGCGCG ATCACATGGT CCTGCTGGAG TTCGTGACCG
CCGCCGGGAT CACTCTCGGC ATGGACGAGC TGTACAAGTA GAATCGTCGA800
CCTGCAGGCA TGCAAGCTTG GCACTGGCCG TCGTTTTACA ACGTCGTGAC
TGGGAAAACC CTGGCGTTAC CCAACTTAAT CGCCTTGCAG CACATCCCC899
```

### 3. Dynamic properties investigations of $i^6A$ (4) incorporated RNA by co-transcription and LC-MS quantification.

#### Cell culture.

HeLa cells (ATCC) were cultured in DMEM (Gibco) supplemented with 10% (v/ v) fetal bovine serum (Gibco), penicillin and streptomycin (Gibco), and grown at 37 °C with 5% $CO_2$. For metabolic labeling, HeLa cells at -80% confluence were treated with $i^6$ATP (500 µM).

#### $i^6$ATP co-transcription and enrichment.

$i^6$ATP was cultured for 5 minutes, 15 minutes, 30 minutes, 1 hour, 2 hours, 4 hours, and 16 hours, respectively. Cellular total RNAs were isolated using Trizol™ Reagent (*Invitrogen*) by following manufacturer's protocol as following.

***Step 1.*** Lyse and homogenize samples in Trizol™ reagent according to your starting material.

Cell grown in monolayer: Remove growth media. Add of Trizol™ reagent (1.0 mL) per 1 × $10^5$-$10^7$ cells directly to the culture dish to lyse the cells. Pipet the lysate up and down several times to homogenize. Incubate for 5 minutes to permit complete dissociation of the nucleoproteins complex. Add chloroform (0.2 mL) or of 4-bromoanisole (50 µL) per 1 mL of Trizol™. Reagent used for lysis, then securely cap the tube. Incubate for 2-3 minutes. Centrifuge the sample for 15 minutes at 12,000 r.p.m. at 4 °C. The mixture separates into a lower red phenol-chloroform, and interphase, and a colorless upper aqueous phase. Transfer ~600 µL of the colorless, upper aqueous phase containing the RNA to a new tube. Add an equal volume of 70% ethanol, then mix well by vertexing. Invert the tube to disperse any visible precipitate that may form after adding ethanol.

***Step 2.*** Bind the RNA to the membrane.

Transfer up to 700 µL of the sample to a spin cartridge (with collection tube). Centrifuge at 12,000 r.m.p. for 15 s. Discard the flow through then reinsert the spin cartridge into the same collection tube. Repeat until the entire sample has been processed.

***Step 3.*** Wash the RNA on the membrane.

Add 700 µL of Wash buffer I to the spin cartridge. Centrifuge at 12,000 r.p.m. for 15 s. Discard the flow through then reinsert the spin cartridge into the same collection tube. Add 500 µL of Wash buffer II to the spin cartridge. Centrifuge at 12,000 × g for 15 s. Discard the flow through then reinsert the spin cartridge into the same collection tube. Repeat once.

***Step 4.*** Elute the RNA.

Centrifuge at 12,000 r.p.m. for 1 minutes to dry the membrane. Discard the collection tube, then insert the spin cartridge into a recovery tube. Add 30 µL-3 × 100 µL (3 sequential elution with 100 µL each) of RNase-free water to the center of the spin cartridge. Note: If you are performing sequential elution, collect all eluates in the same tube. Incubate for 1 minute. Centrifuge at >12,000 × g for 2 minutes. Discard the spin cartridge. The recovery tube contains the purified total RNA. Store the purified RNA on ice-water bath if used within a few hours. For long-term storage, store the purified RNA at -80 °C.

**General procedure for $i^6$ATP co-transcription and enrichment.**

For metabolic labeling, HeLa cells at ~80% confluence were treated with $i^6$ATP (500 $\mu$M) for 5 minutes, 15 minutes, 30 minutes, 1 hour, 4 hours, and 16 hours, respectively. Cellular total RNAs were isolated using manufacturer's protocol. For each sample, around 350 ng total RNA was digested by using 7 U *nuclease P1* (*Wako*) in 2.0 µL reaction containing $NH_4OAc$ (20.0 mM) at 42 °C for 4 hours. Afterwards, 7.0 µL *FastAP*

Thermosensitive alkaline phosphatase and 16.1 µL 10 × FastAP buffer (*Thermo Scientific*) was added and the reaction was incubated at 37 °C for 4 hours.

The adenosine and i$^6$A (**4**) standard samples were injected into LC/ MS/ MS. The adenosine standard samples were quantified by using the nucleoside to base ion mass transitions of 268.2 to 136.1. The i$^6$A standard samples (**4**) was quantified by using the nucleoside to base ion mass transitions of 336.2 to 204.2. The adenosine and i$^6$A (**4**) standard samples were co-injected into LC/ MS/ MS.

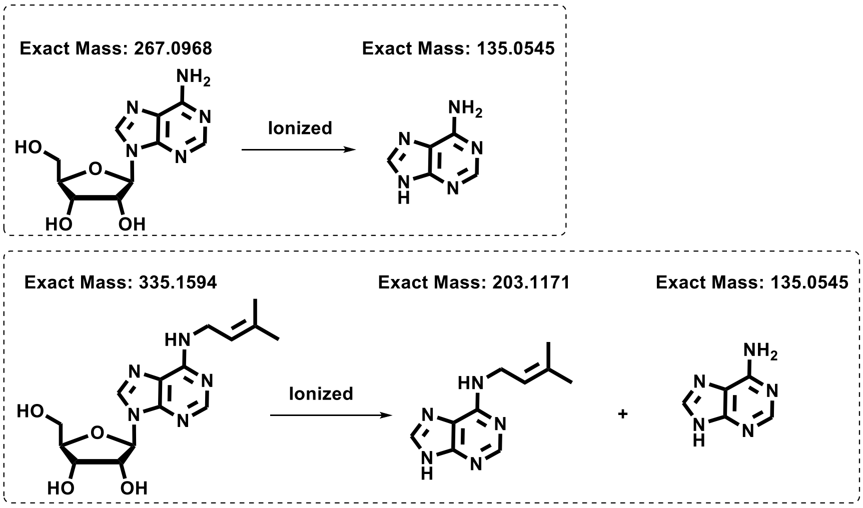

**Figure S45.** Analysis of the i$^6$A (**4**) ion fragement.

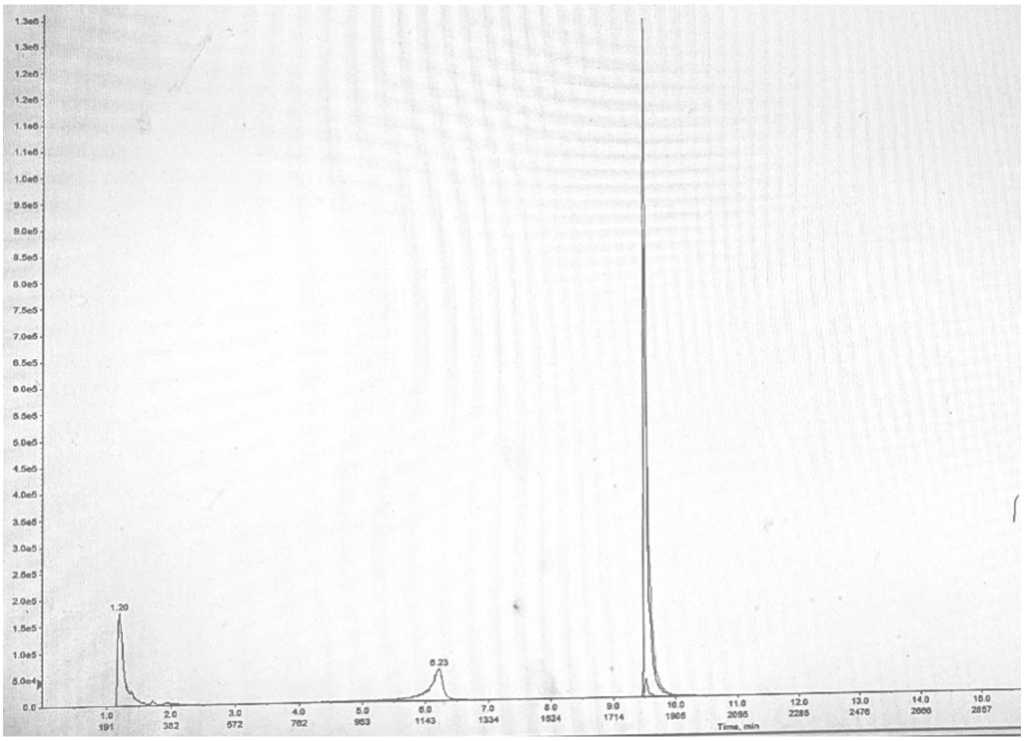

**Figure S46.** HPLC profile of the i$^6$A (**4**) ion fragments.

Samples were then filtered by 0.22 μm filter (*Millipore*) and diluted to 50 μL, and 10 μL was injected into LC/ MS/ MS.

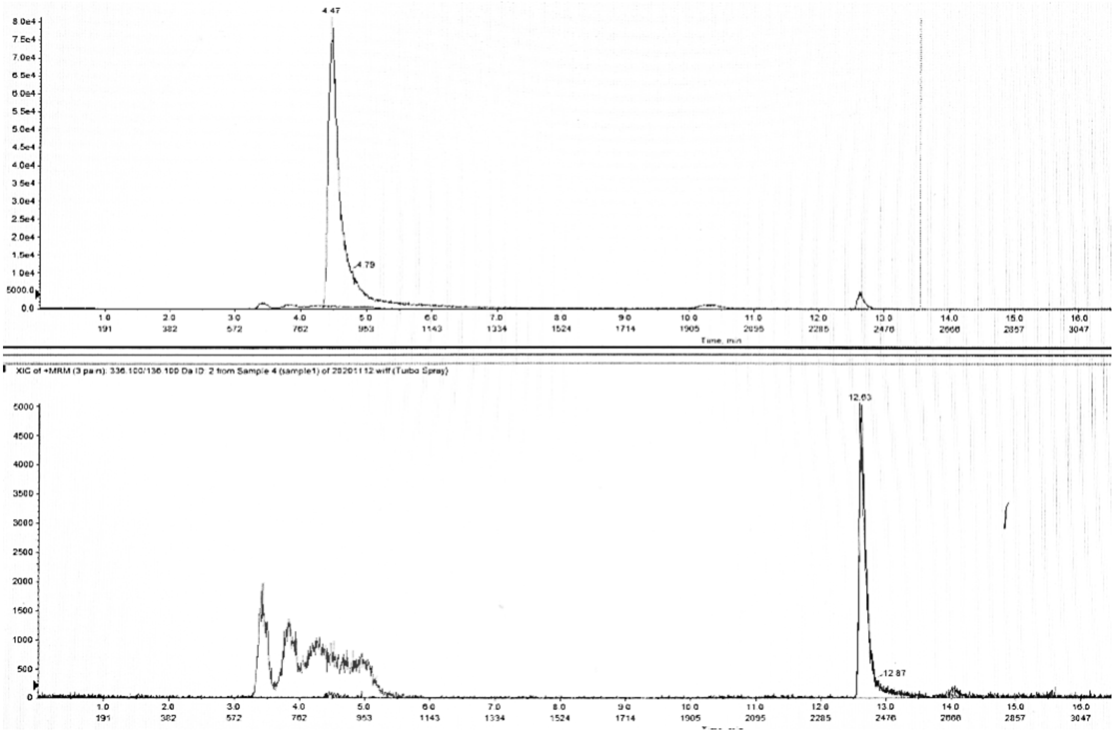

**Figure S47.** HPLC analysis of the i[6]A (**4**) ion fragments.

### Quantification of i[6]A (4) in RNA by LC/MS/MS.

Total RNAs were digested into nucleosides and the amount of i[6]A (**4**) was measured by using Agilent 6460 Triple Quad MS-MS with 1290 UHPLC supplied with a ZORBAX Eclipse XDB-$C_{18}$ column (UHPLCQQQ-MS/MS) and calculated based on the standard curve generated by pure standards. For each sample, around 50 ng RNA was digested by using 1 U *nuclease P1* (*Wako*) in 20.0 μL reaction containing $NH_4OAc$ (20 mM) at 42 °C for 2 hours. Afterwards, 1.0 μL *FastAP* Thermo-Sensitive Alkaline Phosphatase and 3.0 μL 10 × *FastAP* Buffer (*Thermo Scientific*) was added and the reaction was incubated at 37 °C for 4 hours. Samples were then filtered by 0.22 μm filter (*Millipore*) and diluted to 50.0 μL, and 10.0 μL was injected into LC/MS/MS.

The results are shown in Extended section.

### In-gel directly fluorescent labeling of prenylated RNA.

Total tRNA (from *Roche*, 1.0 mg) in PBS buffer (pH = 7.5) was treated with red fluorescent probe PTAD-DBCO-Cy5 (**10**, 1.0 mM, 1.0 $\mu$l) at room temperature for 10 minutes. The excess dye was removed through 3K filter (Millipore, Merck) and further were analyzed *via* Goldview dye staining or Cy5 fluorescent imaging. RNA was analyzed by gel electrophoresis on 1 % agarose. The electrophoresis conditions were as follows: power supply (DYY-6C power supply, Liuyi Biotechnology) was set to 130 V. 1% agarose gels were run at room temperature (25 °C) for 20 minutes and stained with or without GoodView™ (SBS Genentech, Beijing, China). A DNA size marker (1 kb DNA Ladder, TsingKe Biotech, TSJ102, Beijing, China) was used. Gels were analyzed by in-gel fluorescence measurements on a FluorChem® FC3 imager (Alpha Innotech). Fluorescence was measured with a blue light excitation wavelength (475 nm) and a green filter emission (537 nm).

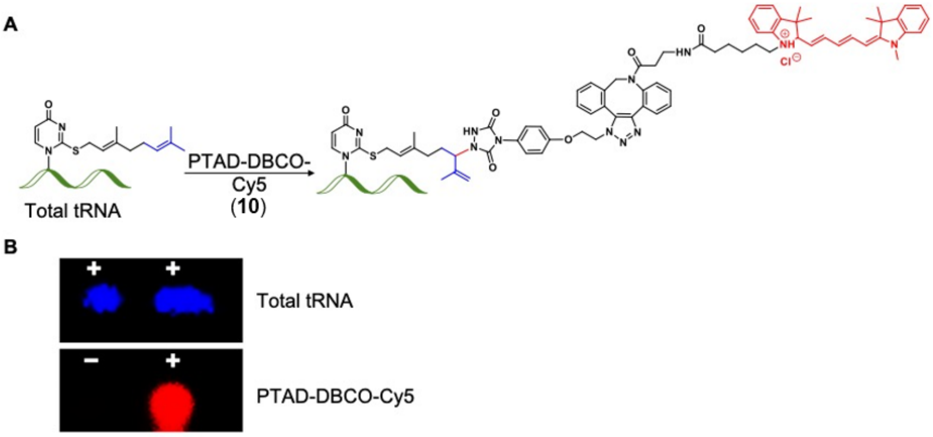

**Figure E1.** Directly fluorescent labeling of prenylated tRNA from *E. coli* using PTAD-DBCO-Cy5 (**10**).

### Quantification of nucleoside i$^6$A (4) in RNA by LC/MS/MS.

Total RNAs were digested into nucleosides and the amount of i$^6$A (**4**) was measured by using Agilent 6460 Triple Quad MS-MS with 1290 UHPLC supplied with a ZORBAX Eclipse XDB-$C_{18}$ column (UHPLCQQQ-MS/MS) and calculated based on the standard curve generated by pure standards. For each sample, around 50 ng RNA was digested by using 1 U *nuclease P1* (*Wako*) in 20.0 $\mu$L reaction containing $NH_4OAc$ (20 mM) at 42 $^o$C for 2 hours. Afterwards, 1.0 $\mu$L *FastAP* Thermo-Sensitive Alkaline Phosphatase and 3.0 $\mu$L 10 × FastAP Buffer (*Thermo Scientific*) was added and the reaction was incubated at 37 °C for 4 hours. Samples were then filtered by 0.22 $\mu$m filter (*Millipore*) and diluted to 50.0 $\mu$L, and 10.0 $\mu$L was injected into LC/MS/MS. The results are shown below.

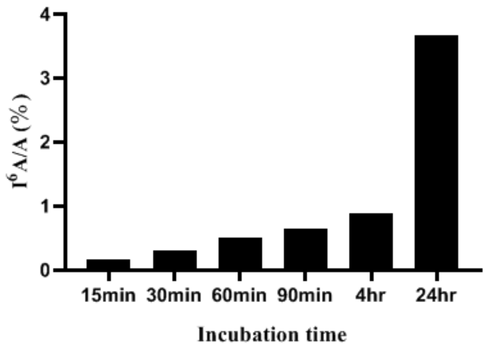

**Figure E2.** The i$^6$A (**4**) /A ratio increased with the incubation time. The co-transcription and qualification data indicated the increased gradually with the time. A represents adenosine herein.

### Table E1. Sequence information for T7 RNA polymerase promoter.

| T7 RNA polymerase promoter sequence |
|---|
| -15 -10 -5 +1 +5 |
| **TAATACGACTCACTATAGGG** |

### Table E2-3. Gene sequences used herein.

### Templated EGPF sequence used here

1 10 20 30 40 50

ACAATGACTG ATTACGATTC GAGCTCGGTA CCCGGGGATC CTCTAGAGAT
TTAATACGAC TCACTATAGG GATGGTGAGC AAGGGCGAGG AGCTGTTCAC$_{100}$
CGGGGTGGTG CCCATCCTGG TCGAGCTGGA CGGCGACGTA AACGGCCACA
AGTTCAGCGT GTCCGGCGAG GGCGAGGGCG ATGCCACCTA CGGCAAGCTG$_{200}$
ACCCTGAAGT TCATCTGCAC CACCGGCAAG CTGCCCGTGC CCTGGCCCAC
CCTCGTGACC ACCCTGACCT ACGGCGTGCA GTGCTTCAGC CGCTACCCCG$_{300}$
ACCACATGAA GCAGCACGAC TTCTTCAAGT CCGCCATGCC CGAAGGCTAC
GTCCAGGAGC GCACCATCTT CTTCAAGGAC GACGGCAACT ACAAGACCCG$_{400}$
CGCCGAGGTG AAGTTCGAGG GCGACACCCT GGTGAACCGC ATCGAGCTGA
AGGGCATCGA CCTCAAGGAG GACGGCAACA TGCTGGGGCA CAAGCTGGAG$_{500}$
TACAACTACA ACAGCCACAA CGTCTATATC ATGGCCGACA AGCAGAAGAA
CGGCATCAAG GTGAACTTCA AGATCCGCCA CAACATCGAG GACGGCAGCG$_{600}$
TGCAGCTCGC CGACCACTAC CAGCAGAACA CCCCCATCGG CGACGGCCCC
GTGCTGCTGC CCGACAACCA CTACCTGAGC ACCCAGTCCG CCCTGAGCAA$_{700}$
AGACCCCAAC GAGAAGCGCG ATCACATGGT CCTGCTGGAG TTCGTGACCG
CCGCCGGGAT CACTCTCGGC ATGGACGAGC TGTACAAGTA GAATCGTCGA$_{800}$
CCTGCAGGCA TGCAAGCTTG GCACTGGCCG TCGTTTTACA ACGTCGTGAC
TGGGAAAACC CTGGCGTTAC CCAACTTAAT CGCCTTGCAG CACATCCCC$_{899}$

### GAPDH  sequence

>NC_000012.12:6534517-6538371 Homo sapiens chromosome 12, GRCh38.p14 Primary Assembly

GCTCTCTGCTCCTCCTGTTCGACAGTCAGCCGCATCTTCTTTTGCGTCGCCAGGTGAAGACGGGCGGAGAGAAACCCGGGAGGCTAGGGACG
GCCTGAAGGCGGCAGGGGCGGGCGCAGGCCGGATGTGTTCGCGCCGCTGCGGGGTGGGCCCGGGCGGCCTCCGCATTGCAGGGGCGGGC
GGAGGACGTGATGCGGCGCGGGCTGGGCATGGAGGCCTGGTGGGGGAGGGGAGGGGAGGCGTGTGTGTCGGCCGGGGCCACTAGGCGC
TCACTGTTCTCTCCCTCCGCGCAGCCGAGCCACATCGCTCAGACACCATGGGGAAGGTGAAGGTCGGAGTCAACGGGTGAGTTCGCGGGTG
GCTGGGGGGCCCTGGGCTGCGACCGCCCCCGAACCGCGTCTACGAGCCTTGCGGGCTCCGGGTCTTTGCAGTCGTATGGGGGCAGGGTAGC
TGTTCCCCGCAAGGAGAGCTCAAGGTCAGCGCTCGGACCTGGCGGAGCCCCGCACCCAGGCTGTGGCGCCCTGTGCAGCTCCGCCCTTGCG
GCGCCATCTGCCCGGAGCCTCCTTCCCCTAGTCCCCAGAAACAGGAGGTCCCTACTCCCGCCCGAGATCCCGACCCGGACCCCTAGGTGGGG
GACGCTTTCTTTCCTTTCGCGCTCTGCGGGGTCACGTGTCGCAGAGGAGCCCCTCCCCCACGGCCTCCGGCACCGCAGGCCCCGGGATGCTA
GTGCGCAGCGGGTGCATCCCTGTCCGGATGCTGCGCCTGCGGTAGAGCGGCCGCCATGTTGCAACCGGGAAGGAAATGAATGGGCAGCCG
TTAGGAAAGCCTGCCGGTGACTAACCCTGCGCTCCTGCCTCGATGGGTGGAGTCGCGTGTGGCGGGGAAGTCAGGTGGAGCGAGGCTAGCT
GGCCCGATTTCTCCTCCGGGTGATGCTTTTCCTAGATTATTCTCTGGTAAATCAAAGAAGTGGGTTTATGGAGGTCCTCTTGTGTCCCCTCCCC
GCAGAGGTGTGGTGGCTGTGGCATGGTGCCAAGCCGGGAGAAGCTGAGTCATGGGTAGTTGGAAAAGGACATTTCCACCGCAAAATGGCC
CCTCTGGTGGTGGCCCCTTCCTGCAGCGCCGGCTCACCTCACGGCCCCGCCCTTCCCCTGCCAGCCTAGCGTTGACCCGACCCCAAAGGCCAG
GCTGTAAATGTCACCGGGAGGATTGGGTGTCTGGGCGCCTCGGGGAACCTGCCCTTCTCCCCATTCCGTCTTCCGGAAACCAGATCTCCCACC
GCACCCTGGTCTGAGGTTAAATATAGCTGCTGACCTTTCTGTAGCTGGGGGCCTGGGCTGGGGCTCTCTCCCATCCCTTCTCCCCACACACAT
GCACTTACCTGTGCTCCCACTCCTGATTTCTGGAAAAGAGCTAGGAAGGACAGGCAACTTGGCAAATCAAAGCCCTGGGACTAGGGGGTTAA
AATACAGCTTCCCCTCTTCCCACCCGCCCCAGTCTCTGTCCCTTTTGTAGGAGGGACTTAGAGAAGGGGTGGGCTTGCCCTGTCCAGTTAATTT
CTGACCTTTACTCCTGCCCTTTGAGTTTGATGATGCTGAGTGTACAAGCGTTTTCTCCCTAAAGGGTGCAGCTGAGCTAGGCAGCAGCAAGCA
TTCCTGGGGTGGCATAGTGGGGTGGTGAATACCATGTACAAAGCTTGTGCCCAGACTGTGGGTGGCAGTGCCCCACATGGCCGCTTCTCCTG
GAAGGGCTTCGTATGACTGGGGGTGTTGGGCAGCCCTGGAGCCTTCAGTTGCAGCCATGCCTTAAGCCAGGCCAGCCTGGCAGGGAAGCTC
AAGGGAGATAAAATTCAACCTCTTGGGCCCTCCTGGGGGTAAGGAGATGCTGCATTCGCCCTCTTAATGGGGAGGTGGCCTAGGGCTGCTC
ACATATTCTGGAGGAGCCTCCCCTCCTCATGCCTTCTTGCCTCTTGTCTCTTAGATTTGGTCGTATTGGGCGCCTGGTCACCAGGGCTGCTTTTA
ACTCTGGTAAAGTGGATATTGTTGCCATCAATGACCCCTTCATTGACCTCAACTACATGGTGAGTGCTACATGGTGAGCCCCAAAGCTGGTGT
GGGAGGAGCCACCTGGCTGATGGGCAGCCCCTTCATACCCTCACGTATTCCCCCAGGTTTACATGTTCCAATATGATTCCACCCATGGCAAAT
TCCATGGCACCGTCAAGGCTGAGAACGGGAAGCTTGTCATCAATGGAAATCCCATCACCATCTTCCAGGAGTGAGTGGAAGACAGAATGGA
AGAAATGTGCTTTGGGGAGGCAACTAGGATGGTGTGGCTCCCTTGGGTATATGGTAACCTTGTGTCCCTCAATATGGTCCTGTCCCCATCTCC
CCCCCACCCCCATAGGCGAGATCCCTCCAAAATCAAGTGGGGCGATGCTGGCGCTGAGTACGTCGTGGAGTCCACTGGCGTCTTCACCACCA
TGGAGAAGGCTGGGGTGAGTGCAGGAGGGCCCGCGGGAGGGGAAGCTGACTCAGCCCTGCAAAGGCAGGACCCGGGTTCATAACTGTCT
GCTTCTCTGCTGTAGGCTCATTTGCAGGGGGGAGCCAAAAGGGTCATCATCTCTGCCCCCTCTGCTGATGCCCCCATGTTCGTCATGGGTGTG
AACCATGAGAAGTATGACAACAGCCTCAAGATCATCAGGTGAGGAAGGCAGGGCCCGTGGAGAAGCGGCCAGCCTGGCACCCTATGGACA
CGCTCCCCTGACTTGCGCCCCGCTCCCTCTTTCTTTGCAGCAATGCCTCCTGCACCACCAACTGCTTAGCACCCCTGGCCAAGGTCATCCATGAC
AACTTTGGTATCGTGGAAGGACTCATGGTATGAGAGCTGGGGAATGGGACTGAGGCTCCCACCTTTCTCATCCAAGACTGGCTCCTCCCTGC
CGGGGCTGCGTGCAACCCTGGGGTTGGGGGTTCTGGGGACTGGCTTTCCCATAATTTCCTTTCAAGGTGGGGAGGGAGGTAGAGGGGTGAT
GTGGGGAGTACGCTGCAGGGCCTCACTCCTTTTGCAGACCACAGTCCATGCCATCACTGCCACCCAGAAGACTGTGGATGGCCCCTCCGGGA
AACTGTGGCGTGATGGCCGCGGGGCTCTCCAGAACATCATCCCTGCCTCTACTGGCGCTGCCAAGGCTGTGGGCAAGGTCATCCCTGAGCTG
AACGGGAAGCTCACTGGCATGGCCTTCCGTGTCCCCACTGCCAACGTGTCAGTGGTGGACCTGACCTGCCGTCTAGAAAAACCTGCCAAATA
TGATGACATCAAGAAGGTGGTGAAGCAGGCGTCGGAGGGCCCCCTCAAGGGCATCCTGGGCTACACTGAGCACCAGGTGGTCTCCTCTGAC
TTCAACAGCGACACCCACTCCTCCACCTTTGACGCTGGGGCTGGCATTGCCCTCAACGACCACTTTGTCAAGCTCATTTCCTGGTATGTGGCTG
GGGCCAGAGACTGGCTCTTAAAAAGTGCAGGGTCTGGCGCCCTCTGGTGGCTGGCTCAGAAAAAGGGCCCTGACAACTCTTTTCATCTTCTA
GGTATGACAACGAATTTGGCTACAGCAACAGGGTGGTGGACCTCATGGCCCACATGGCCTCCAAGGAGTAAGACCCCTGGACCACCAGCCC
CAGCAAGAGCACAAGAGGAAGAGAGAGACCCTCACTGCTGGGGAGTCCCTGCCACACTCAGTCCCCCACCACACTGAATCTCCCCTCCTCAC
AGTTGCCATGTAGACCCCTTGAAGAGGGGAGGGGCCTAGGGAGCCGCACCTTGTCATGTACCATCAATAAAGTACCCTGTGCTCAACCA

### Table E4. Mutant results analysis of i$^6$A-incorporated mRNA (templated EGPF)

1 10 20 30 40 50

**A**CAATGACTG ATTACGATTC GAGCTCGGTA CCCGGGGATC CTCTAGAGAT

TTAATACGAC TCACTATAGG GATGGTG**A**GC AAGGGCGAGG AGCTGTTCAC$_{100}$

CGGGGTGGTG CCCATCCTGG TCGAGCTGGA CGGCGACGTA AACGGCCACA

AGTTCAGCGT GTCCGGCGAG GGCGAGGGCG ATGCCACCTA CGGCAAGCTG$_{200}$

ACCCTGAAGT TC**A**TCTGCAC CACCGGCA**A**G CTGCCCGTGC CCTGGCCCAC

CCTCGTGACC ACCCTGACCT ACGGCGTGCA GTGCTTCAGC CGCTACCCCG$_{300}$

ACCACATGAA GCAGCACGAC TTCTTCAAGT CCGCCATGCC CGAAGGCTAC

GTCCAGGAGC GCACCATCTT CTTCAAGGAC GACGGCAACT ACAAGACCCG$_{400}$

CGCCGAGGTG AAGTTCGAGG GCGACACCCT GGTGAACCGC ATCGAGCTGA

AGGGCATCGA CCTCAAGGAG G**A**CGGCAACA TGCTGGGGCA CAAGCTGGAG$_{500}$

TACAACTACA ACAGCCACAA CGTCTATATC ATGGCCGACA AGCAGAAGAA

CGGCATCAAG GTGAACTTCA **A**GATCCGCCA CAACATCGAG GACGGCAGCG$_{600}$

TGCAGCTCGC CGACCACTAC C**A**GCAGAACA CCCCCATCGG CGACGGCCCC

GTGCTGCTGC CCGACAACCA CT**A**CCTGAGC ACCC**A**GTCCG CCCTGAGCAA$_{700}$

AGACCCCAAC GAGAAGCGCG ATCAC**A**TGGT CCTGCTGGAG TTCGTGACCG

CCGCCGGGAT C**A**CTCTCGGC ATGGACGAGC TGTACAAGTA GAATCGTCGA$_{800}$

CCTGCAGGCA TGCAAGCTTG GCACTGGCCG TCGTTTTACA ACGTCGTGAC

TGGGAAAACC CTGGCGTTAC CCAACTTAAT CGCCTTGCAG CACATCCCC$_{899}$

**Blue A** means i$^6$A insertion and mutation position: A1, A78, A213, A229, A472, A571, A622, A673, A685, A726, A762.
